# A Fluorogenic Green Merocyanine-Based Probe to Detect Heparanase-1 Activity

**DOI:** 10.1101/2024.02.25.581963

**Authors:** Zachary M. Rabinowitz, Zhishen Wang, Jun Liu, Yuzhao Zhang, Alberto Jimenez Ybargollin, Mayrav Saketkhou, Lina Cui

**Author notes:** These authors contributed equally to this work (Zachary M. Rabinowitz, Zhishen Wang).

## Abstract

Heparanase-1 (HPSE-1), an endo-β-D-glucuronidase, is an extracellular matrix (ECM) remodeling enzyme that degrades heparan sulfate (HS) chains of heparan sulfate proteoglycans (HSPGs). HPSE-1 functions to remodel the ECM and thereby disseminate cells, liberate HS-bound bioactive molecules, and release biologically active HS fragments. Being the only known enzyme for the cleavage of HS, HPSE-1 regulates a number of fundamental cellular processes including cell migration, cytokine regulation, angiogenesis, and wound healing. Overexpression of HPSE-1 has been discovered in most cancers, inflammatory diseases, viral infections, among others. As an emerging therapeutic target, the biological role of HPSE-1 remains to be explored but is hampered by a lack of research tools. To expand the chemical tool-kit of fluorogenic probes to interrogate HPSE-1 activity, we design and synthesized a fluorogenic green disaccharide-based HPSE-1 probe using our design strategy of tuning the electronic effect of the aryl aglycon. The novel probe exhibits a highly sensitive 278-fold fluorescence turn-on response in the presence of recombinant human HPSE-1, while emitting green light at 560 nm, enabling the fluorescence imaging of HPSE-1 activity in cells.

## INTRODUCTION

Heparanase-1 (HPSE-1), an endo-β-D-glucuronidase belonging to the glycoside hydrolase 79 (GH79) family, is an extracellular matrix remodeling enzyme.^1-3^ HPSE-1 stands as the sole enzyme known to cleave heparan sulfate (HS) chains of heparan sulfate proteoglycans (HSPGs).^4^ HSPGs consist of glycosaminoglycan (e.g., HS) chains covalently attached to a core protein, localized on the cell surface, in the extracellular matrix (ECM), and in the basement membrane.^5^ These HSPGs are ubiquitously expressed, major components of the ECM and play pivotal roles in maintaining the integrity of the ECM through interacting closely with other ECM components.^6^ Furthermore, due to the structural diversity and functional complexity of HS chains, a plethora of biologically active molecules such as cytokines, chemokines, growth factors, and enzymes are sequestered in the ECM via binding to the HS chains.^7^ Thus, HSPGs also control homeostasis by serving as a storage depot for biologically active molecules, which can be fundamental for physiological as well as pathological processes such as tumor progression, angiogenesis, and inflammation.^8^ HPSE-1 is first produced as an inactive pre-proenzyme, which is converted to a proenzyme in the endoplasmic reticulum.^9, 10^ The inactive proenzyme is then shuttled to the Golgi apparatus, secreted, and internalized where it is further processed to an active heterodimer consisting of the 8 kDa and 50 kDa subunits by cathepsin L in late endosomes or lysosomes.^11-15^ Active HPSE-1 is located primarily in acidic late endosome or lysosomes, where it is believed to play an essential housekeeping role in catabolic processing of internalized HSPGs.^16, 17^ HPSE-1 can be secreted to degrade HS chains extracellularly and its activity is controlled by pH of the microenvironment.^18^ HPSE-1 is most active under acidic conditions and is largely inactive at normal physiological pH.^19, 20^ Via degrading its substrate HS extracellularly, HPSE-1 contributes to the degradation of the ECM and basement membranes, cell migration, and the release of sequestered active biological molecules involved in various biological and pathological processes.^7, 21, 22^ Under normal physiological conditions, HPSE-1 expression is restricted to specific cell and tissue types, including immune cells, platelets, and the placenta.^1, 23-28^ Its physiological functions span wound healing, tissue repair, embryonic morphogenesis, leukocyte trafficking and inflammation, and hair growth.^29-33^ Overexpression of HPSE-1 has been observed in various human cancers, where it is hijacked to enhance tumor progression and promote angiogenesis and metastasis.^16^ Elevated levels of HPSE-1 has been found to be linked poor prognosis of several cancers.^34-38^ Moreover, HPSE-1 plays significant roles in inflammatory diseases^39, 40^ and viral infections^41-50^.

Despite its pathological significance, the biological functions of HPSE-1 remain to be explored under various conditions. Traditional methods utilizing radiolabeled^51-56^ or fluorophore-labeled HS polysaccharides^57, 58^ present limitations for broad applications, due to the heterogeneous nature of HS polysaccharides, concerns about reproducibility, and restriction to *in vitro* samples. Therefore, there is pressing need for robust and feasible tools to interrogate HPSE-1 activity. A significant advancement in this regard involves the development of a new class of tools to detect HPSE-1 activity with high sensitivity that can be applied in various contexts. The concept and design of small molecule-based HPSE-1 probes capitalized on the elucidation of the minimum recognition unit of HPSE-1, a GlcN-GlcUA-GlcN trisaccharide^59^. HPSE-1 is an endo-β-D-glucuronidase, which catalyzes the hydrolysis of the internal glycosidic bond between the GlcN and GlcUA residues using its two catalytic residues Glu225 and Glu343 via a retaining catalytic mechanism.^3^ Therefore, the conventional approach of creating fluorogenic probes targeting hydrolases where a fluorophore is linked to the reducing end is ineffective when dealing with endo-acting carbohydrate-active enzymes.^60, 61^ Furthermore, the omission of the Glc residue on the reducing end of the trisaccharide sequence diminishes significant interactions between the substrate and the active site of HPSE-1, potentially compromising the probe’s activity.^62^ By tuning the electronic properties of the aryl aglycone moiety, we were able to twist the mode of action of HPSE-1 to allow for exo-glycosidic activities. This strategy was proved effective in the development of HADP, the first ultrasensitive small molecule HPSE-1-activatable probe, which demonstrated excellent selectivity and sensitivity in detecting human HPSE-1.^63^ Using this strategy, we also aim to expand the small-molecule probe tool-kit for HPSE-1 detection by developing other novel fluorogenic disaccharide-based probes with various fluorophores bearing different imaging characteristics (**Figure 1**). Fluorescent activity-based molecular imaging probes constructed using merocyanine-based dyes, which are characterized by a conjugated π-electron system with electron-donor and acceptor substituents, offer versatile colorimetric and fluorescent changes (e.g., “OFF-to-ON”) upon uncaging of their phenolic oxygen and restoring the intramolecular charge transfer (ICT) effect.^64-68^ Furthermore, their applicability in one- and two-photon fluorescence imaging and low cytotoxicity make them suitable for cell or *in vivo* imaging.^66, 67^ Herein, we describe our strategies to develop a novel green fluorogenic merocyanine-based HPSE-1 probe that can be used for cell imaging.

**Figure 1.**
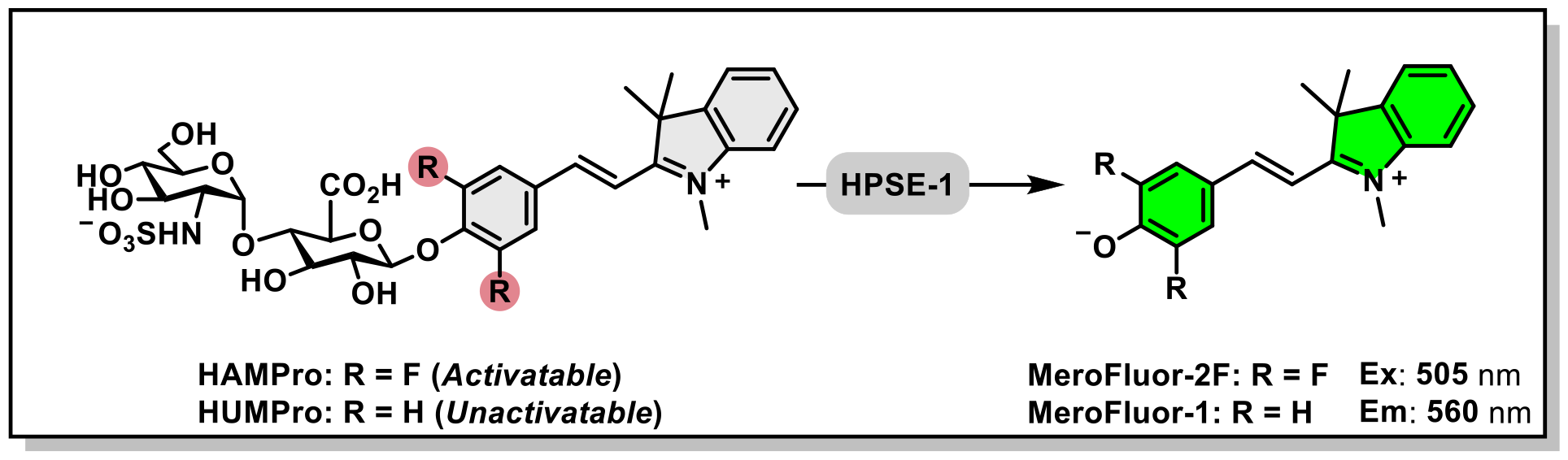
Overview of work presented herein.

## RESULTS and DISCUSSION

To expand the chemical tool-kit of HPSE-1 activity detecting probes, our goal was to design, synthesize, and evaluate a novel fluorogenic probe to detect HPSE-1 activity with green fluorescence emission using our previously successful design strategy^63^ (**Figure 1**). The merocyanine dye, (*E*)-2-(4-hydroxystyrl)-1,3,3-trimethyl-*3H*-indolium (**MeroFluor-1**), was chosen to construct a green fluorogenic HPSE-1 probe due to its excellent spectroscopic properties and water-solubility. Based on previous work, the installation of electron withdrawing groups (e.g., fluorine atoms) flanking the phenolic oxygen on the aryl aglycone is critical to permit HPSE-1 to display *exo*-glycosidase activity against the probe and free the fluorogenic reporter moiety.^63^

Therefore, the fluorophore must also present feasible synthetic manipulation of the phenolic aromatic ring to generate an activatable HPSE-1 probe. Indeed, **MeroFluor-1** analogs bearing electron withdrawing groups flanking the phenolic oxygen can be easily synthesized in one step from commercially available 4-hydroxybenzaldehyde derivatives. We designed **7** to contain the (*E*)-2-(3,5-difluoro-4-hydroxystyryl)-1,3,3-trimethyl-3H-indol-1-ium (**MeroFluor-2F**) fluorophore linked to the reducing end of the disaccharide, GlcN(NS)-GlcUA.^63^ Molecule **7** contains the proper electronic tuning of the aryl aglycone for HPSE-1 activation and thus, is referred to as <u>H</u>PSE-1 <u>A</u>ctivatable <u>M</u>erocyanine-based <u>Pro</u>be (**HAMPro**). Intact **HAMPro** would be non-fluorescent due to the caging of the phenolic oxygen by linkage to the dissacharide. In the presence of HPSE-1, **HAMPro** will be hydrolyzed unleashing **MeroFluor-2F**, which would generate a fluorogenic response. To confirm our previously described design strategy, we also designed probe **6** or <u>H</u>PSE-1 <u>U</u>nactivatable <u>M</u>erocyanine-based <u>Pro</u>be (**HUMPro**) to contain **MeroFluor-1** as the fluorophore, which bears no electronic tuning of the aryl aglycone.

We chemically synthesized **HUMPro** and **HAMPro**, according to the reaction route as outlined in **Figure 2**. We first embarked on the synthesis of these molecules from the previously reported compound **1**.^63^ Transformation of **1** into the corresponding glycosyl bromide intermediate in the presence of iodine bromide (IBr) allowed for further coupling of 4-hydroxybenzaldehyde or 3,5-difluoro-4-hydroxybenzaldehyde using silver oxide (Ag_2_O) as the activating agent in anhydrous acetonitrile to provide **2** and **3**, respectively. Then, global deacetylation, saponification, and reduction of the azido group to the amine afforded the deprotected GlcN-GlcUA dissacharide. Followed by, aldol condensation of the reactive aromatic aldehyde with 1,2,3,3-tetramethyl-*3H*-indolium iodide in the presence of triethylamine in ethanol under reflux provided **4** and **5** with the corresponding intact merocyanine-based fluorophore. It is worthwhile to note that caged **MeroFluor-1** and **MeroFluor-2F** are not stable under basic saponification conditions, due to a significant retro-aldol reaction side reaction. Therefore, with all protecting groups deprotected, late-stage installation of merocyanine-based fluorophores is necessary, which can be used to provide a library of fluorogenic small molecule-based HPSE-1 probes. Lastly, N-sulfation of the secondary amine of **4** and **5** using pyridine sulfur trioxide complex (PySO_3_) in the presence of triethylamine in acetonitrile provided **HUMPro** and **HAMPro** respectively, in excellent purity.

**Figure 2.**
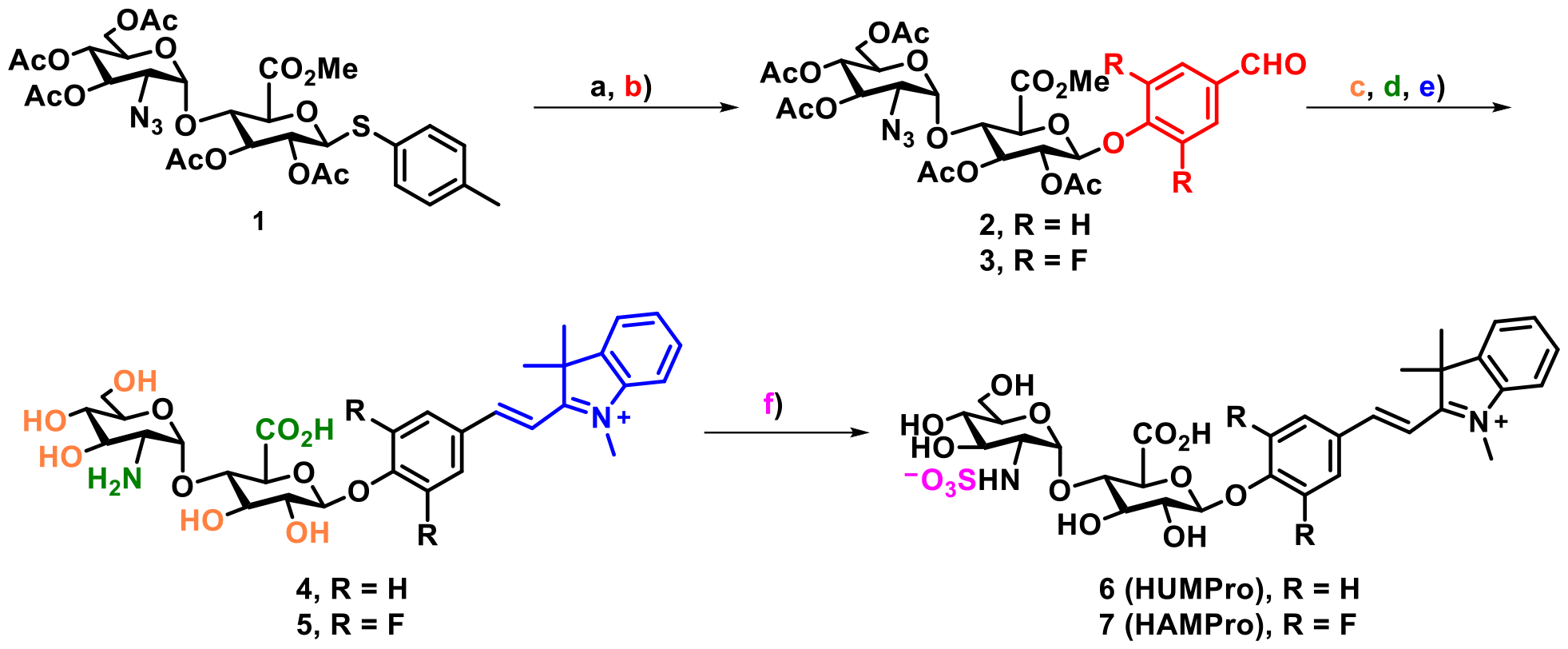
Synthetic Route to **6** (**HUMPro**) and **7** (**HAMPro**). Reagents and conditions: a) IBr, dry DCM, RT; b) 4-hydroxybenzaldehyde (R = H) or 3,5-difluoro-4-hydroxybenzaldehyde (R = F), Ag_2_O, dry MeCN, RT; c) NaOMe (25 wt% in MeOH), MeOH, RT; d) NaOH (1M) then PMe_3_ (1M in THF), Britton-Robinson buffer (pH 10), e) 1,2,3,3-tetramethyl-3*H*-indolium iodide, TEA, EtOH, reflux; f) PySO_3_, TEA, MeCN, RT.

When intact, these compounds are non-fluorescent (‘OFF’). However, in the presence of HPSE-1, the enzyme-catalyzed hydrolysis of the glycosidic linkage results in the dissociation of the disaccharide and the corresponding merocyanine-based fluorophore, resulting in increased fluorescence emission (‘OFF’ to ‘ON’) due to restored ICT (**Figure 3A**). The fluorescence emission of ICT-based merocyanine-based dyes is pH-dependent. Under the acidic (pH 5.0) enzymatic buffer condition, only **MeroFluor-2F** (pKa, ∼4) would have optimal fluorescence emission since the phenolate would dominate. Therefore, under the ideal HPSE-1 activity assay pH of 5.0, **HAMPro** is intrinsically well-suited for a simple one-step ‘mix-and-go’ assay. On the other hand, **MeroFluor-1** (pKa, ∼7) would result in a minimal fluorescence enhancement under the same acidic condition and would require an extra basification step to optimize the assay for **HUMPro**, if it were activated by HPSE-1 at all. With the probes in-hand, we first tested the stability of **HUMPro** and **HAMPro** towards HPSE-1 in 40 mM NaOAc buffer (pH 5.0) at 37°C by tracking the fluorescence emission over time. **HUMPro** is exceptionally stable towards solvolysis, but **HAMPro** was found to be non-enzymatically hydrolyzed after 7 hours under the acidic buffer condition (**Figure S1**). We, therefore, measured the increase in fluorescence emission over 6 hours to track the fluorogenic reaction, while avoiding non-enzymatic hydrolysis of **HUMPro** and **HAMPro**. For **HUMPro**, no increase in fluorescence was observed after incubation for 6 hours, while **HAMPro** immediately exhibited a fluorescence enhancement as early as 5 minutes and proceeded to increase over the 6-hour period (**Figure 3B**). After 6 hours of reaction, the fluorescence emission spectra were measured for all conditions (**Figure 3C**) and the turn-on ratio of **HAMPro** (i.e., the fluorescence intensity of **HAMPro** with HPSE-1 divided by without HPSE-1) was determined to be 278-fold (**Figure 3D**), suggesting that **HAMPro** is a highly sensitive probe. To validate the products of **HUMPro** or **HAMPro** in the presence of HPSE-1, we monitored the reaction products immediately after 6-hour reaction by HPLC. The reaction of **HAMPro** with HPSE-1 to yield **MeroFluor-2F** was supported by HPLC, giving 64% conversion to a new peak in alignment with that of **MeroFluor-2F**. This reaction can also be observed by eye through a color-change of the solution from clear to light-pink (**Figure S2**). Full conversion of **HAMPro** to **MeroFluor-2F** in the presence of HPSE-1 was not able to be achieved in our hands. Similar conversion results were obtained when we incubated **HAMPro** with 3-fold less of HPSE-1 (final concentration, 0.05 μg/μL), suggesting that inhibition of HPSE-1 activity by **HAMPro** or **MeroFluor-2F** may not contribute to this phenomenon (data not shown). Presumably, during the course of the reaction, recombinant human HPSE-1 is not stable and thus, the activity diminishes resulting in partial conversion. In contrast, the HPLC trace of **HUMPro** remained unchanged in the presence of HPSE-1 (**Figure 3F**), suggesting that our previously described design strategy is consistent.

**Figure 3.**
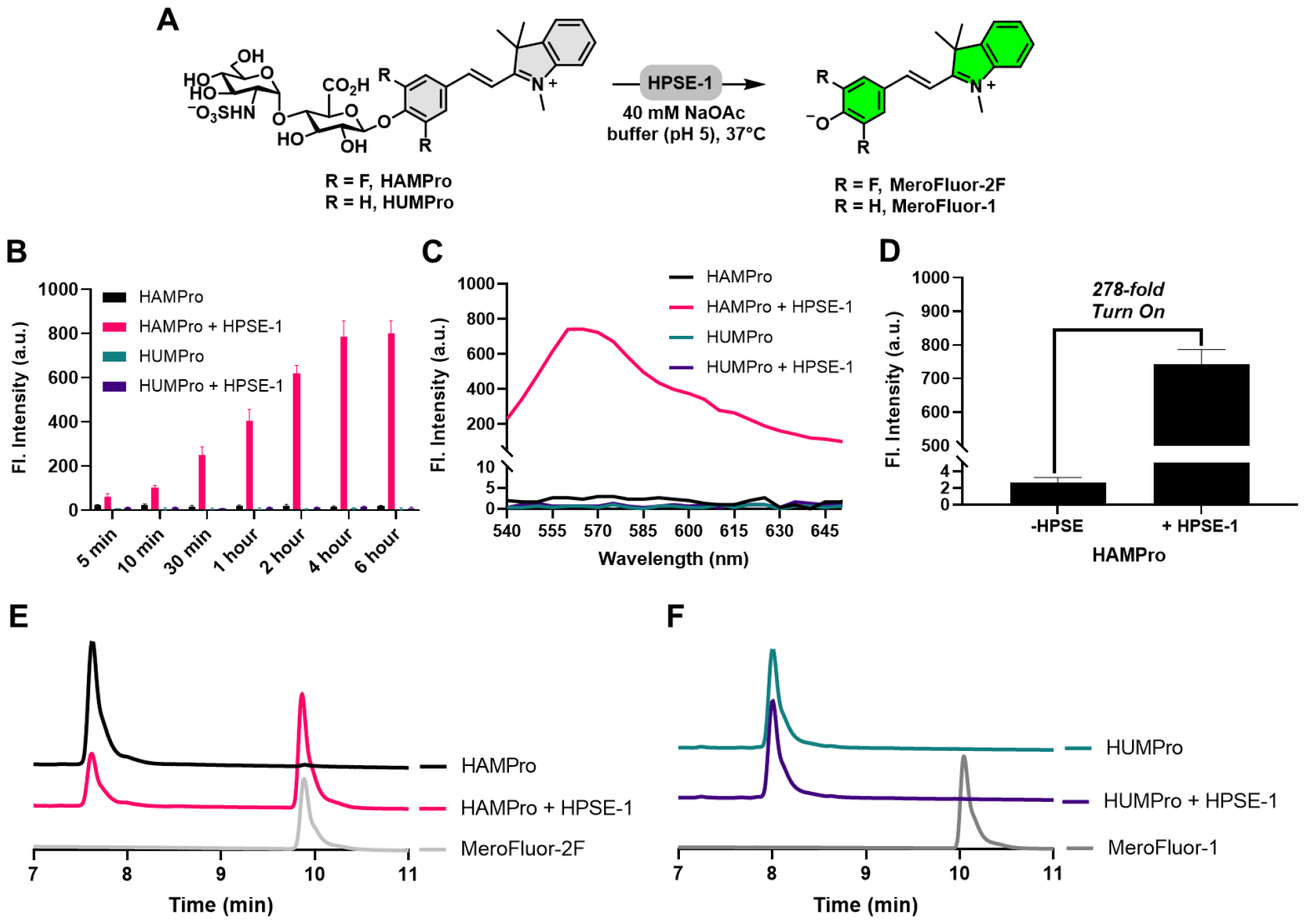
HAMPro and HUMPro Response to HPSE-1. (**A**) Plausible reaction of **HAMPro** or **HUMPro** with HPSE-1 to free the merocyanine-based dyes and thus, generate a fluorogenic response. (**B**) Fluorescence time-course response of **HAMPro** and **HUMPro** (Final concentration, 5 μM) without or with recombinant human HPSE-1 (Final concentration, 0.15 μg/μL) in 40 mM NaOAc buffer (pH 5.0) at 37°C (λ_ex_**/**λ_em_, 505/560 nm) for 6 hours. After 6 hours, the (**C**) fluorescence emission spectra were measured (λ_ex_**/**λ_em_, 505/540-560 nm) and the (**D**) turn-on ratio of **HAMPro** (λ_ex_**/**λ_em_, 505/560 nm) was determined from (**C**). After 6 hours, HPLC traces of the reaction of (**E**) **HAMPro** and (**F**) **HUMPro** with or without HPSE-1 (Observed Wavelength, 400 nm). **MeroFluor-2F and MeroFluor-1** standards are included for comparison.

We conclude that tuning the electronics of the aryl aglycone and thus, altering the strength of glycosidic bond is necessary for probe activation by HPSE-1 and can be expanded to other fluorophores like merocyanine-based dyes.

We next sought to determine if the modification of the merocyanine dye, **MeroFluor-1**, with fluorine atoms flanking to phenolic oxygen affected the fluorescence properties. To mimic caging (non-fluorescent, ‘OFF’) and uncaging (fluorescent, ‘ON’) of the phenolic oxygen, we incubated **MeroFluor-2F** and **MeroFluor-1** in buffers of various pH values (pH,1.5-10.8) to observe the pH-dependent ICT fluorescence mechanism and compare the turn-on ratios (**Figure 4A, B**). We observed that **MeroFluor-2F** has a 2-fold fluorescence turn-on response, whereas **MeroFluor-1** has a 14-fold fluorescence turn-on response. Due to a dramatic increase in the fluorescence background in the protonated state (‘OFF’), **MeroFluor-2F** resulted in a poor turn-on ratio. This suggests that such modification with fluorine atoms impacted the fluorescence properties of this fluorophore. In contrast, compared to our previous study with HADP, we observed that the fluorescence was not deleteriously impacted after modifying the 4MU scaffold with fluorine atoms flanking the phenolic oxygen (**Figure S3**). In fact, the fluorescence properties of 4MU were improved upon modification, when compared to DiFMU. This proposes that modification of fluorophores based on the ICT mechanism with electron-withdrawing groups flanking the phenolic oxygen can be either deleterious or beneficial for sensitive detection of an analyte, depending on the structure and nature of the fluorophore. In the case where it is deleterious, it presents a dilemma for development of fluorogenic HPSE-1 probes using the design strategy described herein. However, it is important to note that even though **HAMPro** did not fully react with HPSE-1 (only 64% conversion), it still displayed a highly sensitive 278-fold turn-on ratio. Therefore, caging of such modified fluorophores with a dissacharide, where it is linked to a carbon atom instead of a hydrogen atom, may be able to fully suppress the high background fluorescence intensity observed for **MeroFluor-2F** in its fully protonated form (**Figure 4E**).

**Figure 4.**
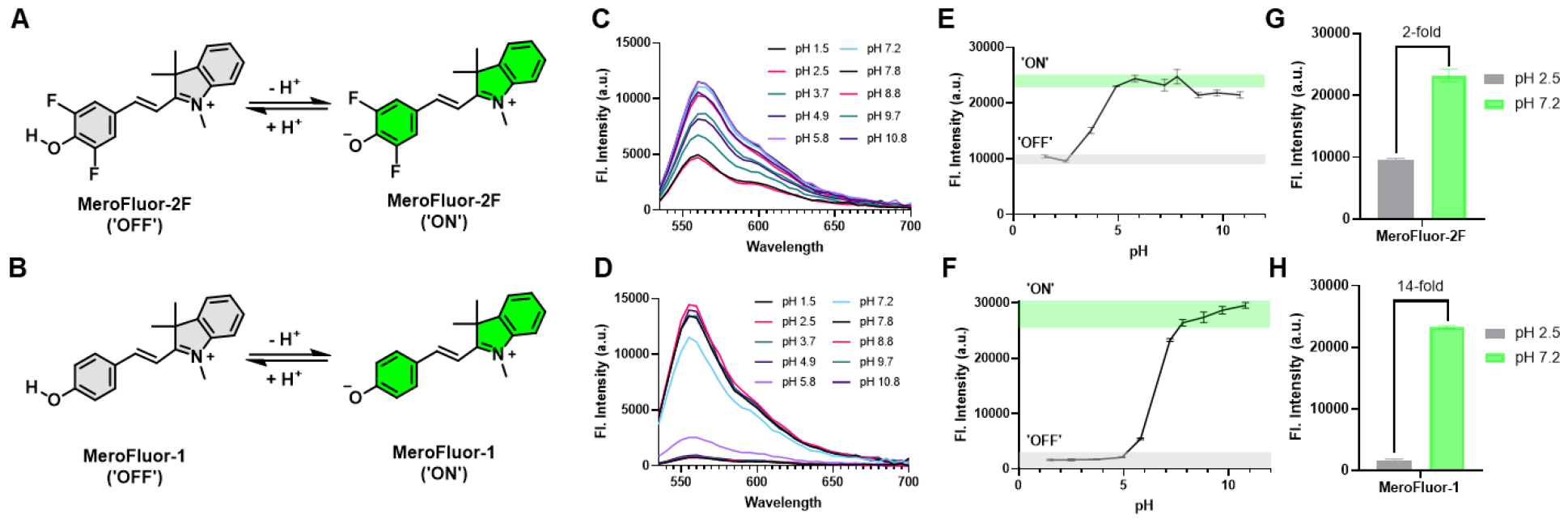
Comparison of the pH-dependent fluorescence properties of MeroFluor-2F and MeroFluor-1. The proposed reaction mechanism to elicit a fluorescence response via the ICT mechanism of (**A**) Merofluor-2F and (**B**) MeroFluor-1. The fluorescence emission spectra at various pH values of (**C**) MeroFluor-2F and (**D**) MeroFluor-1 (λ_ex_, 505 nm; λ_ex_, 540-700 nm). Quantification of the mean fluorescence intensity of (**E**) MeroFluor-2F and (**F**) MeroFluor-1 as a function of pH (λ_ex_,505 nm; λ_em_, 560 nm). ‘ON’/’OFF’ ratio (i.e., turn-on ratio) of (**G**) MeroFlour-2F and (**H**) MeroFluor-1 using pH 2.5 as ‘OFF’ and pH 7.2 as ‘ON’.

We also examined whether the aglycone affects the interaction with HPSE-1. To do so, we performed docking studies of **HUMPro** and **HAMPro** in the active site of HPSE-1 (**Figure 5**). Both compounds have similar binding modes and the aglycone oxygen linked to the cut site of HPSE-1 shows interaction with Glu225 and Glu343, the catalytic residues of HPSE-1. This observation reveals the cleavable nature of these probes. Importantly, the fluorine atoms on the phenolic oxygen of **HAMPro** do not seem to affect the binding of the dissacharide moiety with HPSE-1.

**Figure 5.**
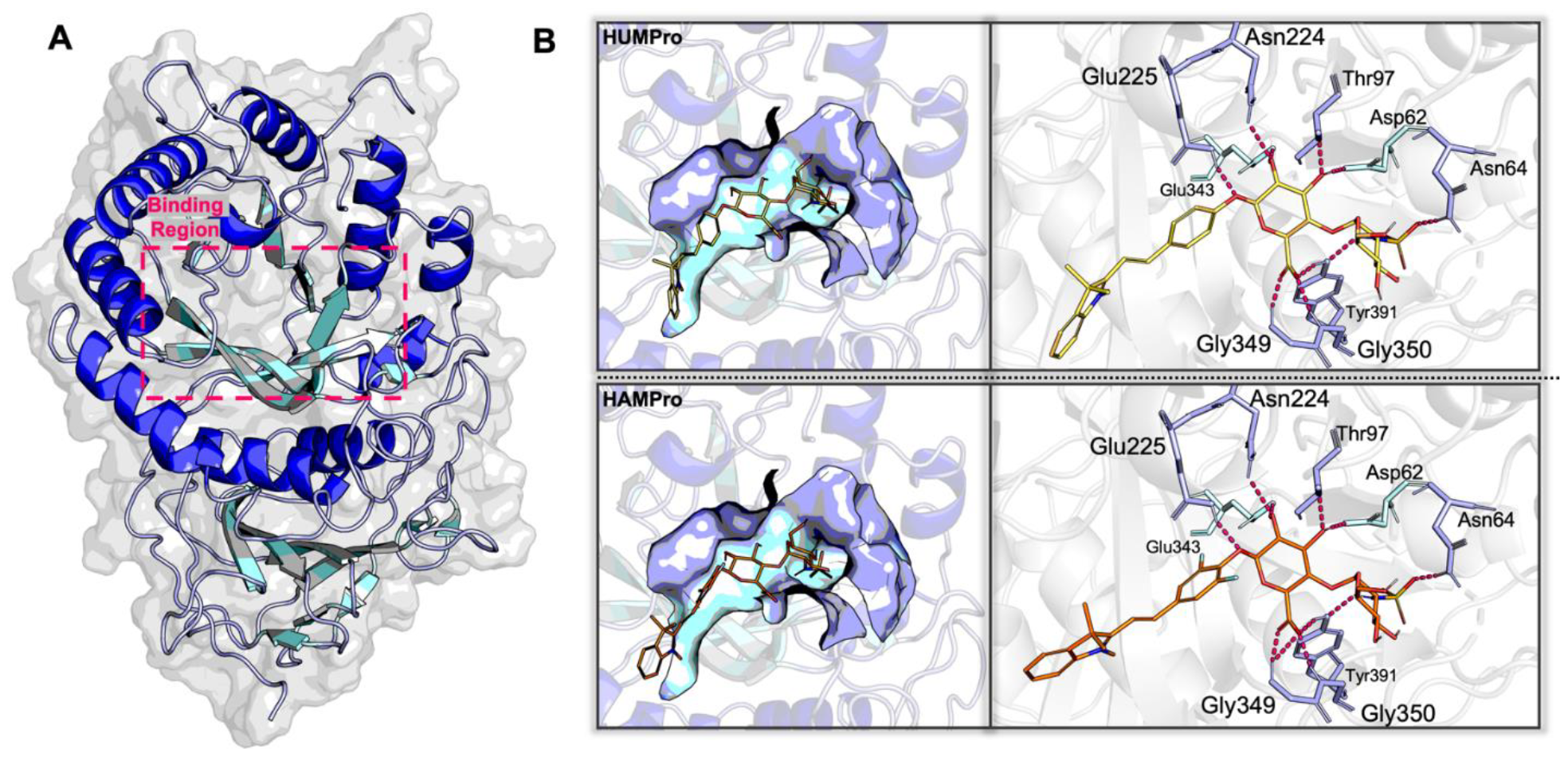
(**A**) The binding region of HPSE-1 we used for the binding mode study. (**B**) Detailed binding modes of **HUMPro** (upper panel) and **HAMPro** (bottom panel). The left-hand figures depict the overall fitness of the probe within the binding pocket. The figures on the right-hand side illustrate the detailed interaction diagrams of **HUMPro** and **HAMPro** within the active site of HPSE-1.

## Conclusion

Using the strategy of modifying the aryl aglycone leaving group to adjust the strength of the glycosidic linkage, we report the successful development of HAMPro, a novel green fluorogenic disaccharide-based probe to detect HPSE-1 activity. Despite not being stable for longer than 6 hours under ideal HPSE-1 enzymatic reaction condition, **HAMPro** was partially converted to **MeroFluor-2F** (64% conversion) in the presence of HPSE-1 for 6 hours, which showed an astoundingly sensitive 278-fold turn-on ratio. Furthermore, we provided some insight into the affect that modification of ICT-based fluorophores with electron-withdrawing groups has on fluorescence properties for future development of fluorogenic probes targeting HPSE-1; in some cases (**4MU**) it is beneficial by improving the fluorescence properties, whereas in other cases (**MeroFluor-1**) it is deleterious for sensitive detection. This poses a potential dilemma where tuning the electronics of the aglycone is necessary for activation but can impact the fluorescence properties in a negative manner. In the future, we aim to provide an in-depth analysis into the above phenomenon using computational studies. Nonetheless, **HAMPro** exhibited a highly sensitive turn-on ratio, which suggests that the previously described phenomenon may not significantly affect the fluorogenic properties of such probes. Although the aglycone must be able to be synthetically manipulated to tune the electronics to allow HPSE-1 activation, we expect that this study will highlight this proven design strategy as a considerable option to enable future development of HPSE-1 activity imaging probes. In our future work, we aim to optimize the probe design to improve the stability of the probe and detect extracellular HPSE-1 in the culture medium of living cells with high temporal and spatial resolution.

## Author contributions

ZMR and ZW contributed equally to the work. JL and LC designed the project. ZMR, ZW, JL, and MS synthesized and purified the compounds, and ZMR, ZW, and JL performed characterization. ZR performed the enzyme assays and evaluation of the fluorophores. AJY expressed the HPSE-1 enzyme. YZ performed computational studies. ZR and LC wrote the paper. All authors reviewed the manuscript and provided feedback.

## Acknowledgements

This work was supported by research grants to L.C. from the National Institute of General Medical Sciences (R35GM124963), the University of Florida (UF), and Maureen Keller-Wood PROSPER award (UF). Z.M.R. was partially supported by UF Graduate School Preeminent Scholarship.

