## Supporting Information for "A Fluorogenic Green Merocyanine-Based Probe to Detect Heparanase-1 Activity"

### for

### 1. Materials and Methods

**General Methods.** All required chemicals were procured from commercial providers and were used without further purification unless otherwise noted. Compound **1**[1] and **MeroFluor-1**[2] were prepared following conditions in the literature and their spectral characteristics agreed with literature values. High-performance liquid chromatography (HPLC) purification was performed on a Dionex Ultimate 3000 HPLC System (Thermo Scientific) equipped with HPG BX gradient pump and an in-line diode array UV-Vis detector. The preparative reversed-phase column used was a Luna® 5 µm C18(2) 100Å, LC column 250 x 21.2 mm (Phenomenex, Part No. 00G-4252-P0-AX). Liquid chromatography-Mass Spectrometry (LC-MS) analysis was performed on a Vanquish HPLC System (Thermo Scientific) equipped with VF gradient binary Pump and an in-line diode array UV-Vis detector coupled to an ISQ EM Mass Spectrometer System (Thermo Scientific). The analytical column used was an Accucore™ C18 HPLC column 50 mm x 4.6 mm, particle size 2.6 µm (Thermo Scientific, Part No. 17126-054630). On the Dionex Ultimate 3000 HPLC System, acetonitrile/water gradient mobile phase containing 0.1% trifluoroacetic acid, whereas on the Vanquish HPLC System, methanol/water gradient mobile phase containing 0.1% formic acid instead. High resolution mass was recorded on the Thermo Fisher Q-Exactive Focus Mass Spectrometer. Deuterated solvents were purchased from Sigma-Aldrich, Merck Millipore, and Acros Organics. NMR spectra were recorded on Bruker instruments (500 MHz and 600 MHz for $^{1}$H NMR; 126 MHz, 151 MHz, and 201 MHz for $^{13}$C NMR) and internally referenced to the residual solvent signals ($^{1}$H: δ 7.26, $^{13}$C: δ 77.16 for $CDCl_3$; $^{1}$H: δ 3.31, $^{13}$C: δ 49.0 for MeOD; $^{1}$H: δ 2.50, $^{13}$C: δ 39.52 for DMSO-*d6*). NMR chemical shifts (δ) and the coupling constants (J) for $^{1}$H and $^{13}$C NMR are reported in parts per million (ppm) and in Hertz, respectively. The following conventions are used for multiplicities: s, singlet; d, doublet; t, triplet; m, multiplet; dd, doublet of doublet. High resolution mass spectrometry was recorded on the Thermo Fisher Q-Exactive Focus Mass Spectrometer. Human Heparanase-1 (HPSE-1) was expressed and purified by reported protocol.[3]

**Assessing the reaction of HAMPro and HUMPro in the presence of recombinant HPSE-1.** To a solution of 3 µL of 50 µM each probe in 24 µL of freshly prepared 40 mM NaOAc buffer (pH 5.0) was added 3 µL of 1.5 µg/µL HPSE-1 to obtain 5 µM probe with 0.15 µg/µL recombinant human HPSE-1 in a total volume of 30 µL Each solution was prepared in triplicate with controls lacking HPSE-1. The solutions (30 µL) were charged into wells of a 384-well plate and then, the 384-well plate was incubated at 37 ºC in a $HERA_{THERM}$ oven (Thermo Scientific). The fluorescence intensity ($\lambda_{ex}/\lambda_{em}$, 505/560 nm, gain: 100) was measured after 5, 10, 30, 60, 120, 240, and 360 minutes using BioTek Synergy H1 plate reader (Agilent Technologies). After 6 hours, the fluorescence emission spectra ($\lambda_{ex}/\lambda_{em}$, 505/540-650 nm, step: 5 nm, gain: 100) of each well was measured. From this fluorescence emission spectra, the turn-on ratio of **HAMPro** (i.e., the fluorescence intensity of probe with HPSE-1 divided by that without HPSE-1) was calculated. The solutions from the wells were then collected and injected into LCMS to characterize the conversion products using the 400 nm channel. The following HPLC Method was used: 2-100%, 0-20 min; 100%, 20-22.5 min; MeOH/$H_2O$ with 0.1% Formic Acid.

**Fluorescence emission spectra at various pH values of Fluorophores.** Solutions of 10 µM **MeroFluor-2F**, **MeroFluor-1**, **DiFMU**, or **4MU** in Britton-Robinson (B&R) buffers (40 mM $H_3PO_4$, 40 mM $CH_3COOH$, 40 mM $H_3BO_3$) with various pH values (1.5, 2.5, 3.7, 4.9, 5.8, 7.8, 8.8, 9.7, and 10.8) were prepared in a final volume of 200 µL in triplicate. Additionally, solutions of 10 µM

**MeroFluor-2F**, **MeroFluor-1**, **DiFMU**, or **4MU** in phosphate-buffer saline (pH 7.2) were prepared in a final volume of 200 µL in triplicate. 150 µL of these solutions were added into a 96-well microplate. For **MeroFluor-2F** and **MeroFluor-1**, the fluorescence emission spectra ($\lambda_{ex}$ = 505 nm, $\lambda_{em}$ = 540-700 nm, step: 5 nm, gain: 100) and the fluorescence emission intensities ($\lambda_{ex}$ = 505 nm, $\lambda_{em}$ = 560 nm, gain: 100) were measured using a BioTek Synergy H1 plate reader. For **DiFMU** and **4MU**, the fluorescence emission spectra ($\lambda_{ex}$ = 365 nm, $\lambda_{em}$ = 400-500 nm, step: 5 nm, gain: 100) and the fluorescence emission intensities ($\lambda_{ex}$ = 365 nm, $\lambda_{em}$ = 455 nm, gain: 100) were measured on the same instrument. The fluorescence emission intensities were plotted as a function of pH.

**Molecular docking study of HAMPro and HUMPro with human HPSE-1.** The molecular docking study of **HUMPro** and **HAMPro** with HPSE-1 was performed using crystal structure obtained from Protein Data Bank at https://www.rcsb.org.[1, 2] The crystal structure 5E98[3] underwent a series of preparation procedures. The preparation procedures included removing water molecules and ions outside of the binding pocket; retaining only one unit of the tetramer. Then, the protein was further prepared following the preparation wizard workflow. The alternative residue positions were solved; Co-crystalized small molecules that locates outside of the binding pocket were removed. Hydrogens were added based on the protonation states of the protein and ligands at a pH of 6.5 ± 2.0. The IPT in the binding pocket was used for grid box generation, with an inner grid box size of 15 Å x 15 Å x 15 Å, buried in an outer box of 30 Å x 30 Å x 30 Å.  The OPLS 2005 force field was applied for the grid box generation. The probes 1 and 2 were prepared using Ligand Preparation module [4] in Schrödinger Maestro, and their pKa values and protonation states were determined under a pH of 7 using Epik.[5] Glide SP[6] was used for generating binding conformations of the prepared ligands with flexible ligand sampling method. The van der Waals radii were scaled with a scaling factor of 0.80, and a partial charge cutoff of 0.15 was applied. Additionally, nitrogen inversion and ring conformations sampling options were applied. Bias sampling of torsions option was kept for all predefined functional groups. Epik state penalties was added to docking score. Post-docking minimizations were performed following OPLS2005 force field. The generated conformations were inspected and chosen for generating interaction profile in PyMOL.[7]

### 2. Supporting Figures

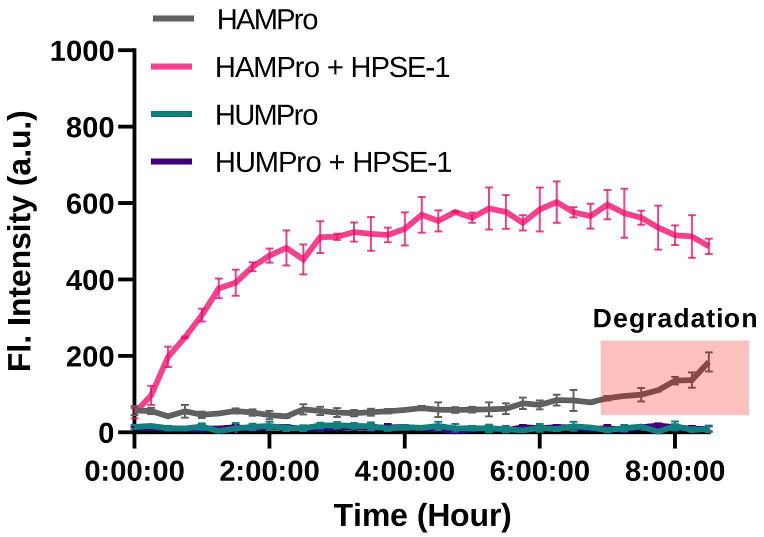

**Figure S1.** Time-course fluorescence assay of **HAMPro** and **HUMPro** without or with HPSE-1 to assess the stability of the probes under enzymatic buffer assay condition. To a solution of 3 µL of 50 µM each probe in 26 µL of 40 mM NaOAc (pH 5.0) was added 1 µL of 1.5 µg/µL HPSE-1 to obtain 5 µM probe with 0.05 µg/µL HPSE in a total volume of 30 µL Each solution was prepared in triplicate with controls lacking HPSE-1. The solutions (30 µL) were charged into wells of a 384-well plate. The fluorescence intensity ($\lambda_{ex}/\lambda_{em}$, 505/560 nm) was measured every 15 minutes for 8.5 hours, while being kept at 37 ºC using BioTek Synergy H1 plate reader (Agilent Technologies).

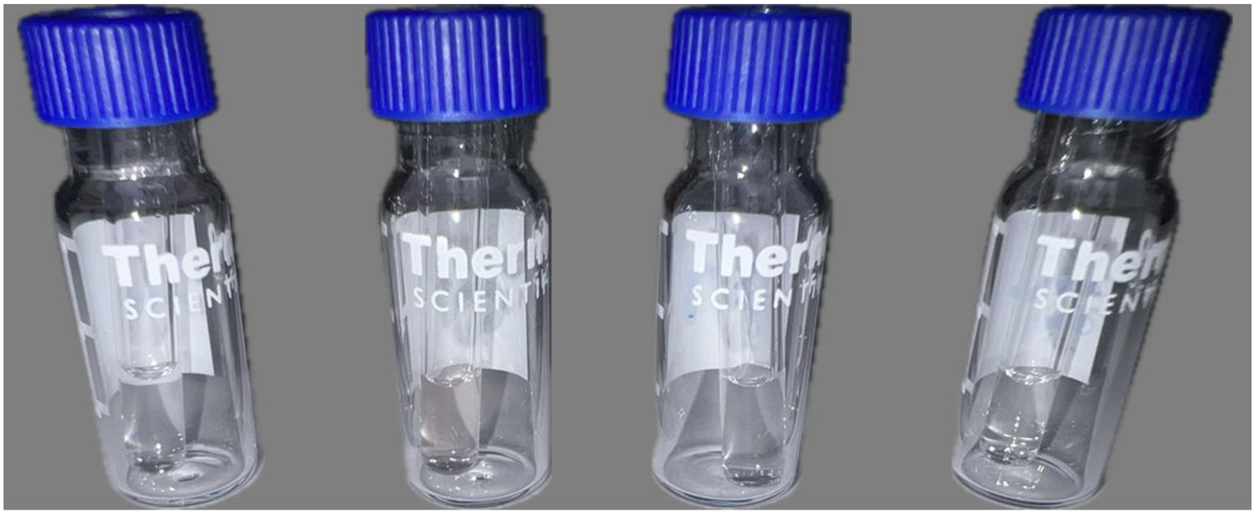

HAMPro HAMPro + HPSE-1 HUMPro HUMPro + HPSE-1

**Figure S2.** Pictures of solutions after reaction of **HAMPro** and **HUMPro** (Final Concentration, 5 µM) without or with HPSE-1 (Final Concentration, 0.15 µg/µL) in 40 mM NaOAc buffer (pH 5.0) at 37°C for 6 hours.

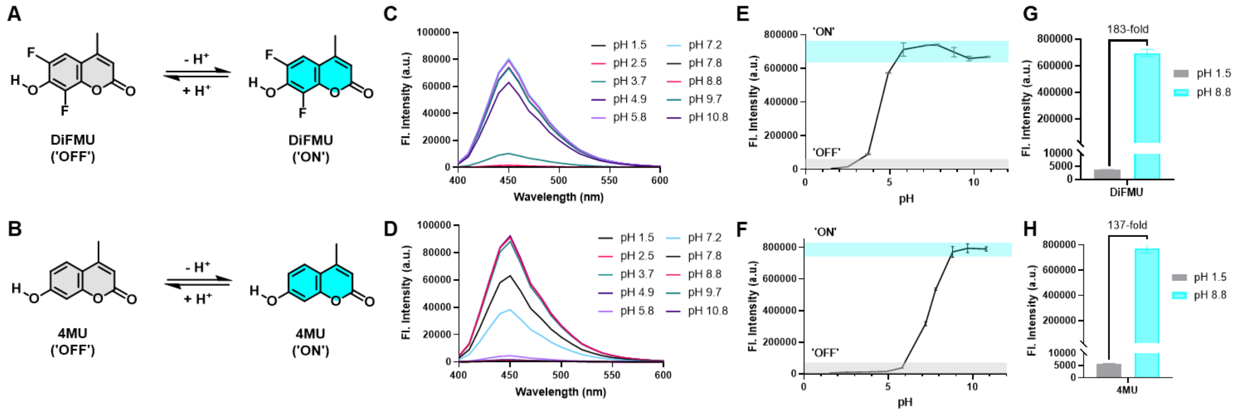

**Figure S3.** Comparison of the pH-dependent fluorescence properties of **4MU** and **DiFMU**. The proposed reaction mechanism to elicit a fluorescence response via the pH-dependent ICT mechanism of (**A**) **DiFMU** and (**B**) **4MU**. The fluorescence emission spectra ($\lambda_{ex}$, 365 nm; $\lambda_{ex}$, 400-600 nm) at various pH values of (**C**) **DiFMU** and (**D**) **4MU** Quantification of the mean fluorescence intensity of (**E**) **DiFMU** and (**F**) **4MU** as a function of pH ($\lambda_{ex}$,365 nm; $\lambda_{em}$, 455 nm). 'ON'/'OFF' ratio (i.e., turn-on ratio) of (**G**) **DiFMU** and (**H**) **4MU** using pH 1.5 as 'OFF' and pH 8.8 as 'ON'.

### 3. Synthesis and Characterization

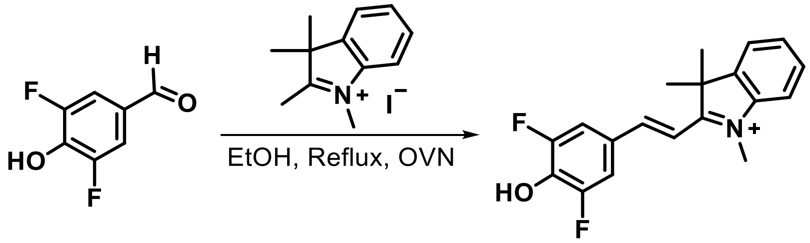

**MeroFluor-2F:** 1,2,3,3-tetramethyl-*3H*-Indol-1-ium Iodide (31. mg, 0.180 mmol, 1 eq) and 3,5-difluoro-4-hydroxybenzaldehyde (31.6 mg, 0.181 mmol, 1 eq) was added to a 100 mL round-bottom flask, followed by anhydrous EtOH (10 mL). The reaction flask was placed in an oil bath under a condenser at reflux (80-90°C). The reaction was allowed to stir overnight. After the reaction was complete as indicated by TLC analysis (10% MeOH/DCM), the reaction solution was purified by silica gel column (5% MeOH/DCM to 10% MeOH/DCM) to give **MeroFluor-2F** as a red powder (Yield: 92%, 52.1 mg).

**$^1$H NMR (600 MHz, DMSO-*d6*)** δ 8.30 (d, *J* = 16.3 Hz, 1H, **H-e**), 8.07 (d, *J* = 9.8 Hz, 2H, **H-c**), 7.93 – 7.83 (m, 2H, **H-m**, **H-j**), 7.66 – 7.55 (m, 2H, **H-k**, **H-l**), 7.58 (d, *J* = 16.3 Hz, 1H, **H-f**), 4.13 (s, 3H, -$N^+CH_3$), 1.76 (s, 6H, 2 x –$CH_3$).

**$^{13}$C NMR (151 MHz, DMSO-*d6*)** δ 181.42 (**C-g**), 153.10 (**C-n**), 151.49 (**C-e**), 151.29 (**C-a**), 142.70 (d, *J* = 256.5 Hz, **C-b**), 139.33 (**C-d**), 129.29 (**C-k**), 128.97 (**C-l**), 122.85 (**C-m**), 115.11 (**C-j**), 114.42 (**C-c**), 112.19 (**C-f**), 52.04 (-$N^+CH_3$), 34.38 (**C-h**), 25.30 (2 x $CH_3$).

**HRMS** *m/z* $[M]^+$ = 314.1351, calc'd for $C_{19}H_{18}F_2NO^+$, found: 314.1350 ($[M]^+$).

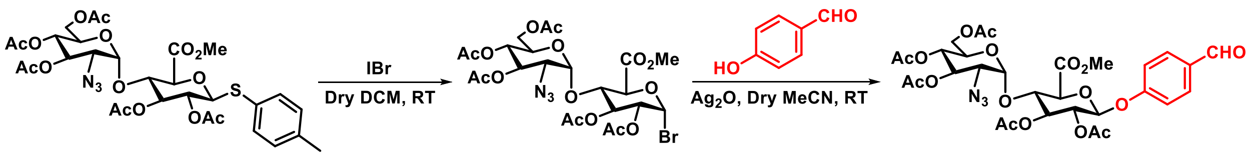

**Compound 2:** To a solution of compound **1** (1.13 g, 1.19 mmol) in dry DCM (10 mL), IBr (493. mg, 2.39 mmol, 1.5 eq) was added. The reaction was allowed to proceed for 10 minutes and then, quenched with a saturated solution of $NaS_2O_3$. The solution was then extracted with DCM three times, washed with brine, and concentrated in-vacuo. The crude product was purified by silica gel column (2:1 to 1:1 Hexanes/Ethyl Acetate) to give the glycosyl bromide intermediate as a white powder (Yield: 723 mg, 68%). The glycosyl bromide intermediate (668 mg, 1 mmol) was immediately dissolved in dry acetonitrile (5 mL). To this solution, 4-hydroxybenzaldehyde (138 mg, 1.30 mmol, 1.3 eq) and silver oxide (463 mg, 2 mmol, 2 eq) was added. The reaction was allowed to stir at room temperature for 24 hours. After the reaction was complete as indicated by TLC, the solution was filtered through Celite®, concentrated in-vacuo, and then purified by silica gel column (2:1 to 1.5:1 Hexanes: Ethyl Acetate) to give the compound **2** as a white powder (Yield: 411 mg, 58%).

**$^{1}$H NMR (600 MHz, $CDCl_3$)** δ 9.92 (s, 1H, -C**H**O), 7.86 (d, *J* = 8.7 Hz, 2H, **H-c**), 7.09 (d, *J* = 8.7 Hz, 2H, **H-b**), 5.42 – 5.32 (m, 3H, **H-1'**, **H-3**, **H-3'**), 5.26 (d, *J* = 3.8 Hz, 1H, **H-1**), 5.18 (dd, *J* = 8.3, 6.0 Hz, 1H, **H-2**), 5.02 (dd, *J* = 10.3, 9.3 Hz, 1H, **H-4'**), 4.45 (t, *J* = 8.6 Hz, 1H, **H-4**), 4.30 (d, *J* = 8.5 Hz, 1H, **H-5**), 4.24 (dd, *J* = 12.6, 3.8 Hz, 1H, **H-6'**), 4.11 (dd, *J* = 12.5, 2.3 Hz, 1H, **H-6'**), 3.90 (ddd, *J* = 10.4, 3.7, 2.2 Hz, 1H, **H-5'**), 3.64 (s, 3H, $-CO_2$**Me**), 3.42 (dd, *J* = 10.7, 3.8 Hz, 1H, H-2'), 2.10 (s, 3H, -OCO**Me**), 2.08-2.07 (m, 9H, 3 x -OCO**Me**), 2.02 (s, 3H, -OCO**Me**).

**$^{13}$C NMR (151 MHz, $CDCl_3$)** δ 190.84 (**C**HO), 170.77 (-O**C**OMe), 170.02 (-O**C**OMe), 169.74 (-O**C**OMe), 169.72 (-O**C**OMe), 167.68 (-**C**$O_2$Me), 161.04 (**C-a**), 131.99 (**C-c**), 131.96 (**C-d**), 116.76 (**C-b**), 98.94 (**C-1**), 97.93 (**C-1'**), 74.99 (**C-4**), 74.19 (**C-5**), 72.83 (**C-3**), 71.54 (**C-2**), 70.21 (**C-3'**), 68.68 (**C-5'**), 68.20 (**C-4'**), 61.34 (**C-6'**), 61.01 (**C-2'**), 52.98 ($-CO_2$**Me**), 20.84 (-OCO**Me**), 20.80 (-OCO**Me**), 20.75 (-OCO**Me**), 20.69 (-OCO**Me**).

**HRMS** *m/z* $[M]^+$ = 709.1966, calc'd for $C_{30}H_{35}N_3O_{17}$, found: 732.1848 ($[M+Na]^+$), 748.1800 ($[M+K]^+$).

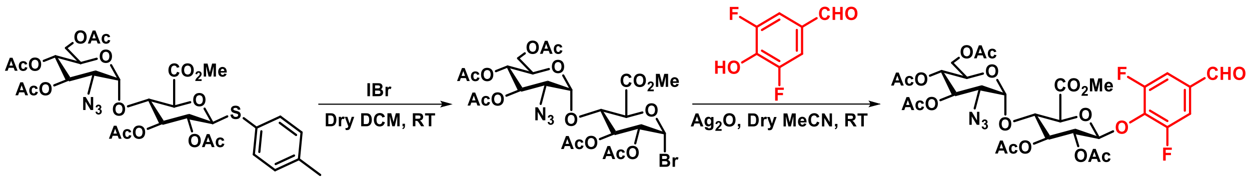

**Compound 3:** To a solution of compound **1** (2.12 g, 2.98 mmol) in dry DCM (10 mL), IBr (1.07 g, 5.17 mmol, 1.74 eq) was added. The reaction was allowed to proceed for 10 minutes and then, quenched with a saturated solution of $NaS_2O_3$. The solution was then extracted with DCM three times, washed with brine, and concentrated in-vacuo. The crude was purified by silica gel column (2:1 to 1:1 Hexanes/ Ethyl Acetate) to give the glycosyl bromide intermediate as a white powder (Yield: 1.74 g, 87%). To a solution of the glycosyl bromide intermediate (722.7 mg, 1.08 mmol, 1 eq) in dry acetonitrile (2 mL), 3,5-difluoro-4-hydroxybenzaldehyde (175.0 mg, 1.1 mmol, 1 eq) and silver oxide (504.1 mg, 2.18 mmol, 2 eq) was added. The reaction was allowed to stir at room temperature for 20 hours. After the reaction was complete as indicated by TLC, the solution was filtered through Celite®, concentrated in-vacuo, and then purified by silica gel column (2:1 Hexanes: Ethyl Acetate to 1:1 Hexanes: Ethyl Acetate) to give compound **3** as a white powder (Yield: 707 mg, 88%).

**$^1$H NMR (600 MHz, $CDCl_3$)** δ 9.86 (s, 1H, -C**H**O), 7.46 (d, $J$ = 7.4 Hz, 2H, **H-c**), 5.36 – 5.29 (m, 3H, **H-1'**, **H-3**, **H-3'**), 5.24 – 5.18 (m, 2H, **H-1**, **H-2**), 5.01 (dd, $J$ = 10.3, 9.3 Hz, 1H, **H-4'**), 4.38 (t, $J$ = 8.8 Hz, 1H, **H-4**), 4.23 (dd, $J$ = 12.6, 3.7 Hz, 1H, **H-6'**), 4.14 (d, $J$ = 9.1 Hz, 1H, **H-5**), 4.08 (dd, $J$ = 12.6, 2.2 Hz, 1H, **H-6'**), 3.84 (ddd, $J$ = 10.3, 3.6, 2.2 Hz, 1H, **H-5'**), 3.75 (s, 3H, -$CO_2$**Me**), 3.41 (dd, $J$ = 10.7, 3.7 Hz, 1H, **H-2'**), 2.09 (s, 6H, 2 x -OCO**Me**), 2.07 (s, 3H, -OCO**Me**), 2.06 (s, 3H, -OCO**Me**), 2.01 (s, 3H, -OCO**Me**).

**$^{13}$C NMR (151 MHz, $CDCl_3$)** δ 188.77 (-**C**HO), 170.89 (-O**C**OMe), 170.06 (-O**C**OMe), 169.84 (-O**C**OMe)), 169.79 (-O**C**OMe), 169.71 (-**C**$O_2$Me), 155.69 (dd, $J$ = 254.0, 4.2 Hz, **C-b**), 137.56 (t, $J$ = 14.1 Hz, **C-a**), 132.55 (t, $J$ = 6.5 Hz, **C-d**), 113.50 (dd, $J$ = 18.5, 4.8 Hz, **C-c**), 100.96 (**C-1'**), 98.87 (**C-1**), 75.61 (**C-4**), 74.40 (**C-5**), 73.23 (**C-3**), 71.41 (**C-2**), 70.24 (**C-3'**), 68.68 (**C-5'**), 68.11 (**C-4'**), 61.27 (**C-6'**), 60.99 (**C-2'**), 53.13 (-$CO_2$**Me**), 20.79 (-OCO**Me**), 20.73 (-OCO**Me**), 20.66 (-OCO**Me**).

**HRMS** $m/z$ $[M]^+$ = 745.1778, calc'd for $C_{30}H_{33}F_2N_3O_{17}$, found: 745.5008 ($[M]^+$).

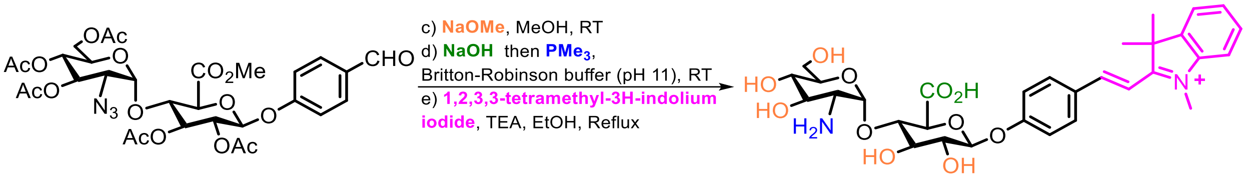

**Compound 4:** To a solution of compound **2** (150. mg, 0.211 mmol) in dry MeOH (2 mL), NaOMe (25 wt% in MeOH, 48 µL, 0.211 mmol, 1 eq) was added. The reaction was allowed to proceed for 2 hours and then, quenched with DOWEX ® 50WX8 100-200 mesh ion-exchange resin (CAS: 11119-67-8). The solution was filtered through Celite, which was washed with MeOH, and then concentrated *in-vacuo*. The crude was immediately used for the next step. The crude was dissolved in an aqueous Britton-Robinson buffer (pH 11, 2 mL) and then, NaOH (1M, 206 µL, 0.206 mmol, 0.98 eq) was added. After overnight, the reaction was complete as indicated by LC-MS analysis. To the same pot, $PMe_3$ (1M in THF, 422 µL, 0.422 mmol, 2 eq) was added. After overnight, the reaction was complete as indicated by LC-MS analysis. The reaction was then purified immediately by preparative HPLC (2-100%, 0-20 minutes, MeCN/$H_2O$ with 0.1% TFA) and lyophilized to give the product of step d) as a white powder (Yield: 43. mg, 44% over two steps). The product was then immediately dissolved in anhydrous EtOH (4 mL) and 1,2,3,3-tetramethyl-*3H*-indolium iodide (60. mg, 0.20 mmol, 2.1 eq) and TEA (26 µL, 0.188 mmol, 2 eq) was added. Then, the solution was placed in an oil bath at reflux (80-90°C) under a condenser overnight. After overnight, the reaction was complete as indicated by LC-MS analysis and the reaction was then purified immediately by preparative HPLC (2-100%, 0-20 minutes, MeCN/$H_2O$ with 0.1% TFA) and lyophilized to give compound **4** as a dark-red powder (Yield: 29.5 mg, 51%).

**$^1$H NMR (600 MHz, MeOD)** δ 8.40 (d, *J* = 16.3 Hz, 1H, **H-e**), 8.07 (d, *J* = 9.0 Hz, 2H, **H-c**), 7.82 – 7.79 (m, 1H, **H-m**), 7.77 – 7.74 (m, 1H, **H-l**), 7.66 – 7.63 (m, 2H, **H-j**, **H-k**), 7.55 (d, *J* = 16.3 Hz, 1H, **H-f**), 7.25 (d, *J* = 8.9 Hz, 2H, **H-b**), 5.57 (d, *J* = 3.7 Hz, 1H, **H-1'**), 5.19 (d, *J* = 7.7 Hz, 1H, **H-1**), 4.24 (d, *J* = 9.6 Hz, 1H, **H-3**), 4.15 (s, 3H, -$N^+C$**$H_3$**), 3.96 (t, *J* = 9.4 Hz, 1H, **H-3'**), 3.83 – 3.72 (m, 5H, **H-2**, **H-4'**, **H-6'**), 3.65 – 3.58 (m, 2H, **H-4**, **H-5**), 3.50 (dd, *J* = 10.0, 8.9 Hz, 1H, **H-5'**), 3.16 (dd, *J* = 10.7, 3.7 Hz, 1H, **H-2'**), 1.84 (s, 6H, 2 x –C**$H_3$**).

**$^{13}$C NMR (151 MHz, MeOD)** δ 183.93 (**C-g**), 171.30 (-**C**$O_2$H), 163.10 (**C-a**), 155.27 (**C-e**), 144.76 (**C-n**), 143.23 (**C-i**), 133.66 (**C-c**), 130.85 (**C-d**), 130.50 (**C-k**), 130.35 (**C-j**), 123.89 (**C-l**), 118.39 (**C-b**), 115.84 (**C-m**), 111.90 (**C-f**), 101.29 (**C-1**), 98.41 (**C-1'**), 79.18 (**C-3'**), 77.12 (**C-4'**), 75.56 (**C-3**), 74.56 (**C-5**), 74.46 (**C-4**), 71.09 (**C-2**), 70.83 (**C-5'**), 61.28 (**C-6'**), 56.12 (**C-2'**), 53.81 (**C-h**), 34.56 (-$N^+$**C**$H_3$), 26.33 (2 x -C**$H_3$**).

**HRMS** *m/z* $[M]^+$ = 615.2548, calc'd for $C_{31}H_{39}N_2O_{11}^+$, found: 615.2557 ($[M]^+$).

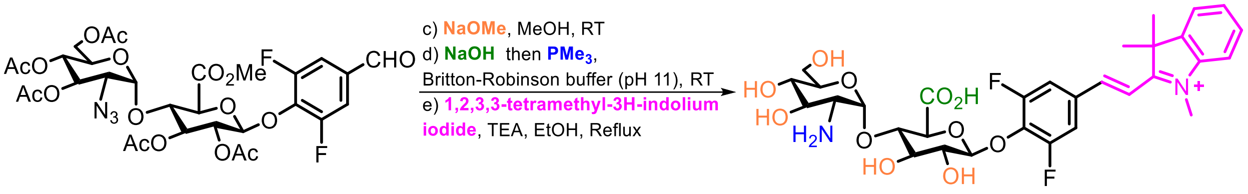

**Compound 5:** To a solution of compound **3** (160. mg, 0.215 mmol) in dry MeOH (2 mL), NaOMe (25 wt% in MeOH, 49 µL, 0.215 mmol, 1 eq) was added. The reaction was allowed to proceed for 2 hours and then, quenched with DOWEX ® 50WX8 100-200 mesh ion-exchange resin (CAS: 11119-67-8). The solution was filtered through Celite, which was washed with MeOH, and then concentrated *in-vacuo*. The crude was immediately used for the next step. The crude material was dissolved in an aqueous Britton-Robinson buffer (pH 11, 2 mL) and then, NaOH (1M, 208 µL, 0.208 mmol, 0.97 eq) was added. After overnight, the reaction was complete as indicated by LC-MS analysis. To the same pot, $PMe_3$ (1M in THF, 430 µL, 0.430 mmol, 2 eq) was added. After overnight, the reaction was complete as indicated by LC-MS analysis. The reaction was then purified immediately by preparative HPLC (2-100%, 0-20 minutes, $MeCN/H_2O$ with 0.1% TFA) and lyophilized to give the product of step d) as a white powder (Yield: 43. mg, 31% over 3 steps). The product was then immediately dissolved in anhydrous EtOH (4 mL) and 1,2,3,3-tetramethyl-*3H*-indolium iodide (43. mg, 0.14 mmol, 2.1 eq) and TEA (19 µL, 0.134 mmol, 2 eq) was added. Then, the solution was placed in an oil bath at reflux (80-90°C) under a condenser overnight. After overnight, the reaction was complete as indicated by LC-MS analysis and then the reaction was then purified immediately by preparative HPLC (2-100%, 0-20 minutes, $MeCN/H_2O$ with 0.1% TFA) and lyophilized to give compound **5** as a dark-red powder (Yield: 20.7 mg, 47%).

**$^1$H NMR (600z MHz, DMSO-*d6*)** δ 8.31 (d, *J* = 16.4 Hz, 1H, **H-e**), 8.14 (d, *J* = 9.4 Hz, 2H, **H-c**), 7.97 (d, *J* = 5.3 Hz, 2H, -N**H**$_2$), 7.96 – 7.89 (m, 1H, **H-m**), 7.91 – 7.85 (m, 1H, **H-l**), 7.71 (d, *J* = 16.4 Hz, 1H, **H-f**), 7.68 – 7.59 (m, 2H, **H-j**, **H-k**), 5.47 (d, *J* = 3.7 Hz, 1H, **H-1'**), 5.15 (d, *J* = 7.7 Hz, 1H, **H-1**), 4.18 (s, 3H, -N$^+$C**H**$_3$), 3.91 (d, *J* = 9.5 Hz, 1H, **H-3**), 3.77 (t, *J* = 9.2 Hz, 1H, **H-3'**), 3.69 (t, *J* = 8.9 Hz, 1H, **H-4'**), 3.60 (dd, *J* = 12.0, 2.7 Hz, 1H, **H-6'**), 3.58 – 3.49 (m, 2H, **H-6'**, **H-2**), 3.42 (dd, *J* = 16.8, 8.2 Hz, 1H, **H-5**), 3.39 – 3.32 (m, 2H, **H-4**, **H-2'**), 2.99 (dt, *J* = 10.2, 4.8 Hz, 1H, **H-5'**), 1.77 (s, 6H, 2 x –C**H**$_3$).

**$^{13}$C NMR (151 MHz, DMSO-*d6*)** δ 181.69 (**C-g**), 169.06 (-**C**O$_2$H), 155.77 (**C-b**), 154.12 (**C-a**), 149.63 (**C-e**), 143.84 (**C-n**), 141.83 (**C-i**), 130.90 (**C-k**), 129.83 (**C-j**),122.94 (**C-l**), 117.61 (**C-m**), 115.64 (**C-d**), 115.57 (**C-c**), 114.84 (**C-f**), 103.28 (**C-1**), 95.93 (**C-1'**), 76.94 (**C-3'**), 75.30 (**C-4'**), 74.49 (**C-3**), 73.16 (**C-5**), 73.12 (**C-2'**), 69.46 (**C-2**), 68.86 (**C-4**), 59.28 (**C-6'**), 54.33 (**C-5'**), 52.42 (**C-h**), 34.84 (-N$^+$CH$_3$), 25.01 (–**C**H$_3$), 25.00 (–**C**H$_3$).

**HRMS** *m/z* $[M]^+$ = 651.2360, calc'd for $C_{31}H_{37}F_2N_2O_{11}^+$, found: 651.2357 ($[M]^+$).

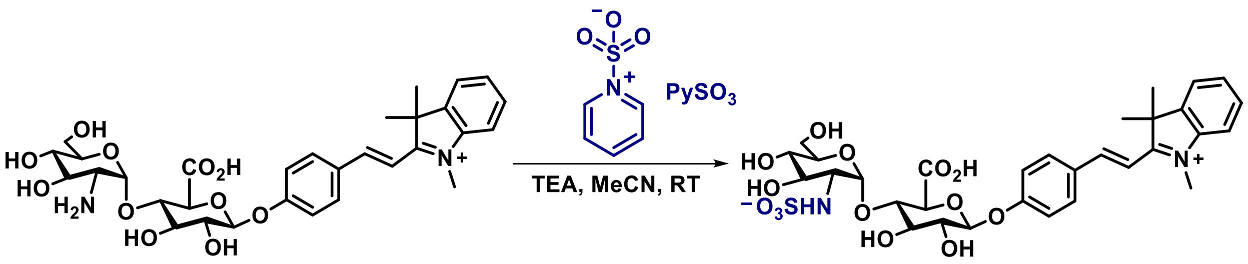

**Compound 6:** To a solution of compound **4** (29.5 mg, 0.048 mmol) in MeCN (2 mL), TEA (200 µL) and $PySO_3$ (44 mg, 0.28 mmol, 5.8 eq) was added. The reaction was allowed to stir overnight. After the reaction was complete as indicated by LC-MS, the solution was immediately purified by preparative HPLC (2-70%, 0-30 min; 70%, 30-40 min; MeCN/H2O with 0.1% TFA). The aliquots containing product (verified by LC-MS) were diluted with $H_2O$, concentrated *in-vacuo* to remove MeCN, and then passed through a Sep-Pak® Plus Short C-18 cartridge, and lyophilized to give compound **6** (**HUMPro**) as yellow fluffy solid (Yield: 7.5 mg, 23%).

**$^1$H NMR (600z MHz, MeOD)** δ 8.41 (d, J = 16.2 Hz, 1H, **H-e**), 8.12 – 8.02 (m, 2H, **H-c**), 7.84 – 7.80 (m, 1H, **H-m**), 7.80 – 7.75 (m, 1H, **H-i**), 7.69 – 7.61 (m, 2H, **H-j**, **H-k**), 7.56 (d, J = 16.2 Hz, 1H, **H-f**), 7.31 – 7.21 (m, 2H, **H-b**), 5.67 (d, J = 3.8 Hz, 1H, **H-1'**), 5.24 (d, J = 7.7 Hz, 1H, **H-1**), 4.19 (d, J = 9.5 Hz, 1H, **H-5**), 4.17 (s, 3H, -$N^+$C**H**3), 3.97 (t, J = 9.2 Hz, 1H, **H-4**), 3.89 (t, J = 9.0 Hz, 1H, **H-3**), 3.79 (dd, J = 12.0, 2.6 Hz, 1H, **H-6'**), 3.75 (dd, J = 12.0, 3.9 Hz, 1H, **H-6'**), 3.65 – 3.54 (m, 3H, **H-2**, **H-5'**, **H-3'**), 3.48 (dd, J = 10.0, 8.9 Hz, 1H, **H-4'**), 3.30 (dd, J = 10.3, 3.8 Hz, 1H, **H-2'**), 1.86 (s, 6H, 2 x –C**H**3).

**$^{13}$C NMR (151 MHz, MeOD)** δ 183.92 (**C-g**), 172.20 (-**C**$O_2$H), 163.32 (**C-a**), 155.49 (**C-e**), 144.77 (**C-n**), 143.26 (**c-i**), 133.70 (**C-c**), 130.74 (**C-d**), 130.46 (**C-k**), 130.13 (**C-j**), 123.87 (**C-l**), 118.44 (**C-b**), 115.81 (**C-m**), 111.73 (**C-f**), 101.14 (**C-1**), 99.80 (**C-1'**), 78.75 (**C-4**), 77.40 (**C-3**), 76.29 (**C-5**), 74.16 (**C-2**), 73.63 (**C-5'**), 73.49 (**C-3'**), 71.54 (**C-4'**), 62.16 (**C-6'**), 59.98 (**C-2'**), 53.77 (**C-h**), 34.58 (-$N^+$**C**$H_3$), 26.38 (2 x -**C**$H_3$).

**HRMS** *m/z* $[M]^+$ = 694.2044, calc'd for $C_{31}H_{38}N_2O_{14}S$, found: 695.2108 ($[M+H]^+$), 717.1925 ($[M+Na]^+$).

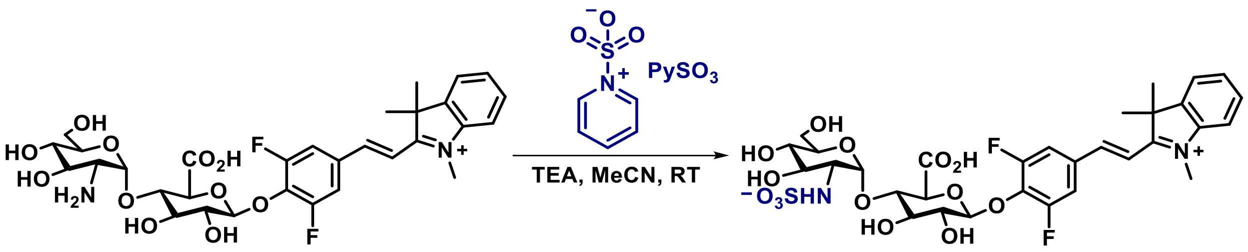

**Compound 7:** To a solution of compound **5** (20.7 mg, 0.032 mmol) in MeCN (2 mL), TEA (200 µL) and $PySO_3$ (33 mg, 0.21 mmol, 6.6 eq) was added. The reaction was allowed to stir overnight. After the reaction was complete as indicated by LC-MS, the solution was immediately purified by preparative HPLC (2-70%, 0-30 min; 70%, 30-40 min; MeCN/H2O with 0.1% TFA). The aliquots containing product (verified by LC-MS) were diluted with $H_2O$, concentrated *in-vacuo* to remove MeCN, and then passed through a Sep-Pak® Plus Short C-18 cartridge, and lyophilized to give compound **7** (**HAMPro**) as yellow fluffy solid (Yield: 3.4 mg, 15%).

**$^1$H NMR (600 MHz, MeOD)** δ 8.32 – 8.26 (m, 1H, **H-e**), 7.88 – 7.85 (m, 1H, **H-m**), 7.81 (d, *J* = 8.9 Hz, 2H, **H-c**), 7.80 – 7.77 (m, 1H, **H-k**), 7.69 – 7.65 (m, 2H, **H-i**, **h-j**), 7.63 (d, *J* = 16.3 Hz, 1H, **H-f**), 5.63 (d, *J* = 3.8 Hz, 1H, **H-1'**), 5.24 (d, *J* = 7.7 Hz, 1H, **H-1**), 4.21 (s, 3H, -$N^+CH_3$), 3.99 (d, *J* = 9.6 Hz, 1H, **H-4**), 3.94 (dd, *J* = 9.6, 8.5 Hz, 1H, **H-5**), 3.84 (t, *J* = 8.9 Hz, 1H, **H-3**), 3.76 – 3.70 (m, 2H, 2 x **H-6'**), 3.58 (dd, *J* = 9.1, 7.7 Hz, 1H, **H-2**), 3.53 (ddd, *J* = 8.8, 6.0, 3.0 Hz, 1H, **H-5'**), 3.51 – 3.44 (m, 2H, **H-3'**, **H-4'**), 3.26 (dd, *J* = 10.1, 3.8 Hz, 1H, **H-2'**), 1.83 (s, 6H, 2 x – $CH_3$).

**$^{13}$C NMR (151 MHz, MeOD)** δ 183.97 (**C-g**), 171.54 (-$CO_2H$), 157.73 (**C-b**), 156.13 (**C-a**), 152.23 (**C-e**), 145.14 (**C-n**), 143.20 (**C-i**), 137.78 (**C-m**), 131.87 (**C-k**), 131.40 (**C-j**), 130.60 (**C-l**), 123.98 (**C-d**), 116.35 (**C-c**), 115.17 (**C-f**), 104.46 (**C-1'**), 99.88 (**C-1**), 78.78 (**C-3'**), 77.31 (**C-4'**), 76.39 (**C-3**), 74.82 (**C-5**), 73.64 (**C-2'**), 73.43 (**C-2**), 71.39 (**C-4**), 62.05 (**C-6'**), 54.17 (**C-5'**), 49.57 (**C-h**), 35.13 (-$N^+CH_3$), 25.90 (2 x -$CH_3$).

**$^{19}$F NMR (565 MHz, MeOD)** δ -126.62 (d, *J* = 9.0 Hz, $C_b$-F).

**HRMS** *m/z* $[M]^+$ = 730.1855, calc'd for $C_{31}H_{36}F_2N_2O_{14}S$, found: 731.1931 ($[M+H]^+$), 753.1747 ($[M+Na]^+$).

### 4. NMR Spectra, HRMS Spectra, and Purity Analysis (TLC or HPLC)

#### A. MeroFluor-2F

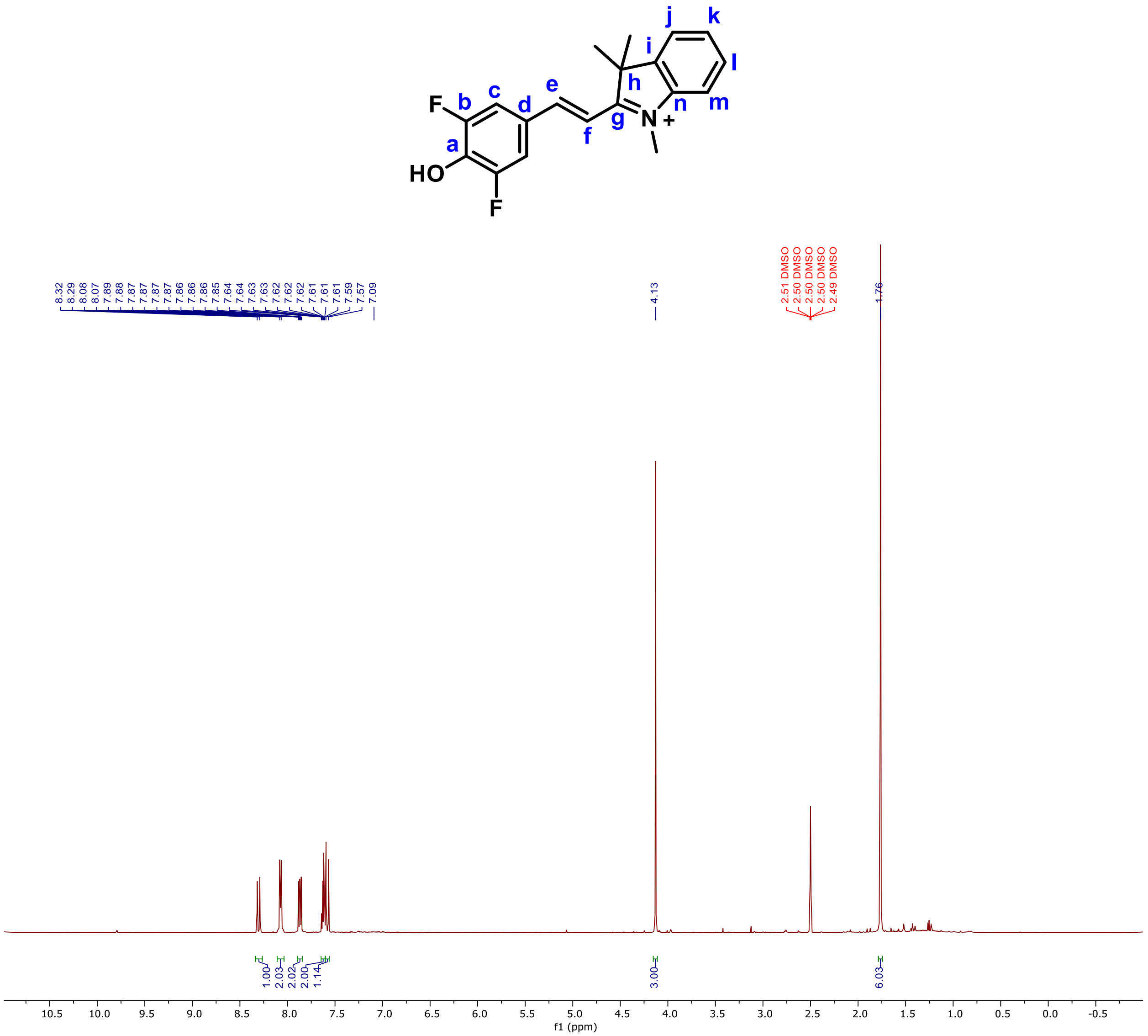

**Figure S4**. $^{1}H$ NMR spectrum of **MeroFluor-2F** in DMSO-*d6*

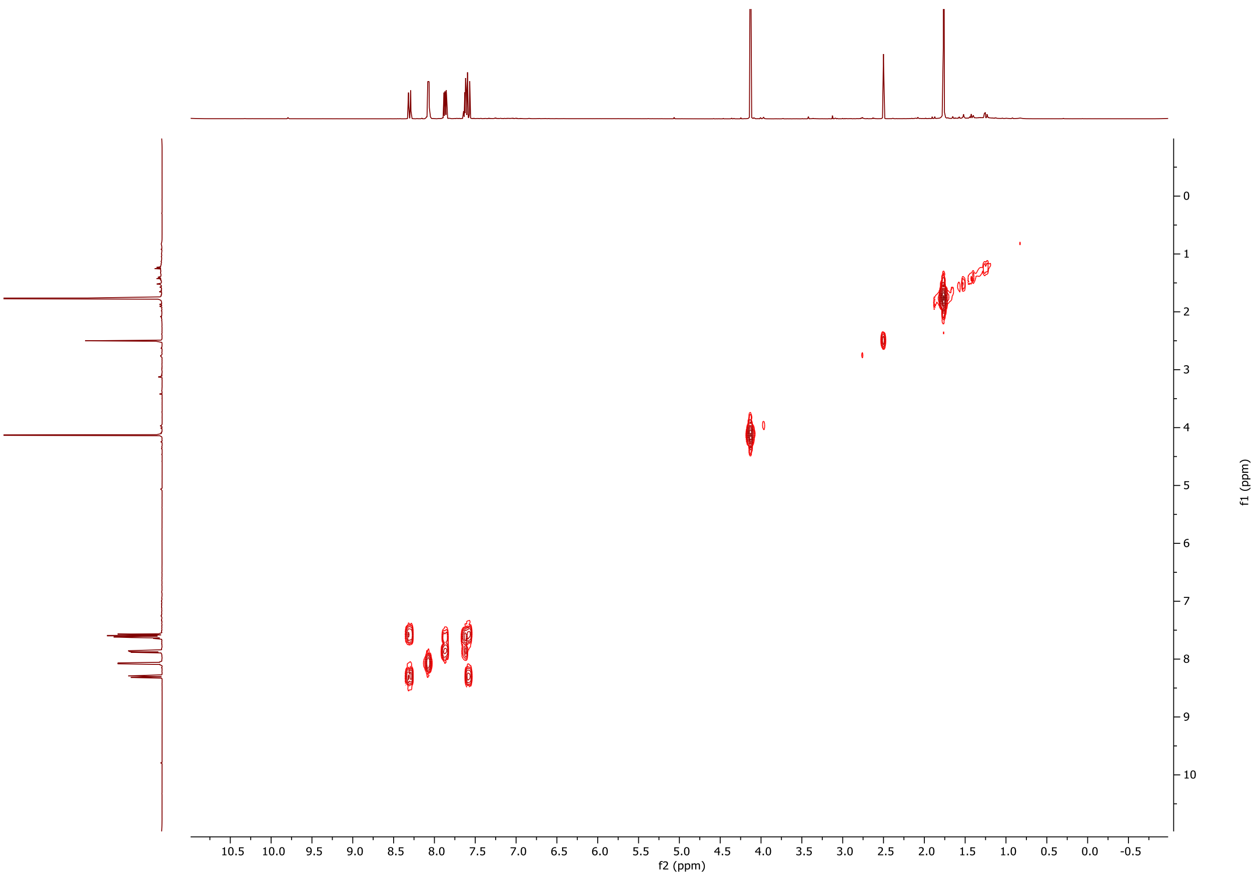

**Figure S5**. $^1$H-$^1$H COSY NMR spectrum of **MeroFluor-2F** in DMSO-*d6*

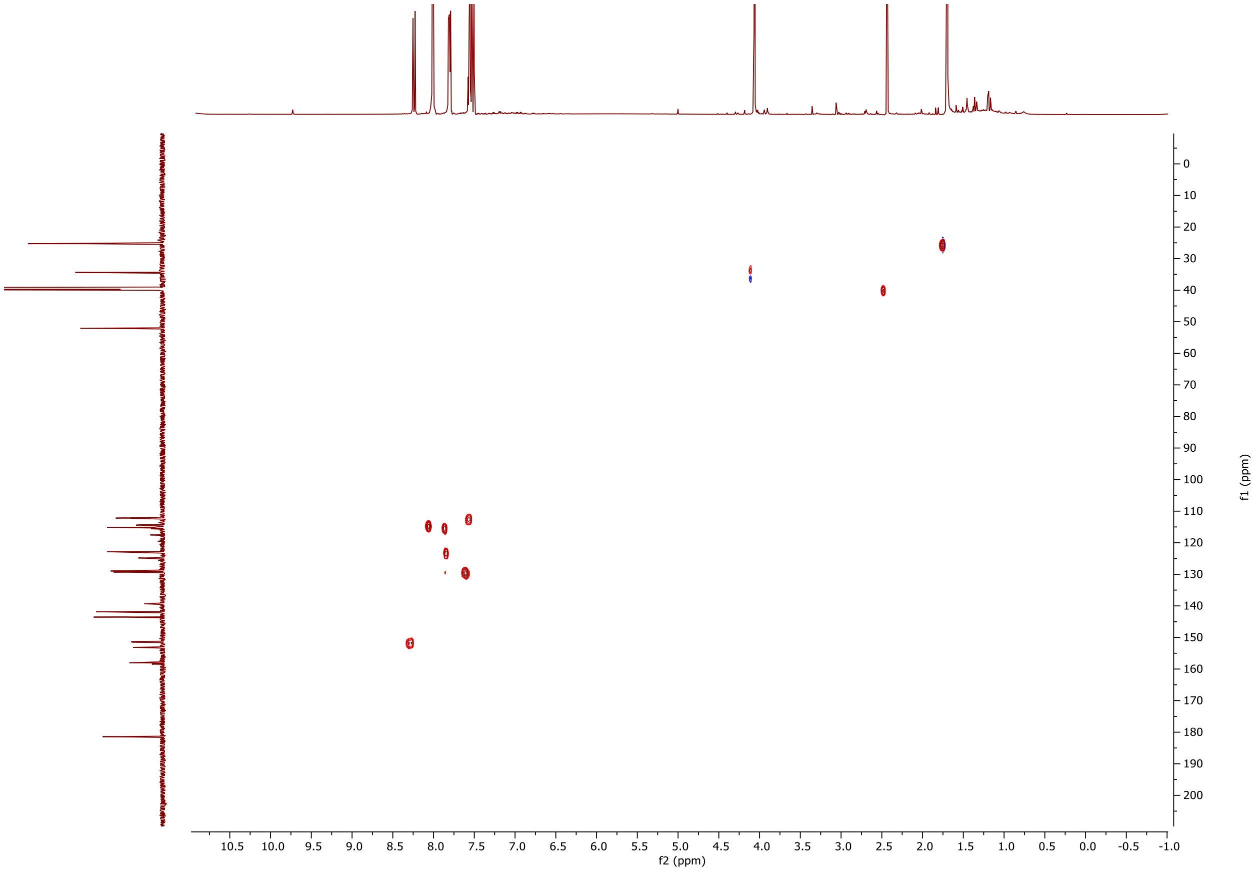

**Figure S6**. $^{1}H$-$^{13}C$ HSQC NMR spectrum of **MeroFluor-2F** in DMSO-*d6*

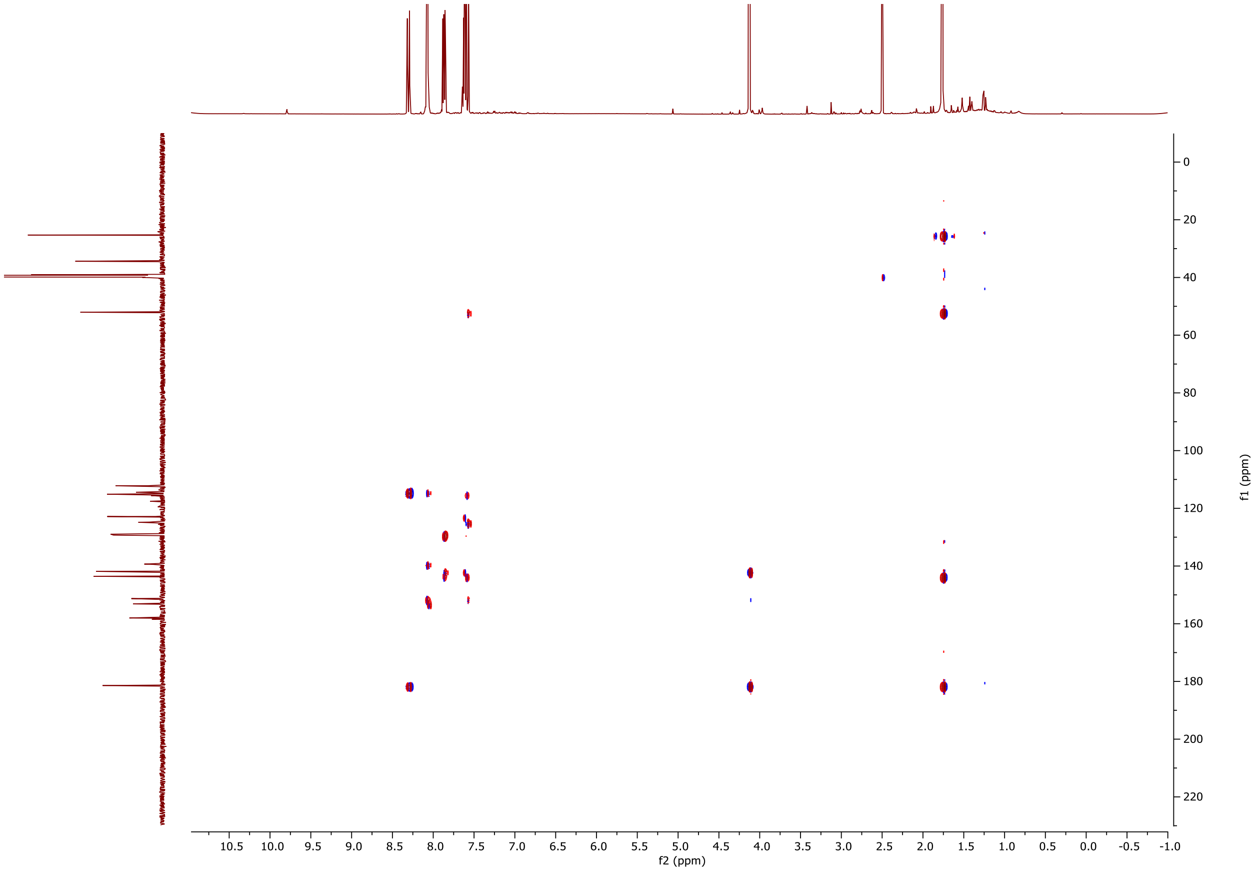

**Figure S7**. $^{1}$H-$^{13}$C HMBC NMR spectrum of **MeroFluor-2F** in DMSO-*d6*

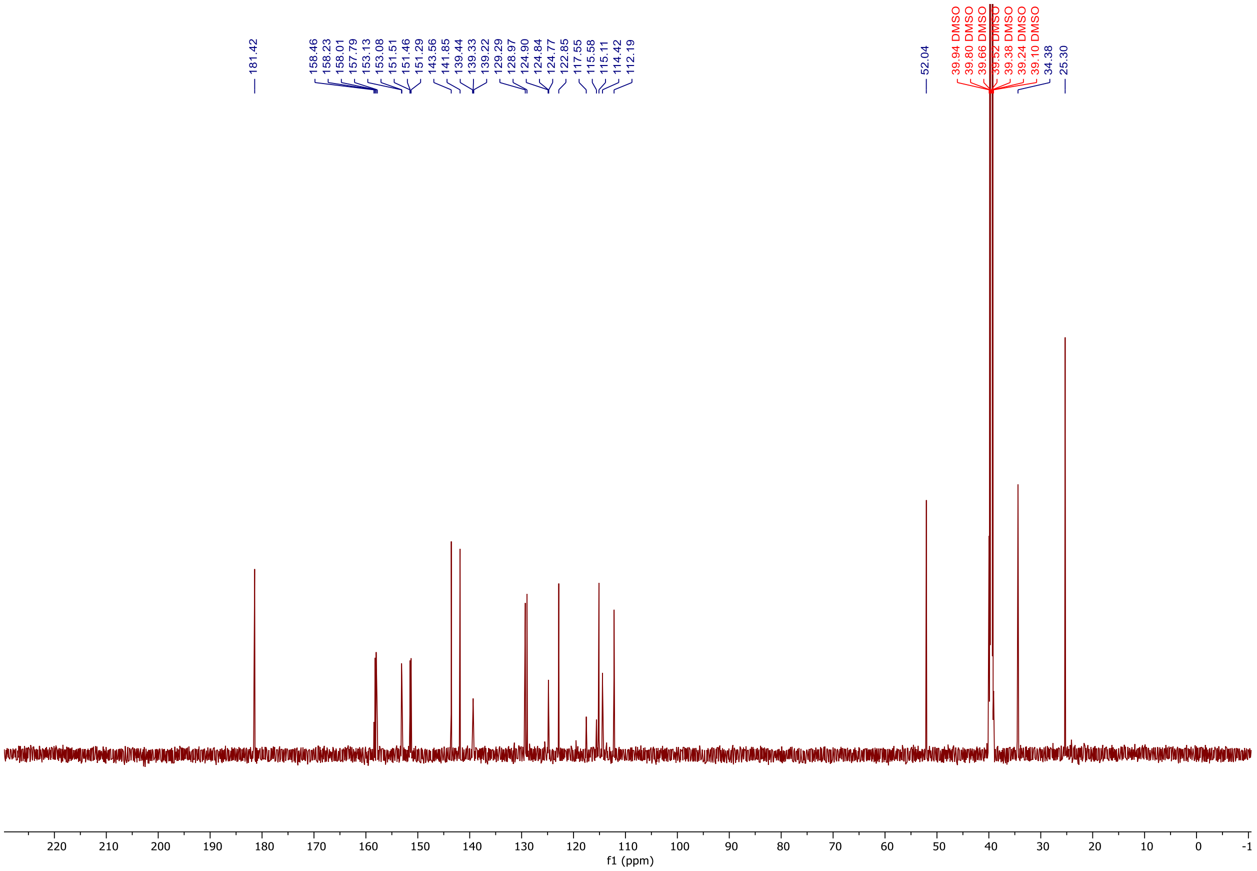

**Figure S8**. $^{13}C$ NMR spectrum of **MeroFluor-2F** in DMSO-*d6*

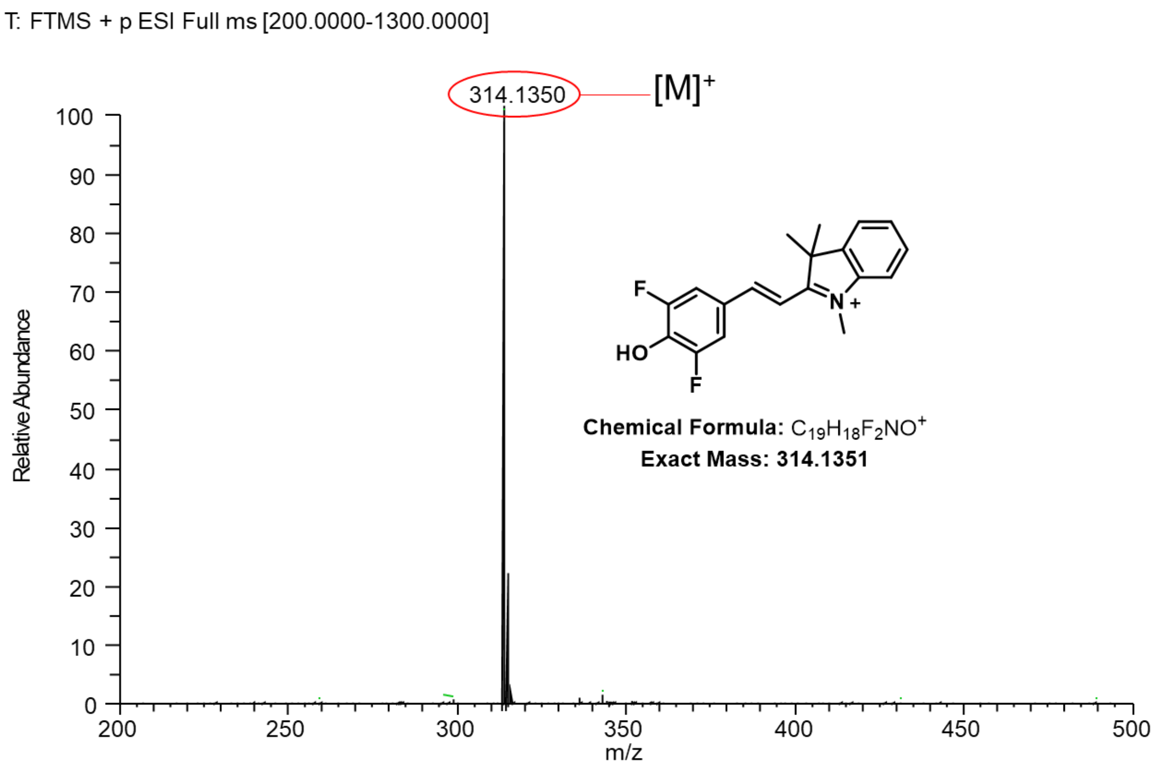

**Figure S9**. HRMS spectrum of compound **MeroFluor-2F**

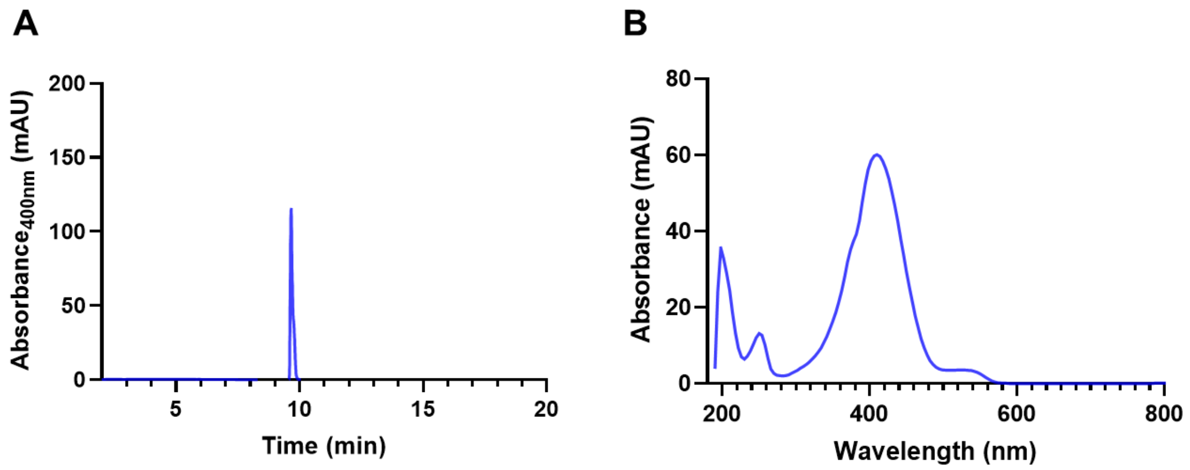

**Figure S10**. (**A**) HPLC trace of **MeroFluor-2F** (retention time, 9.66 min). (**B**) UV-VIS spectrum of **MeroFluor-2F** ($\lambda_{max}$, 409.62 nm). HPLC Method: 2-100%, 0-20 min; 100%, 20-22.5 min; MeOH/$H_2O$ with 0.1% Formic Acid.

### B. Compound 2

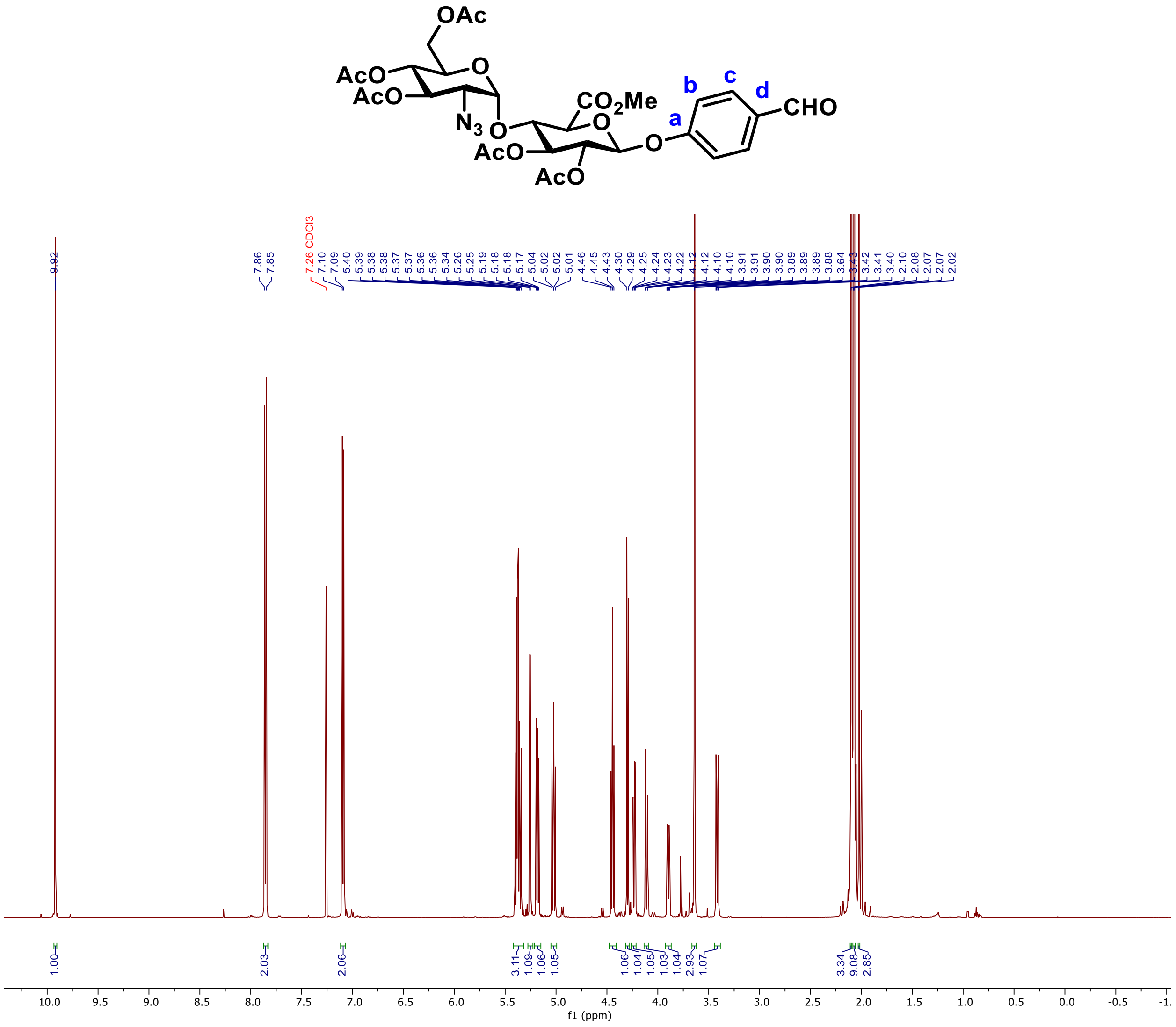

**Figure S11**. $^1H$ NMR spectrum of compound **2** in $CDCl_3$

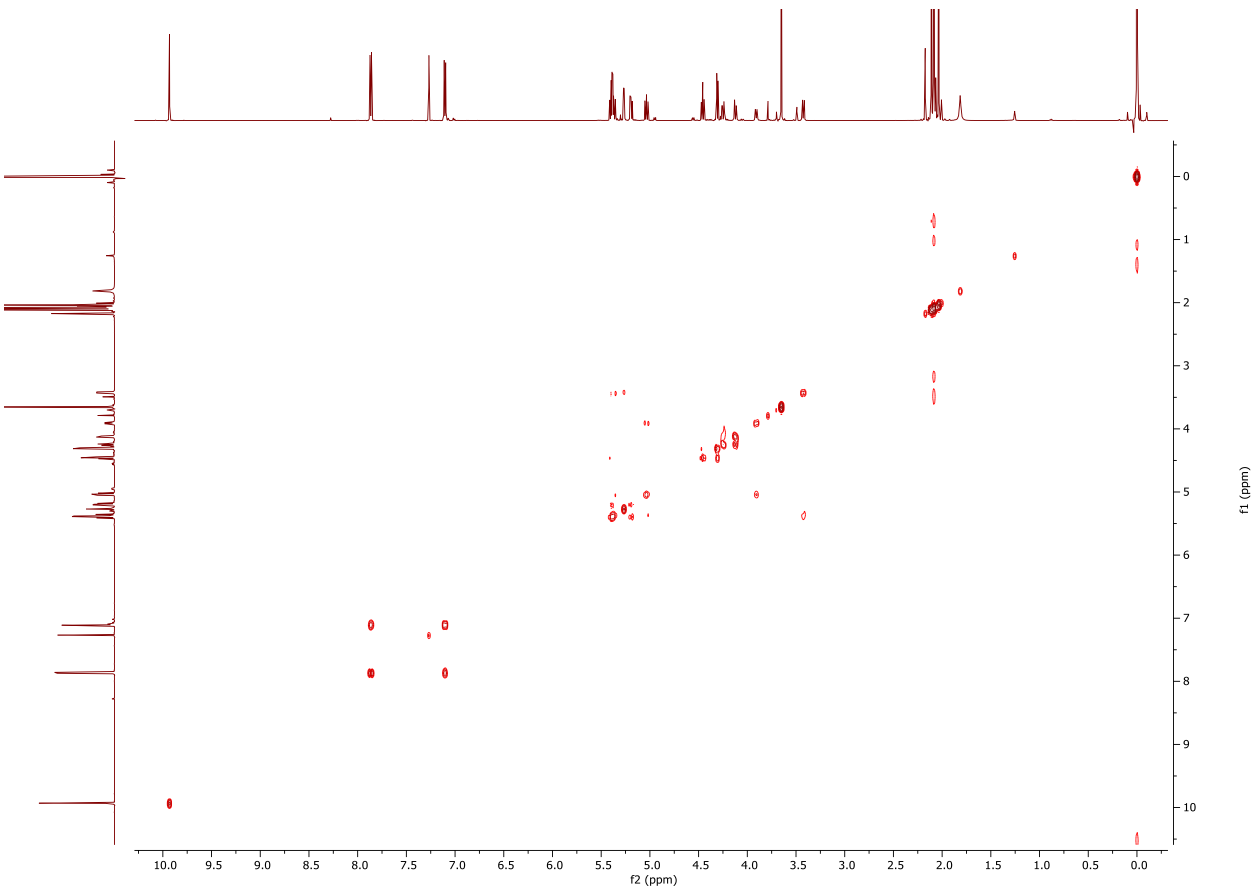

**Figure S12**. $^{1}H$-$^{1}H$ COSY NMR spectrum of compound **2** in $CDCl_3$

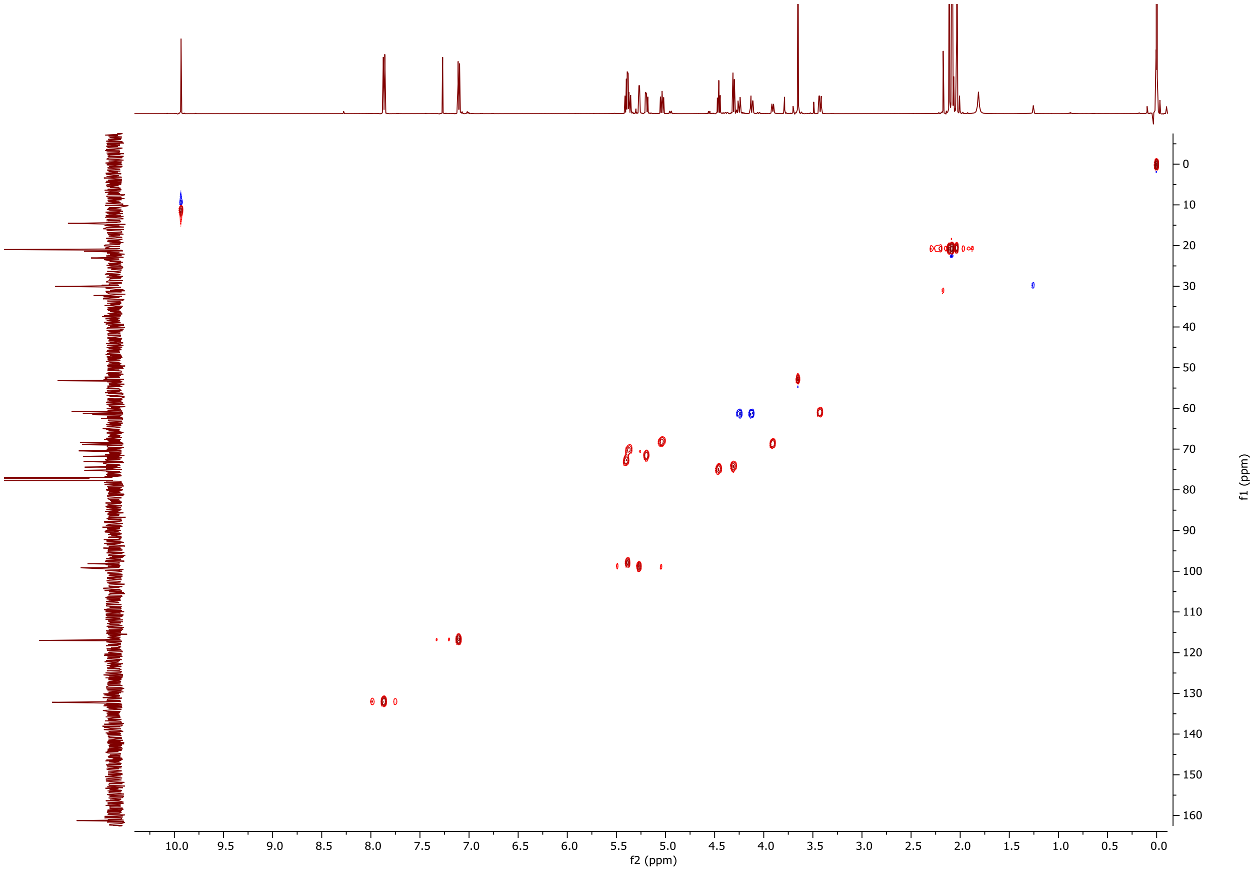

**Figure S13**. $^{1}H$-$^{13}C$ HMBC NMR spectrum of compound **2** in $CDCl_3$

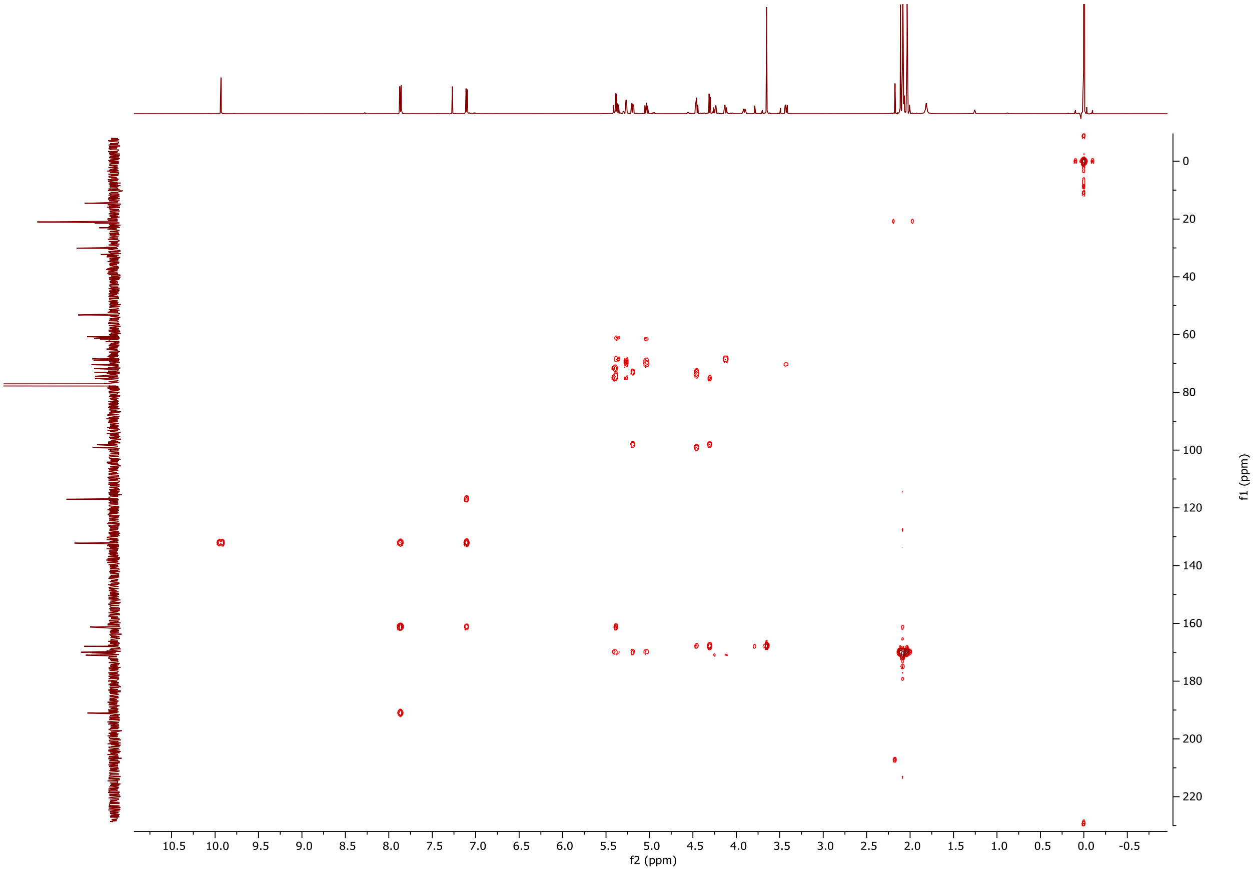

**Figure S14**. $^{1}H$-$^{13}C$ HSQC NMR spectrum of compound **2** in $CDCl_3$

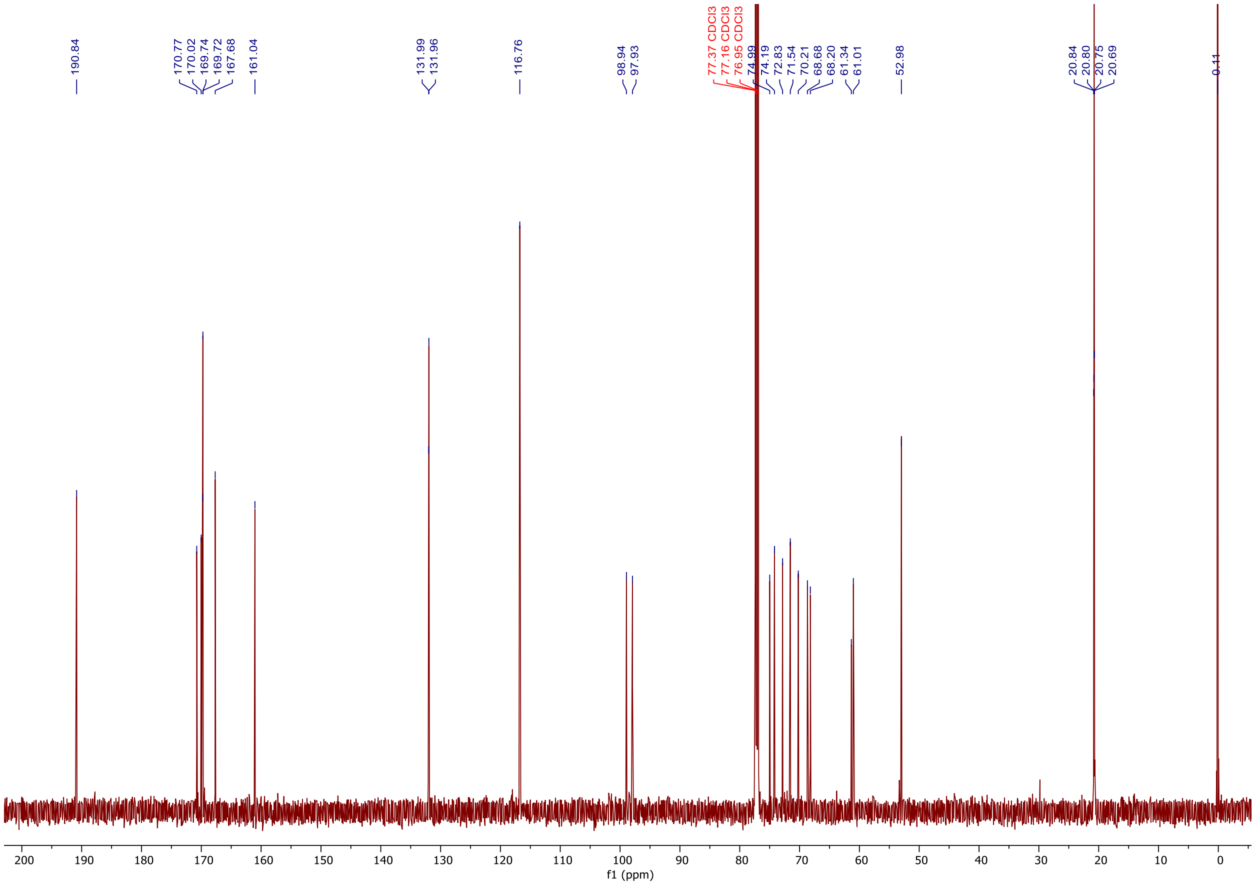

**Figure S15**. $^{13}C$ NMR spectrum of compound **2** in $CDCl_3$

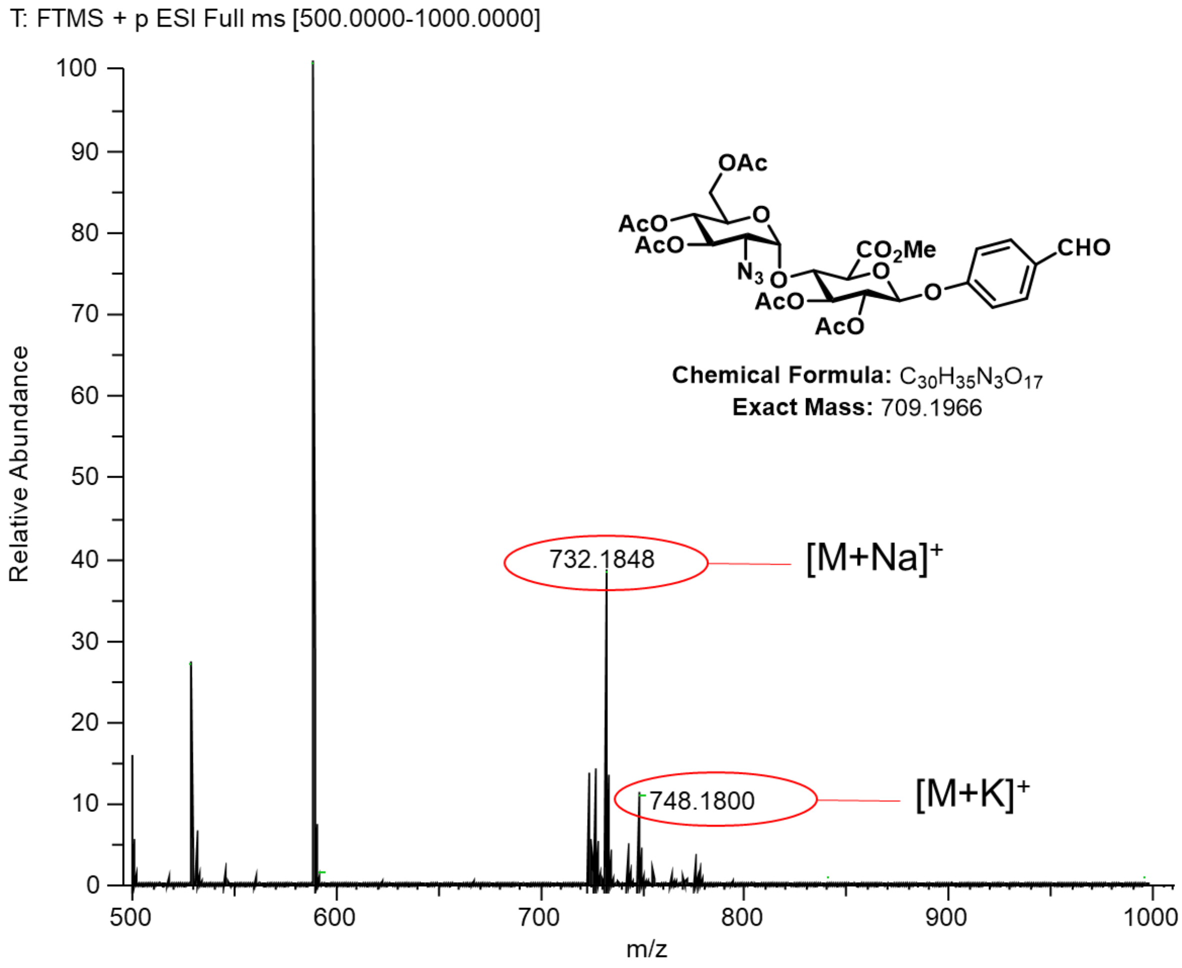

**Figure S16**. HRMS spectrum of compound **2**

1:1 Hexanes: Ethyl Acetate

$R_f$ = 0.39

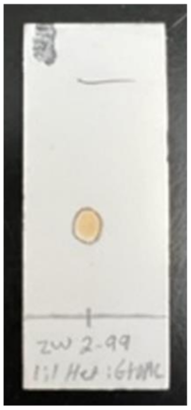

Circle: UV, 254 nm
Brown: 10% $H_2SO_4$ in Methanol

**Figure S17.** TLC of compound **2**

### C. Compound 3

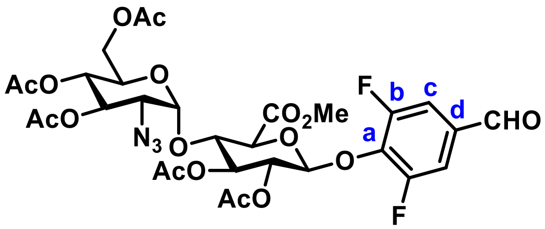

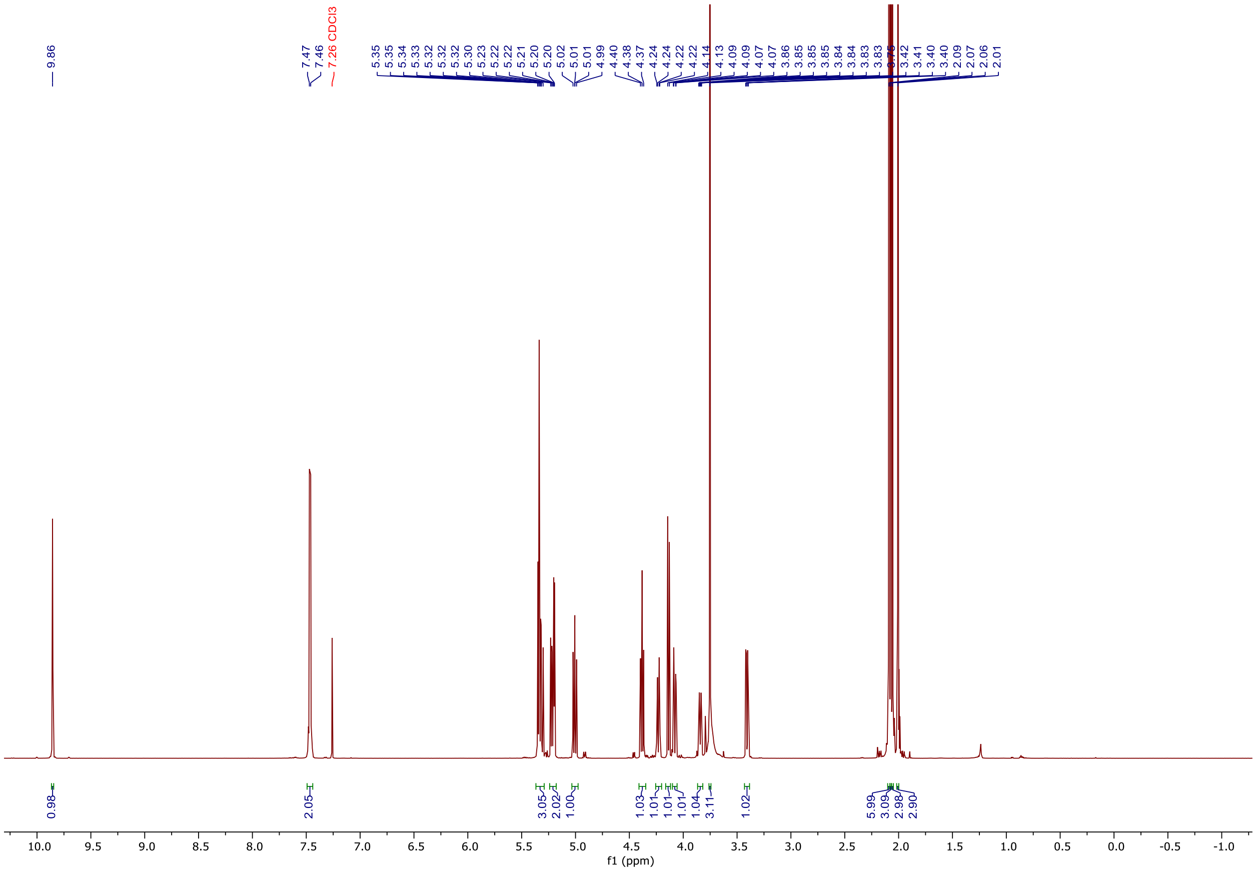

**Figure S18**. $^1$H NMR spectrum of compound **3** in $CDCl_3$

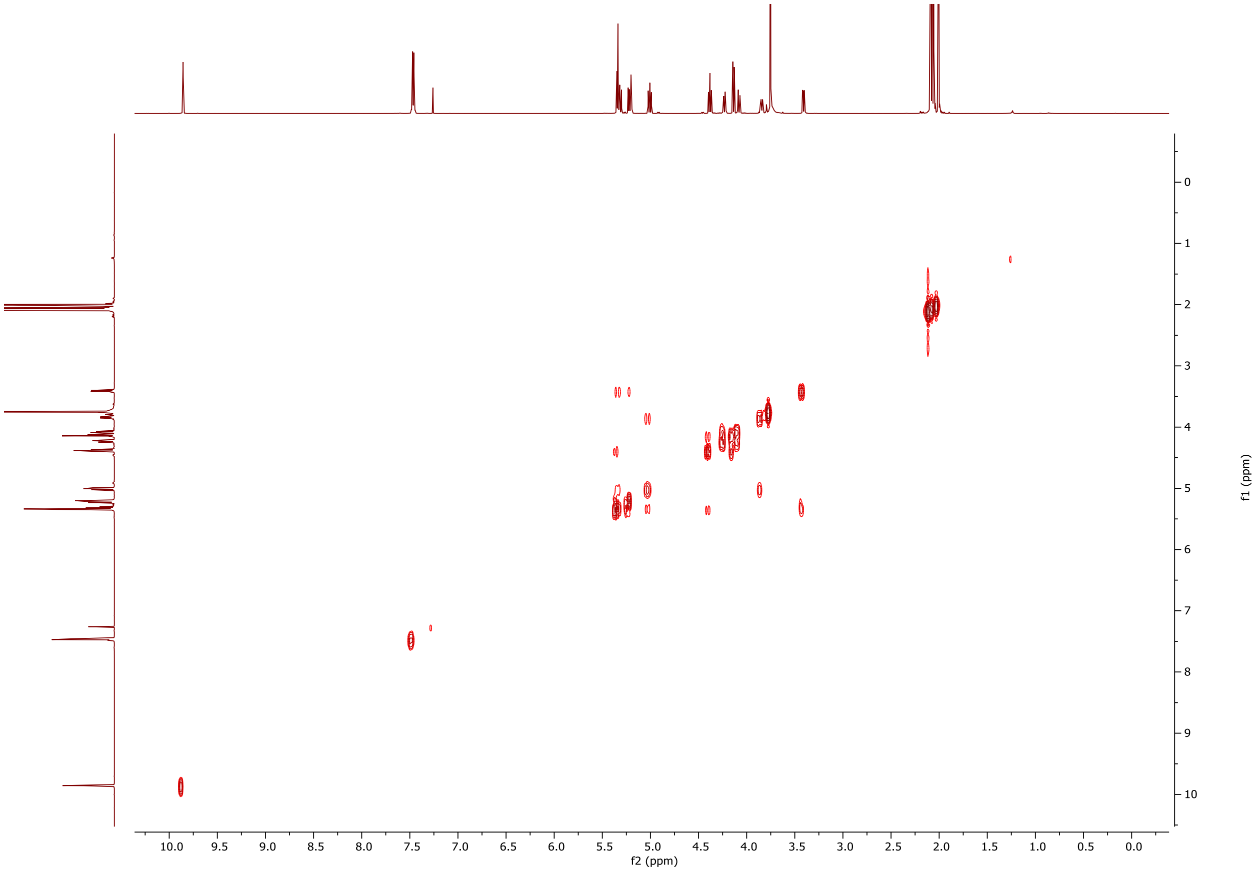

**Figure S19**. $^{1}H$-$^{1}H$ COSY NMR spectrum of compound **3**

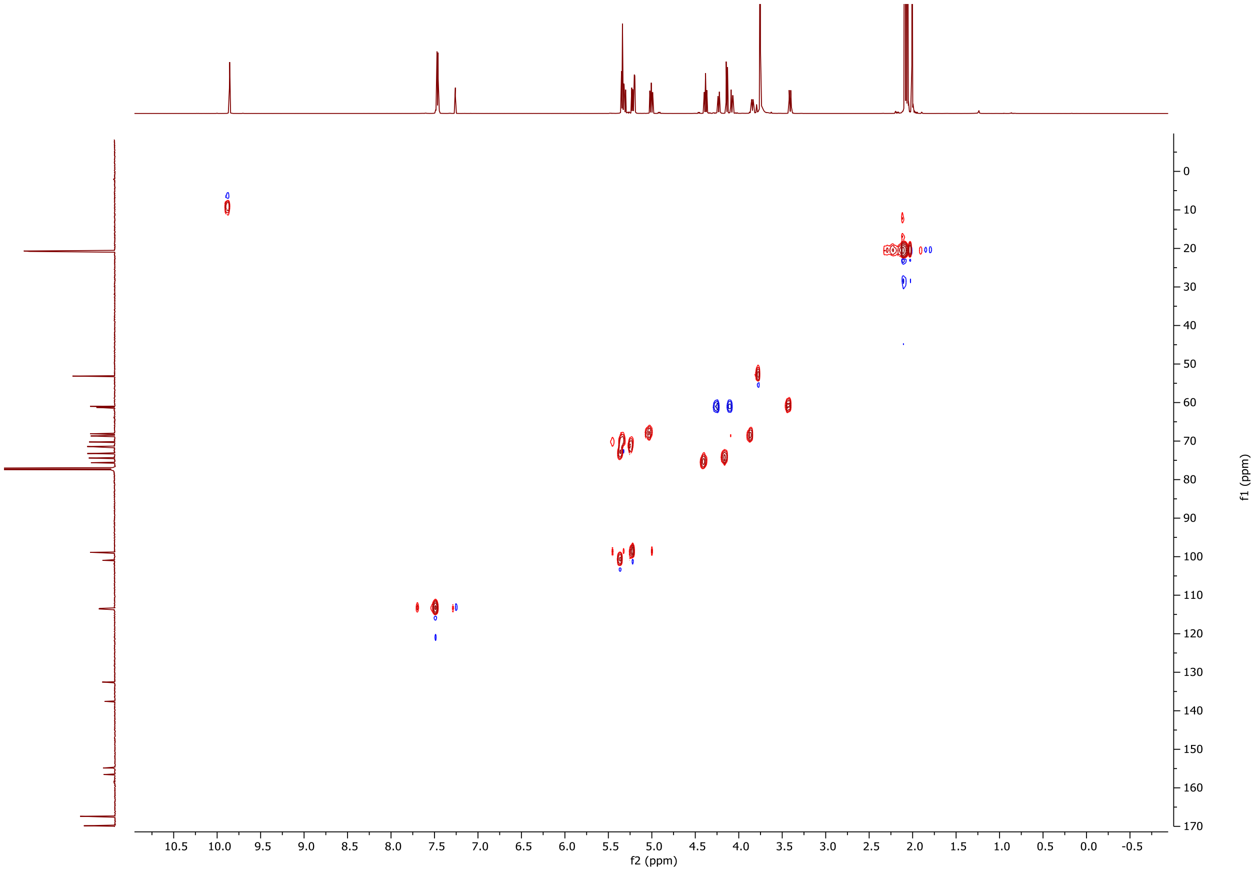

**Figure S20**. $^{1}H$-$^{13}C$ HSQC NMR spectrum of compound **3**

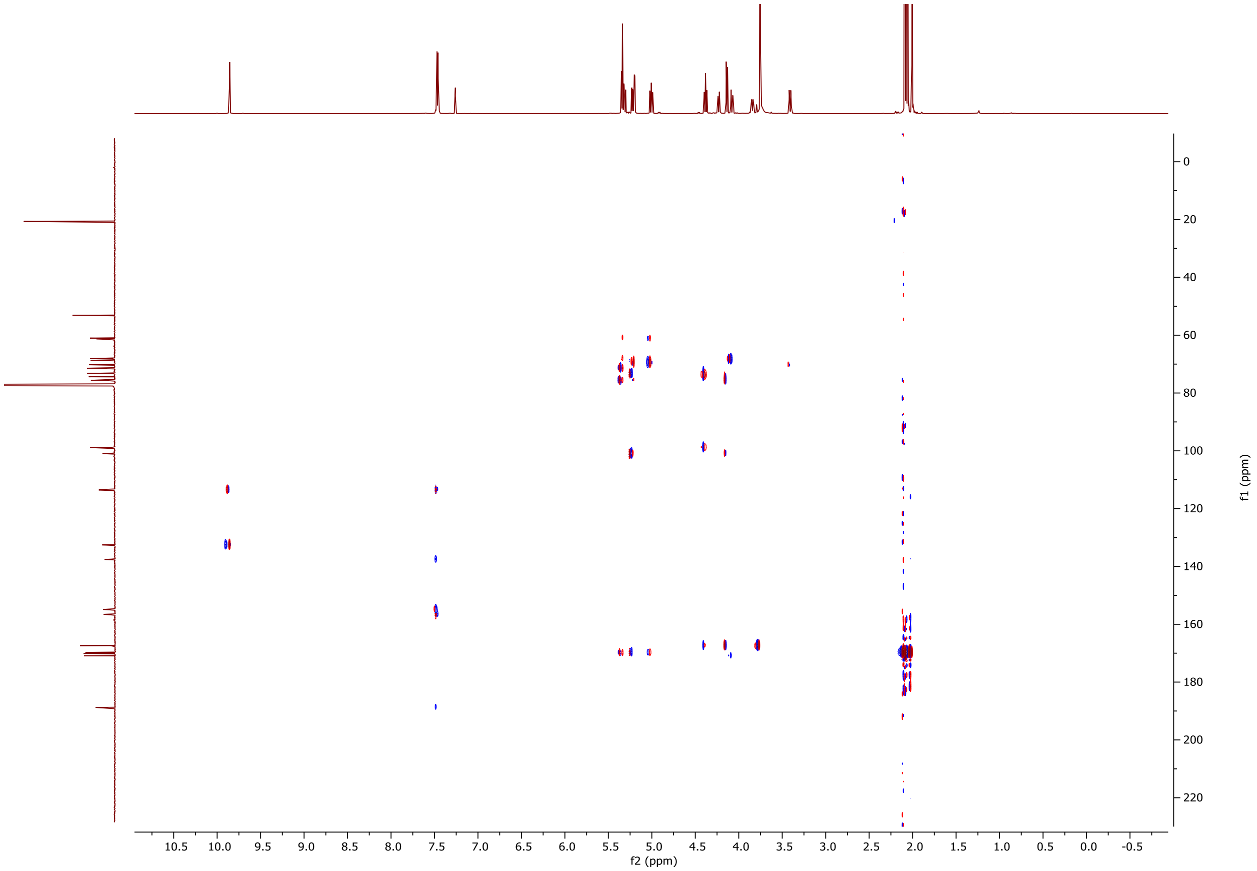

**Figure S21**. [1]H-[13]C HMBC NMR spectrum of compound **3**

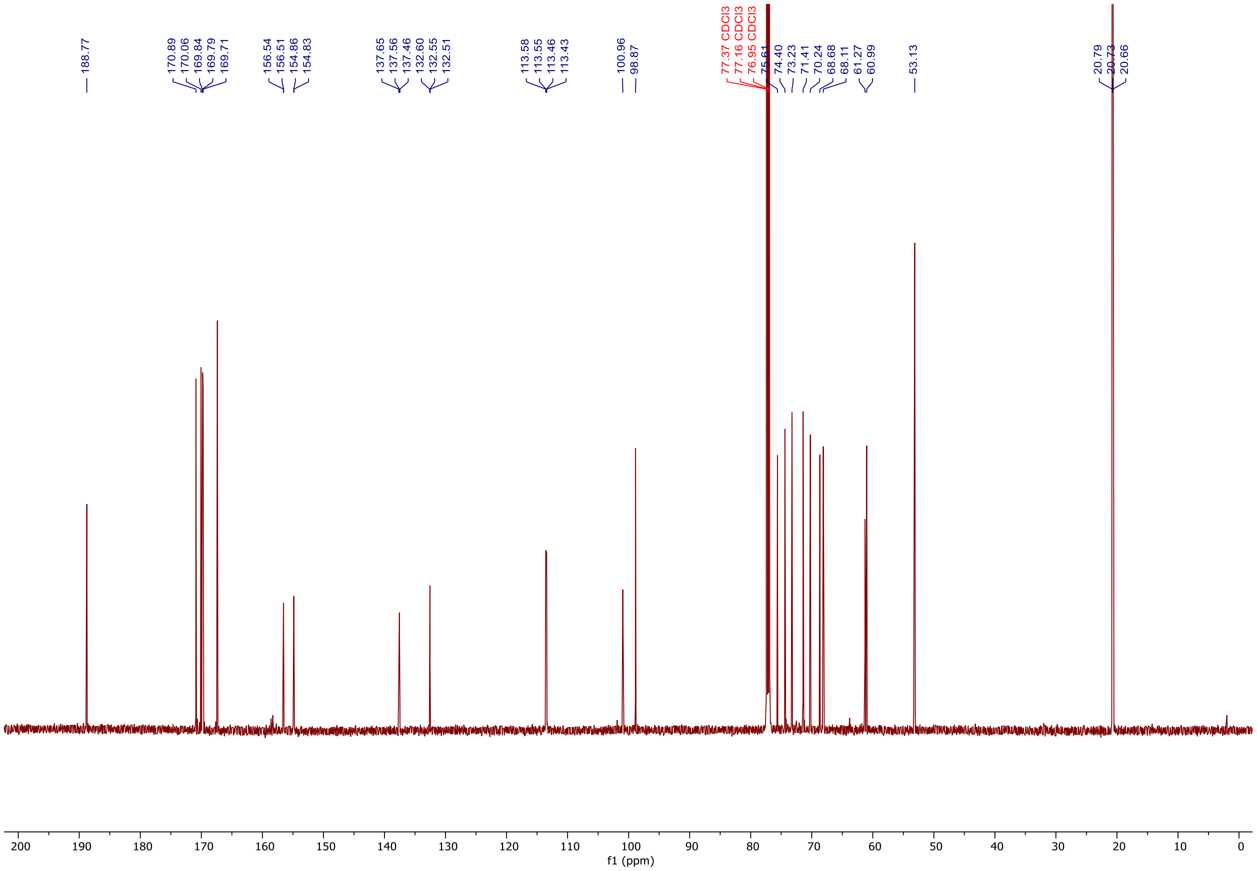

**Figure S22**. $^{13}C$ NMR spectrum of compound **3** in $CDCl_3$

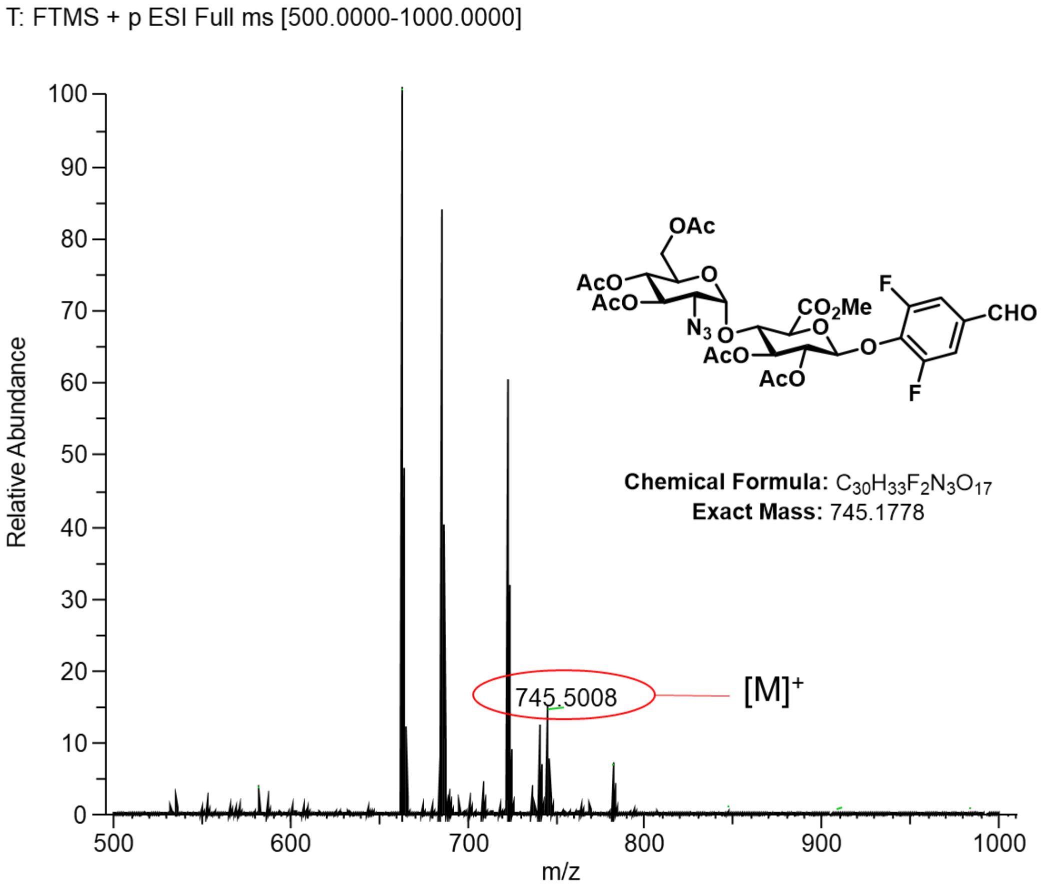

**Figure S23**. HRMS spectrum of compound **3**

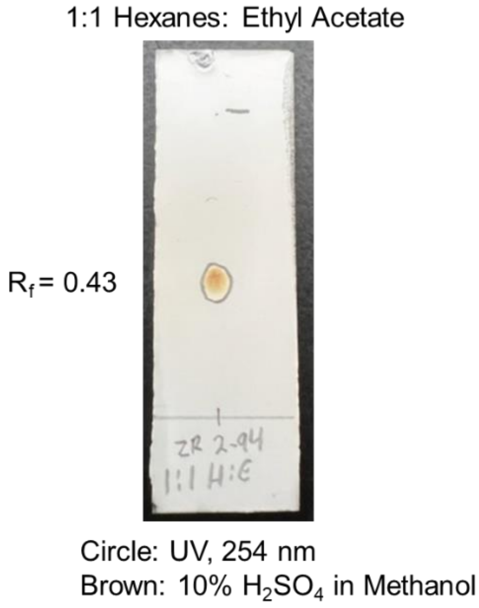

**Figure S24**. TLC of compound **3**

### D. Compound 4

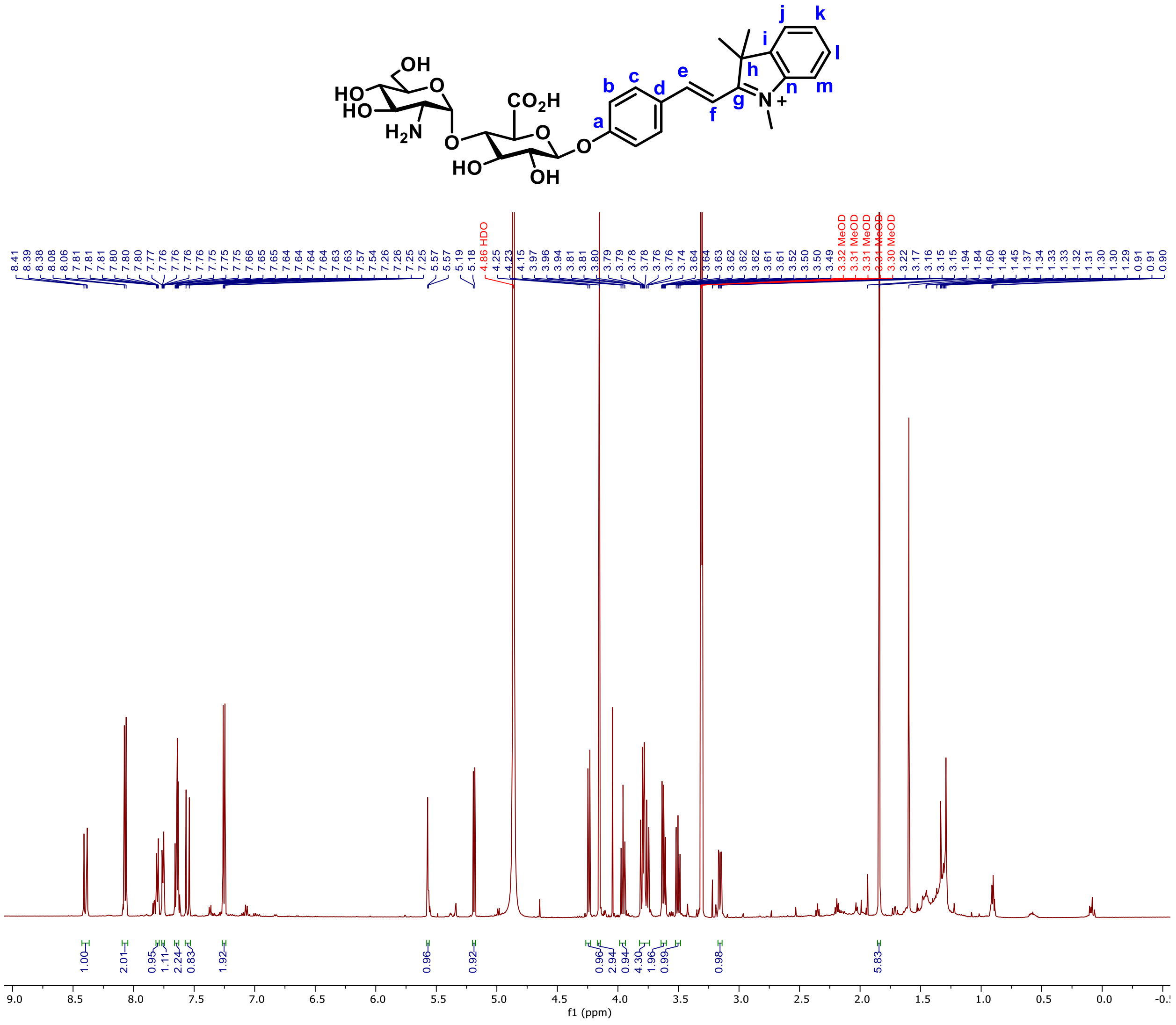

**Figure S25**. $^{1}$H NMR spectrum of compound **4** in MeOD

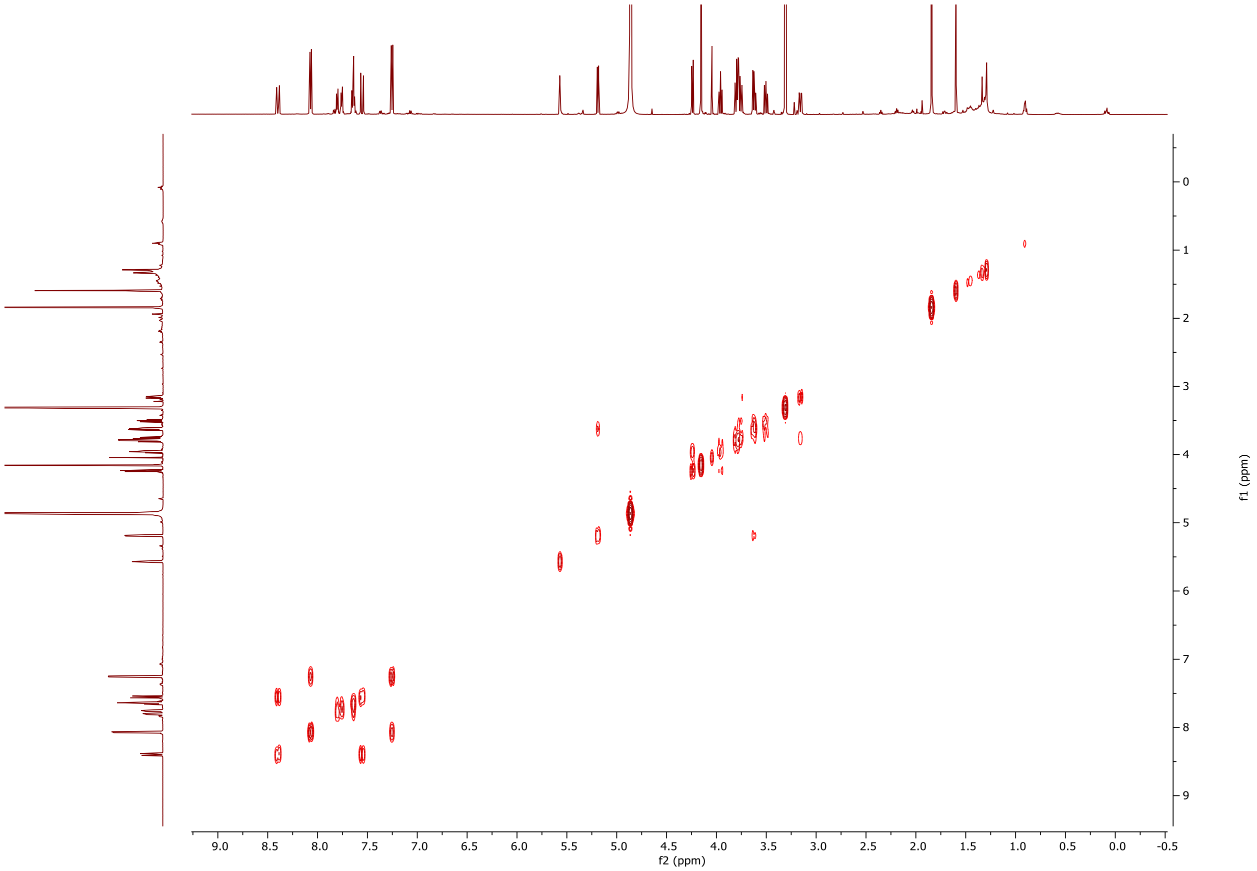

**Figure S26**. $^1H$-$^1H$ COSY NMR spectrum of compound **4** in MeOD

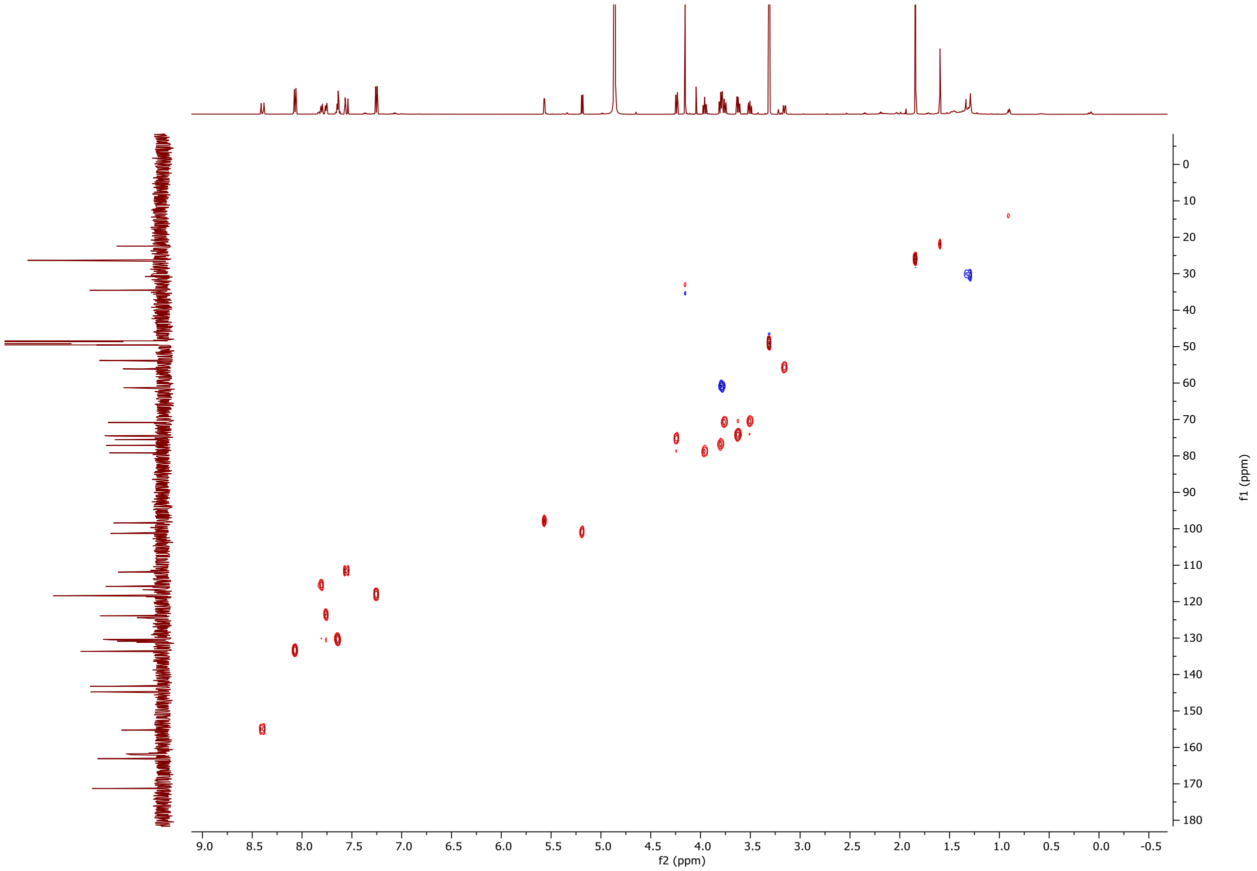

**Figure S27**. $^1H$-$^{13}C$ HSQC NMR spectrum of compound **4** in MeOD

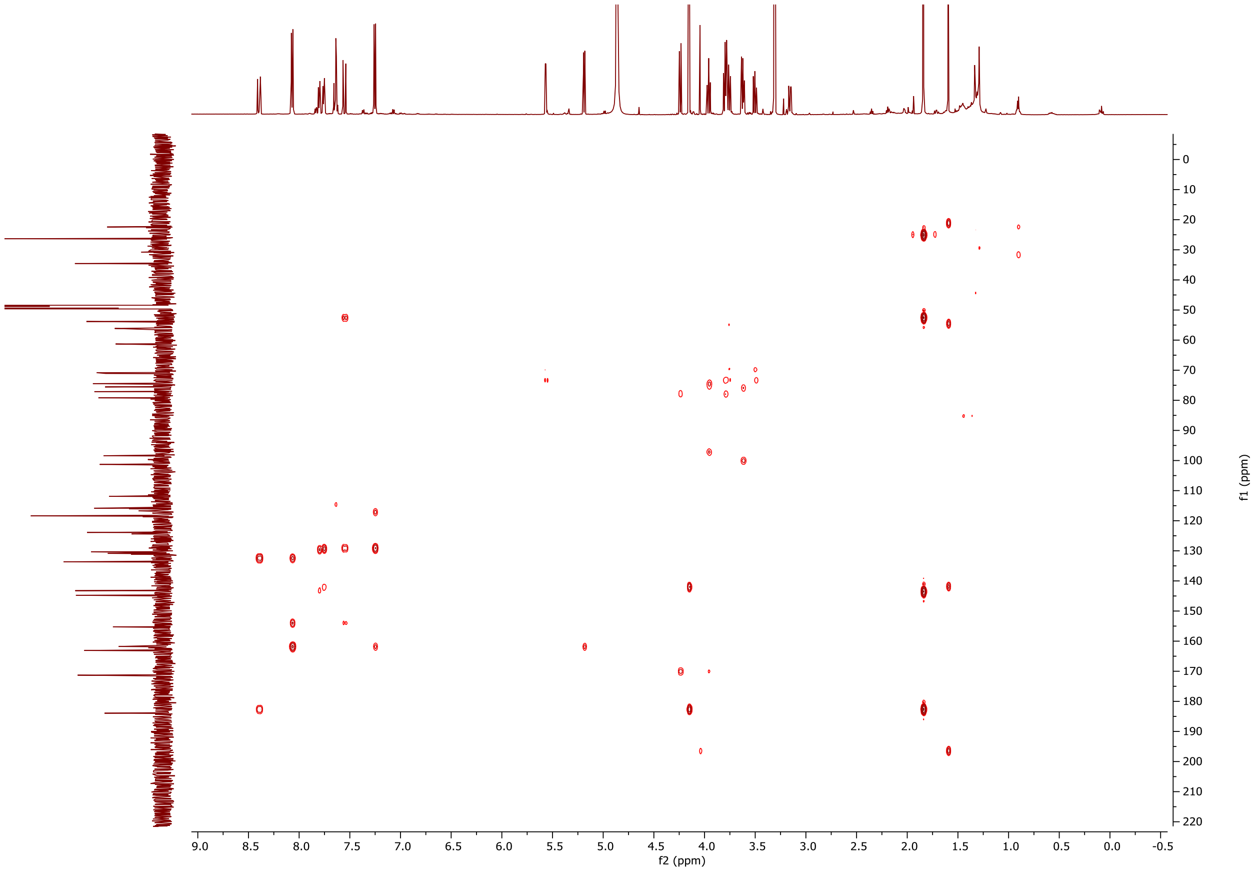

**Figure S28**. $^{1}H$-$^{13}C$ HMBC NMR spectrum of compound **4** in MeOD

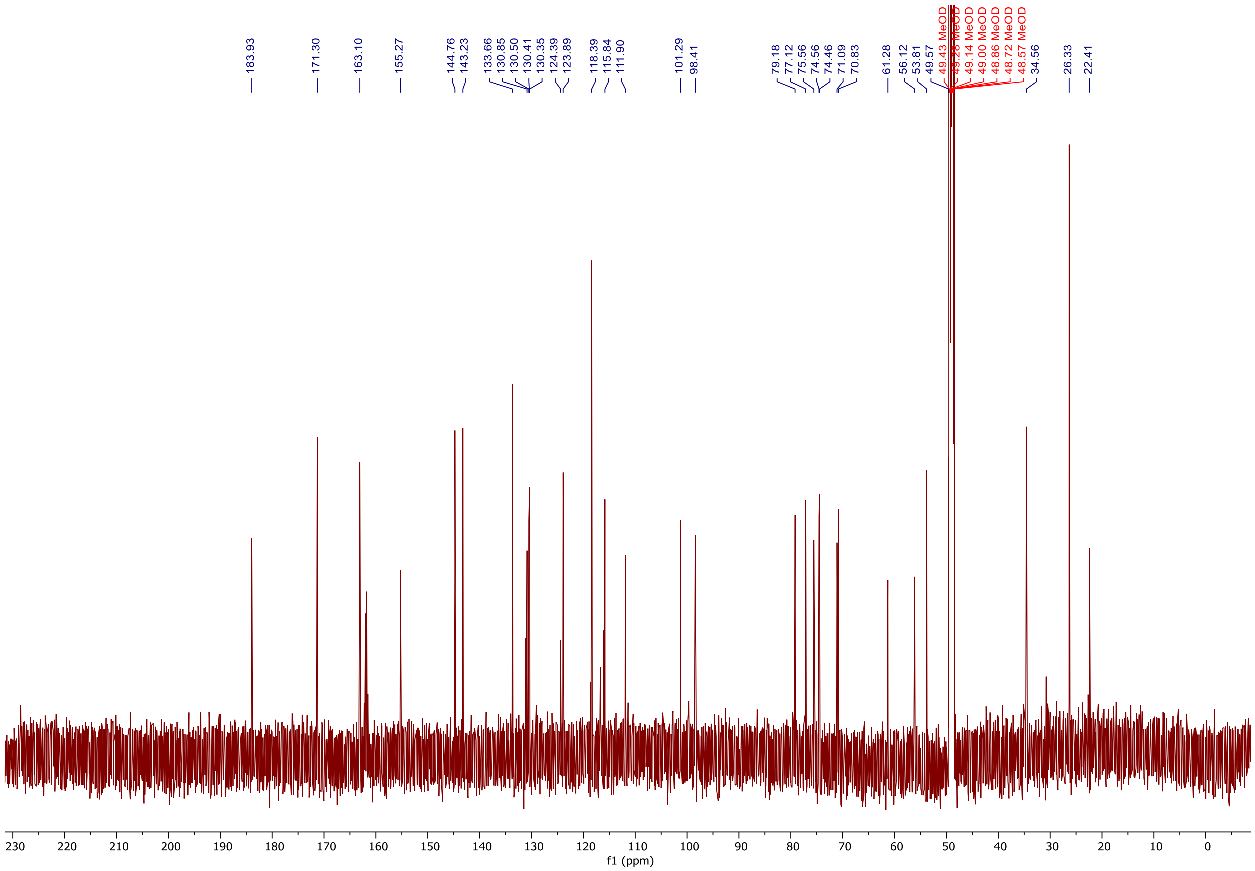

**Figure S29**. $^{13}C$ NMR spectrum of compound **4** in MeOD

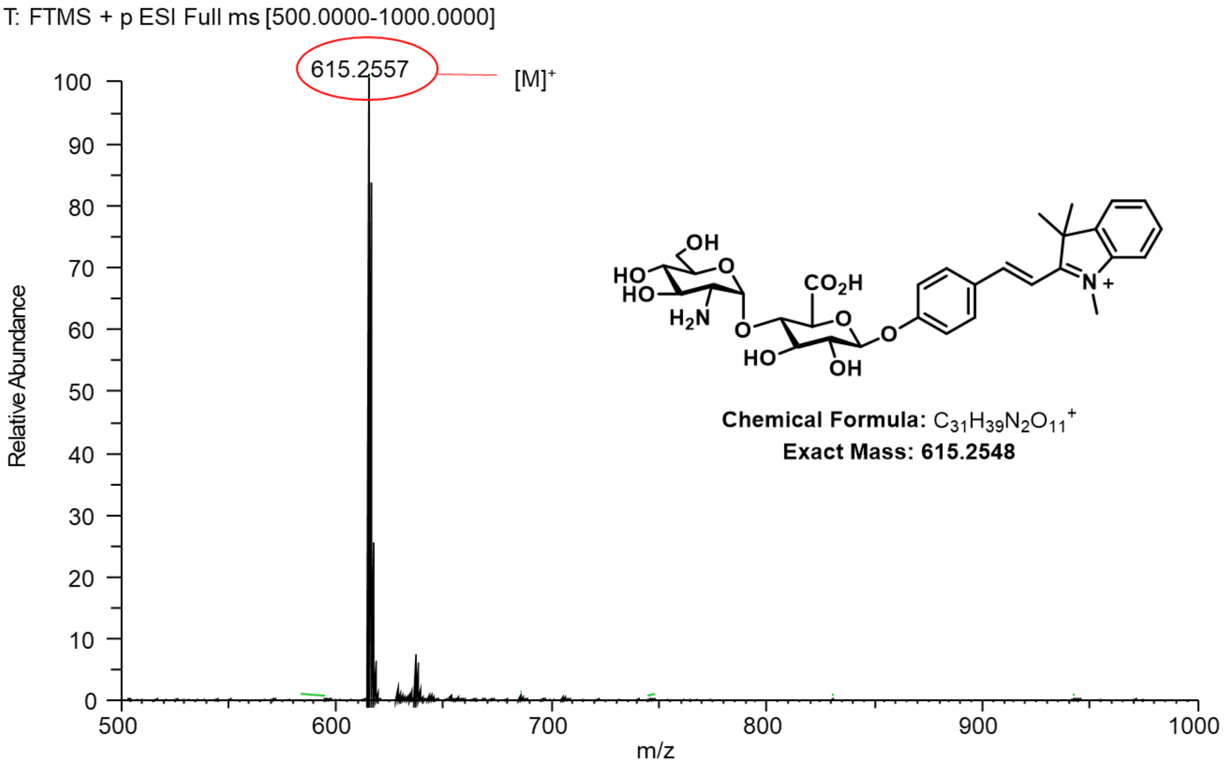

**Figure S30**. HRMS spectrum of compound **4**

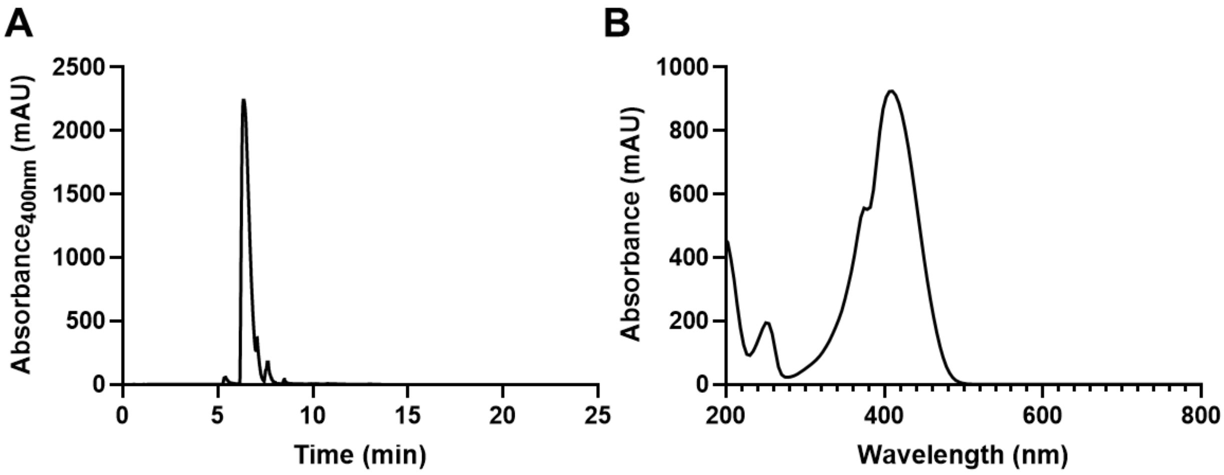

**Figure S31.** (**A**) HPLC trace of compound **4** (retention time, 6.36 min. (**B**) UV-VIS spectrum of compound **4** ($\lambda_{max}$, 407 nm). HPLC Method: 2-100%, 0-20 min; 100%, 20-22.5 min; MeOH/$H_2O$ with 0.1% Formic Acid.

### E. Compound 5

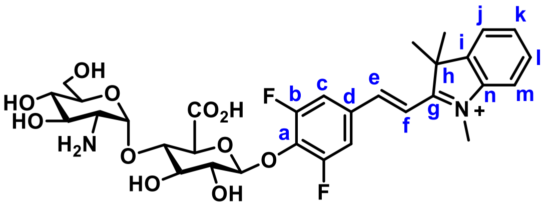

Compound **5**

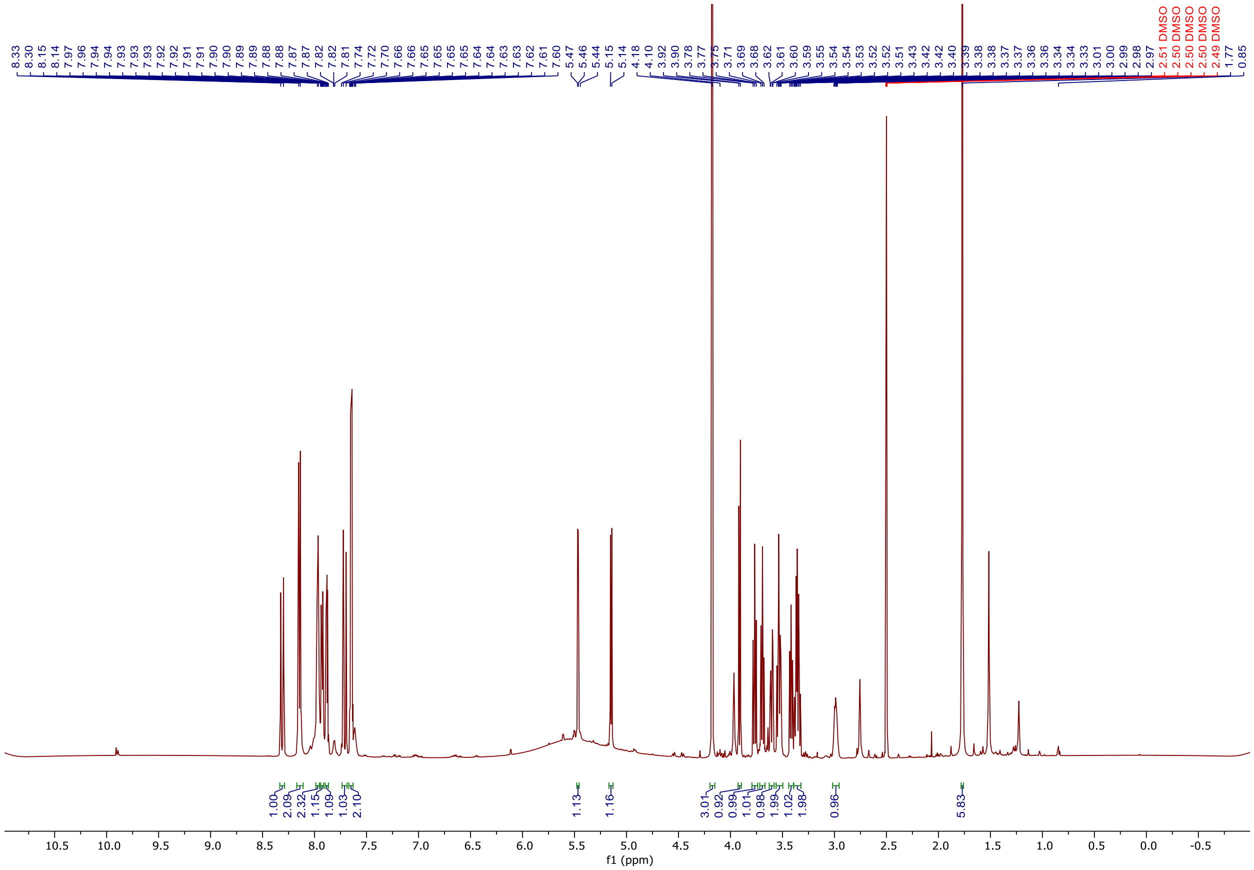

**Figure S32**. $^{1}$H NMR spectrum of compound **5** in DMSO-*d6*

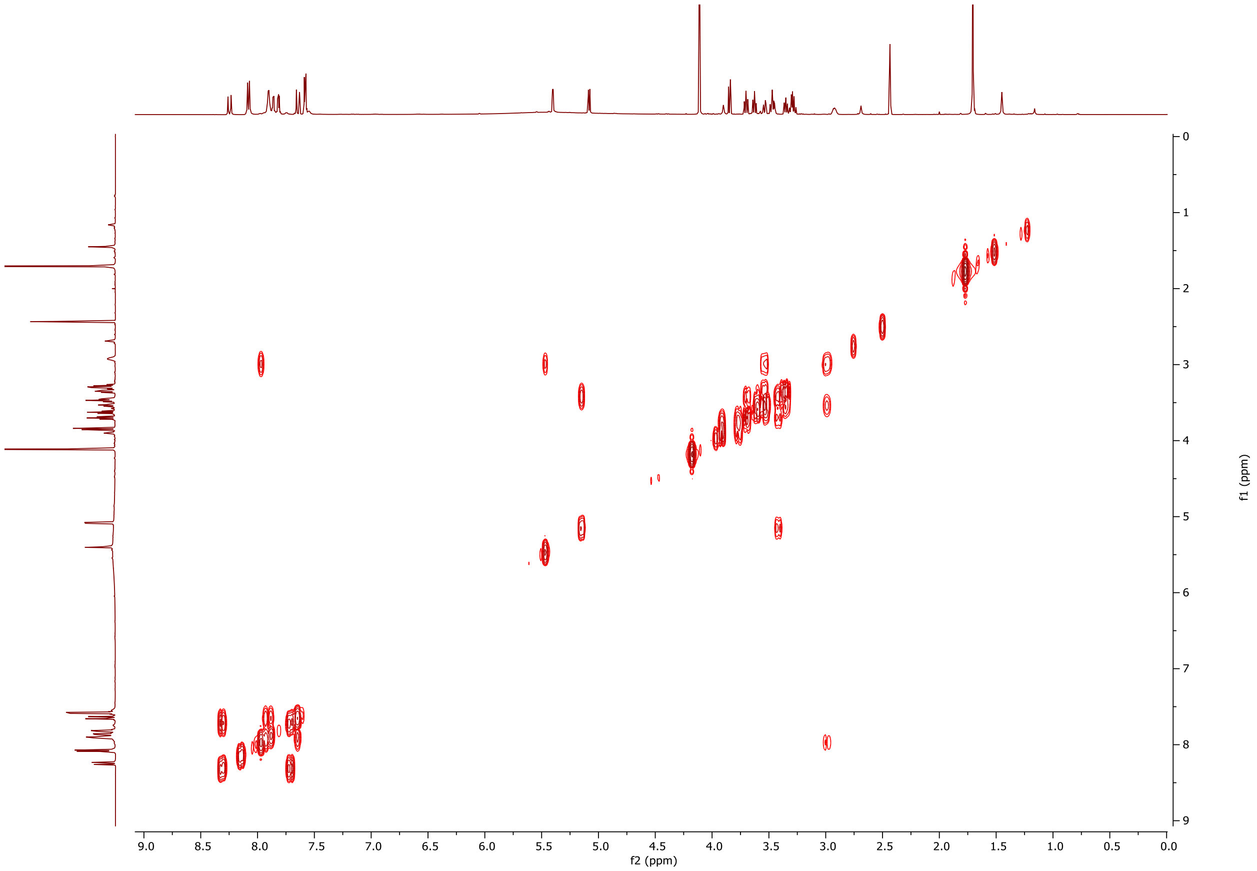

**Figure S33**. $^1H$-$^1H$ COSY NMR spectrum of compound **5** in DMSO-*d6*

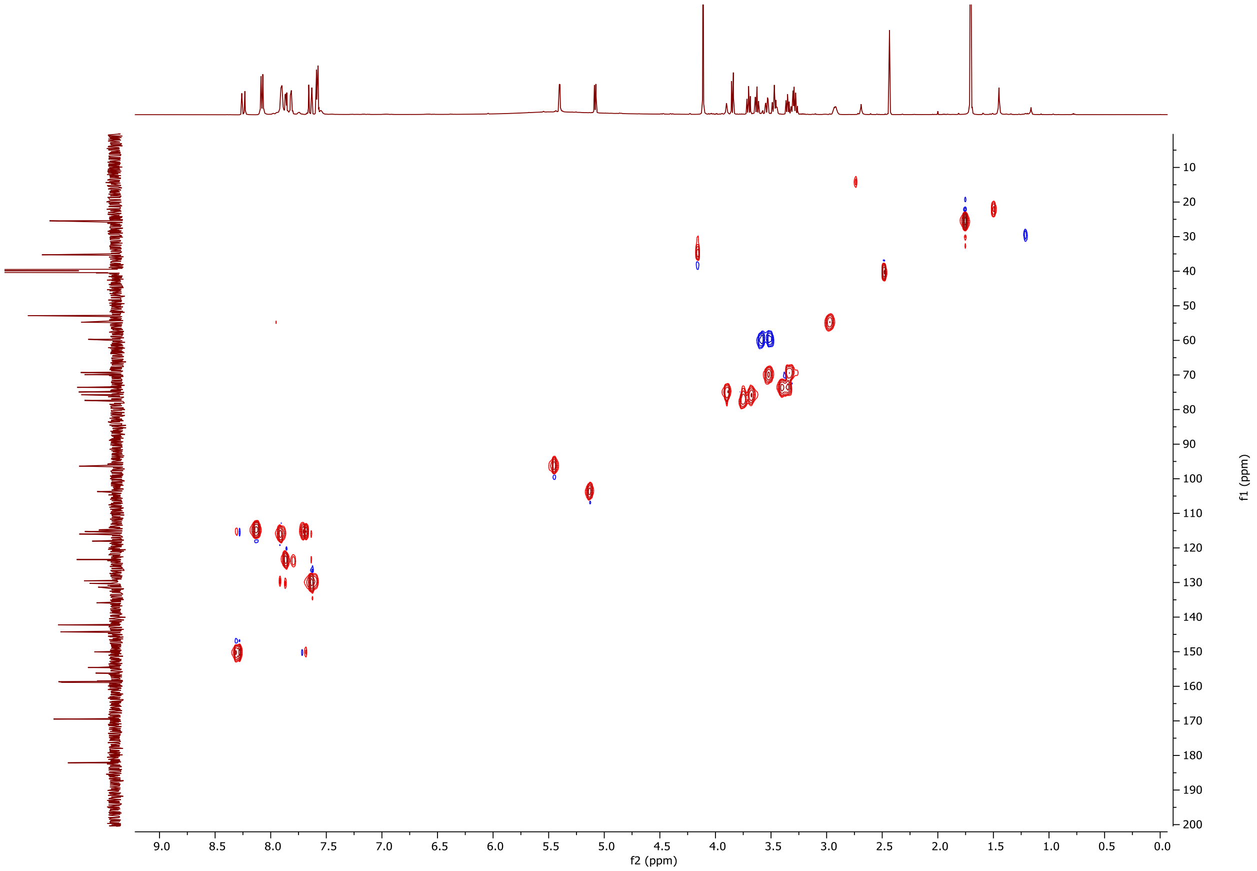

**Figure S34**. $^{1}H$-$^{13}C$ HSQC NMR spectrum of compound **5** in DMSO-*d6*

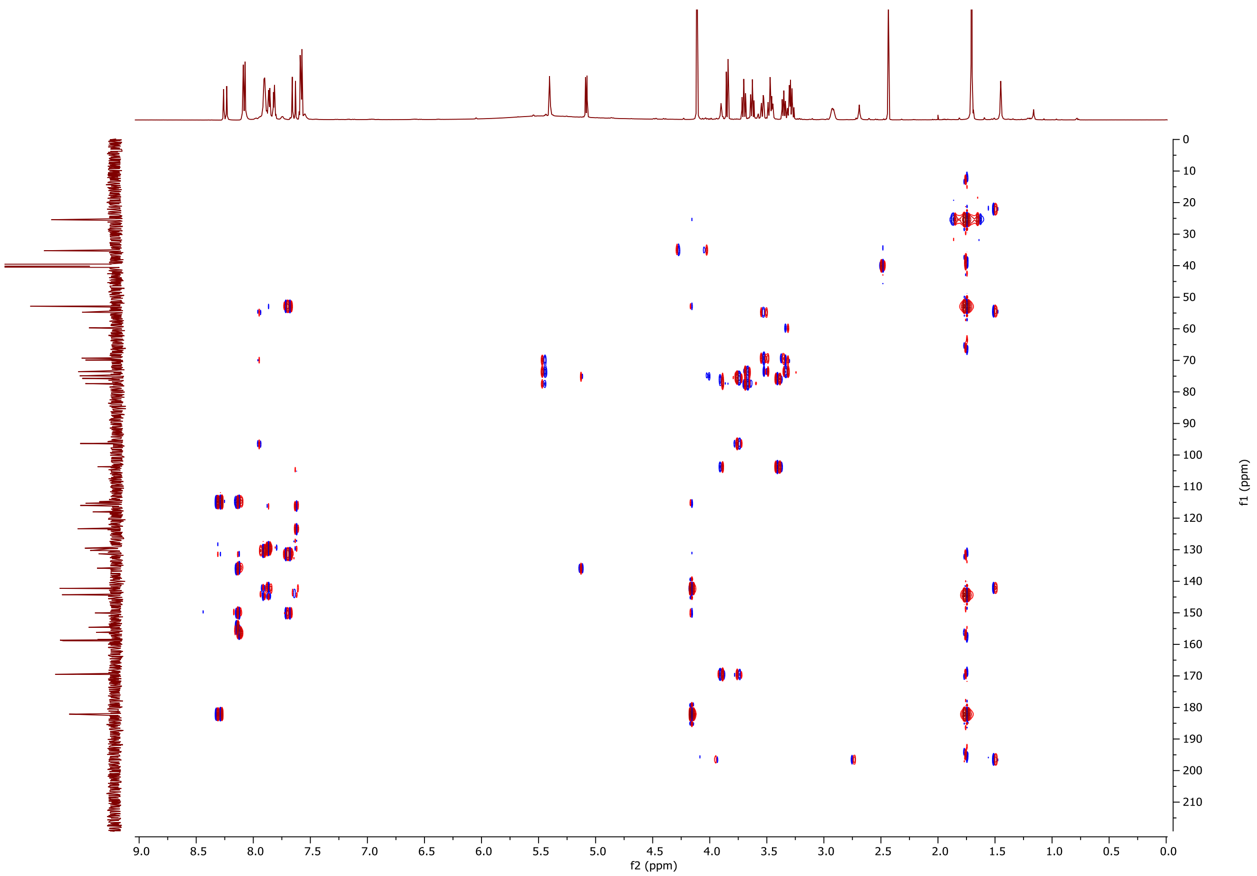

**Figure S35**. $^{1}$H-$^{13}$C HMBC NMR spectrum of compound **5** in DMSO-*d6*

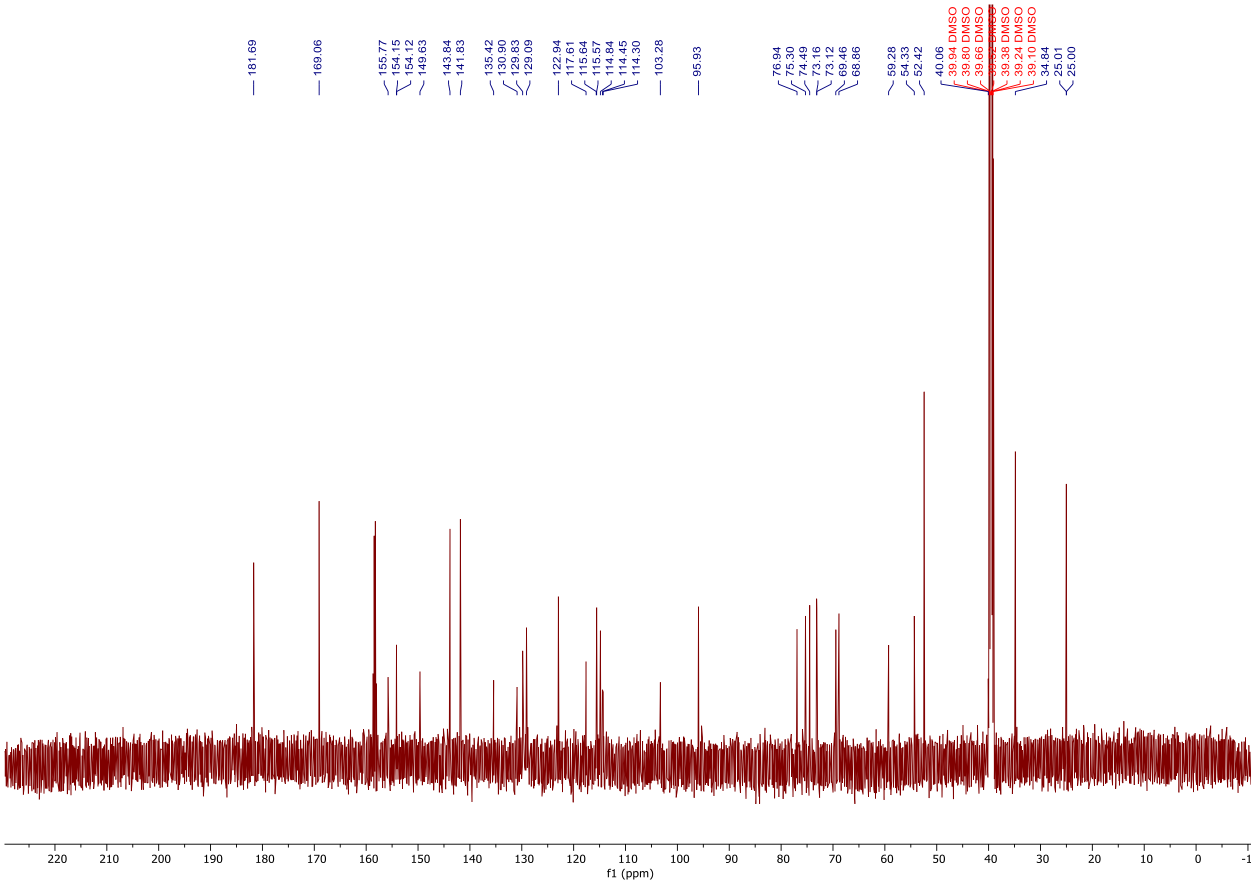

**Figure S36**. $^{13}C$ NMR spectrum of compound **5** in DMSO-*d6*

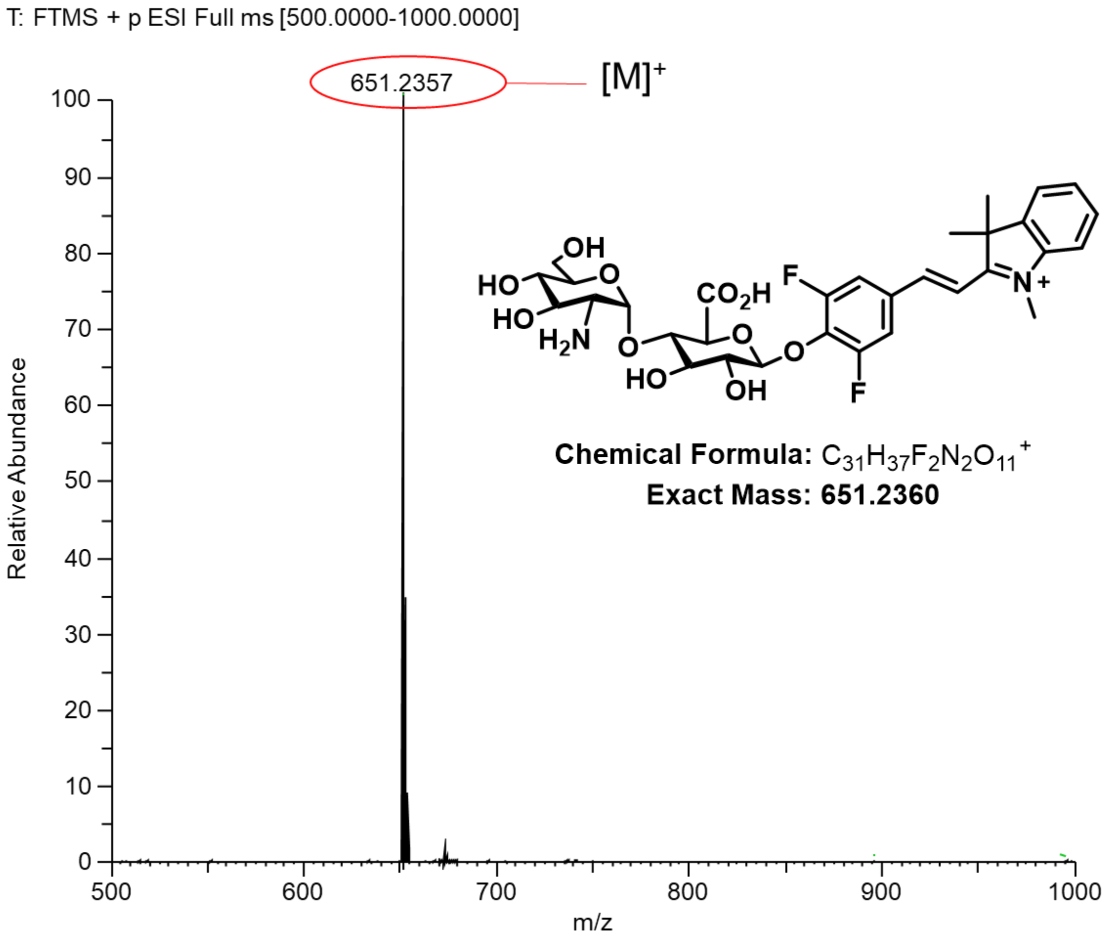

**Figure S37**. HRMS spectrum of compound **5**

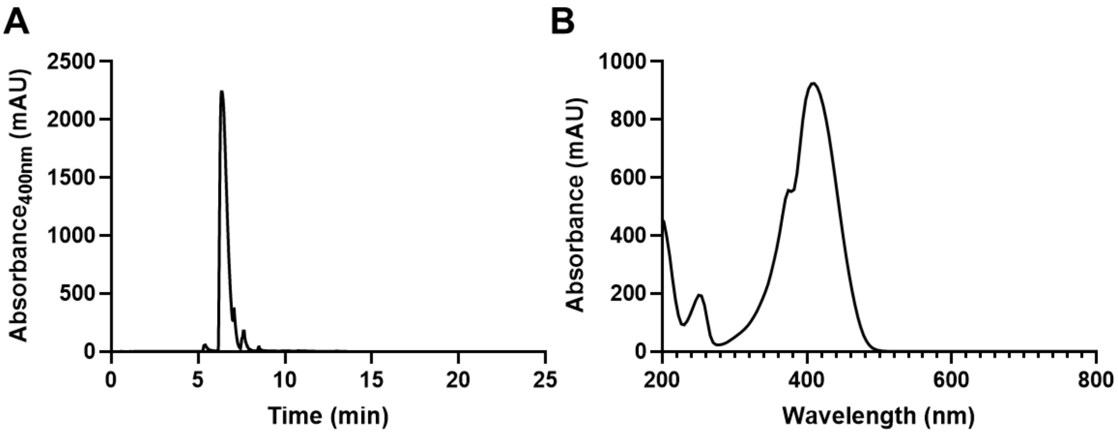

**Figure S38.** (**A**) HPLC trace of compound **4** (retention time, 6.36 min. (**B**) UV-VIS spectrum of compound **4** ($\lambda_{max}$, 407 nm). HPLC Method: 2-100%, 0-20 min; 100%, 20-22.5 min; MeOH/$H_2O$ with 0.1% Formic Acid.

### F. Compound 6

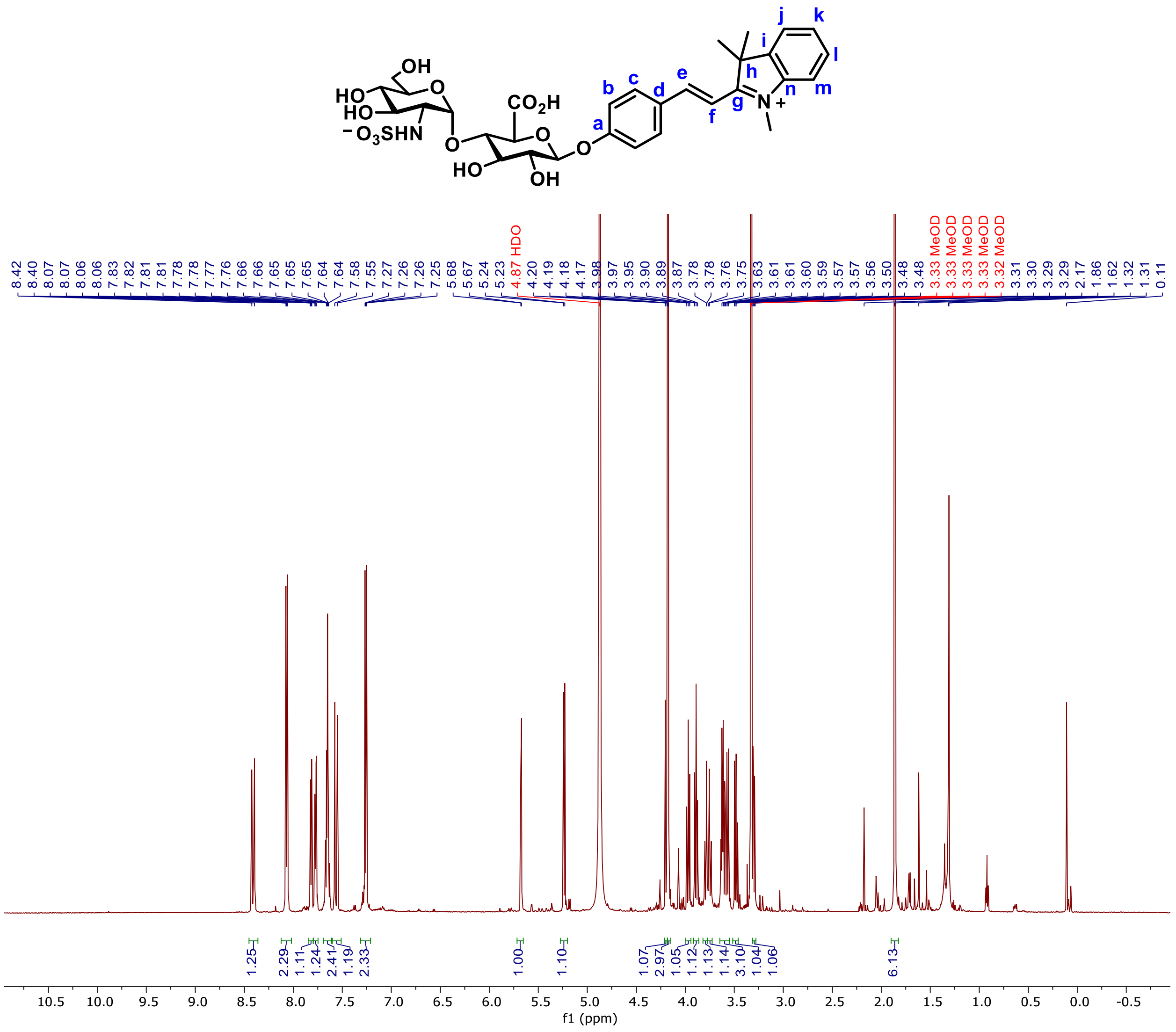

**Figure S39**. $^1$H NMR spectrum of compound **6** in MeOD

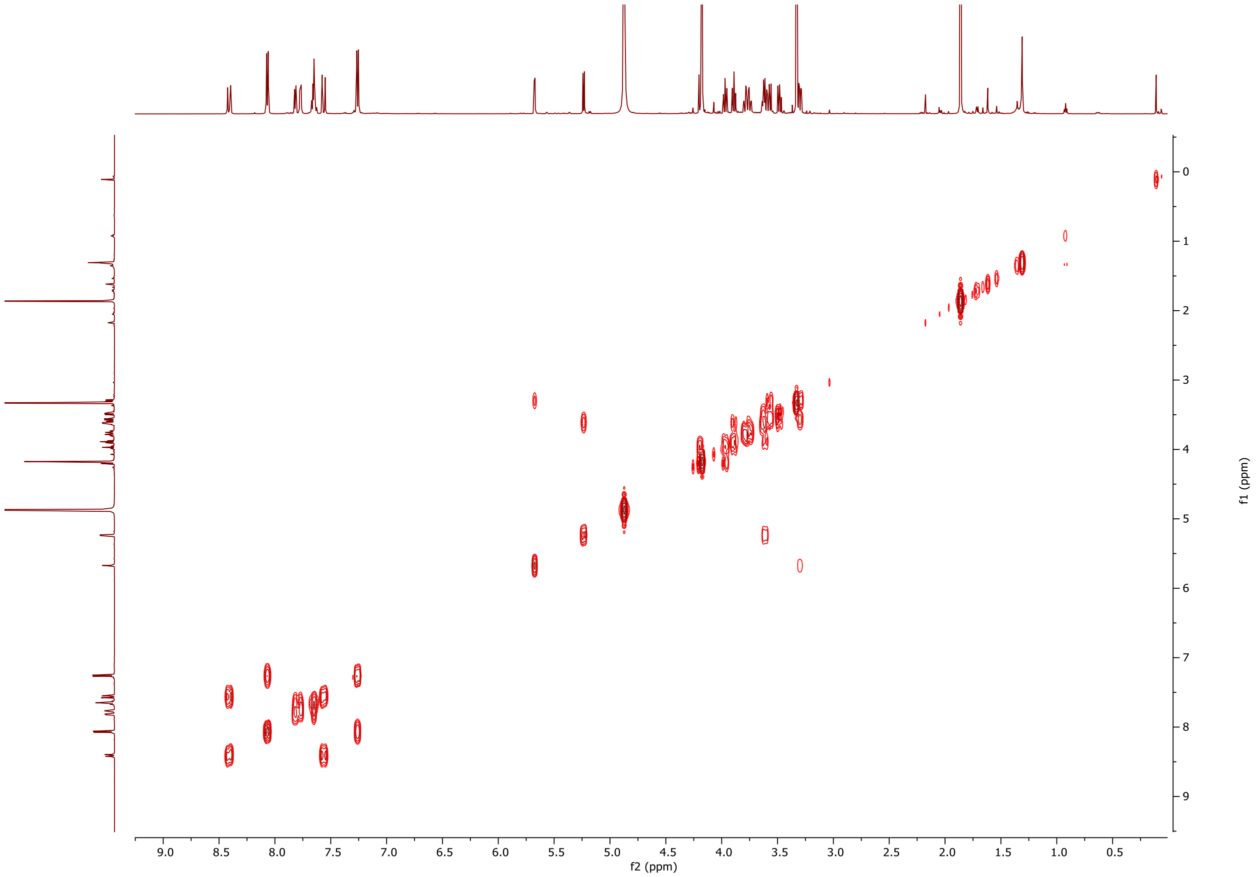

**Figure S40**. $^{1}H$-$^{1}H$ COSY NMR spectrum of compound **6** in MeOD

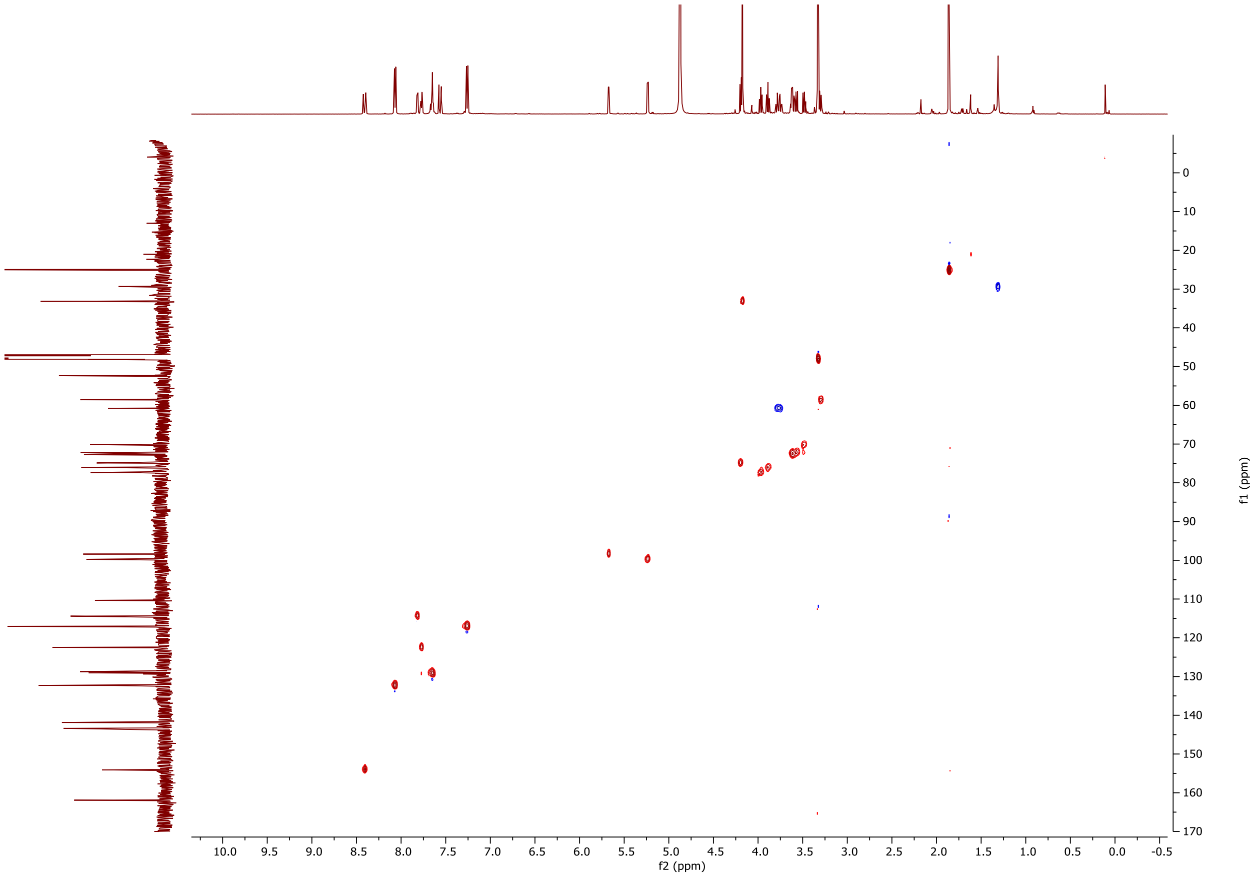

**Figure S41**. $^{1}H$-$^{13}C$ HSQC NMR spectrum of compound **6** in MeOD

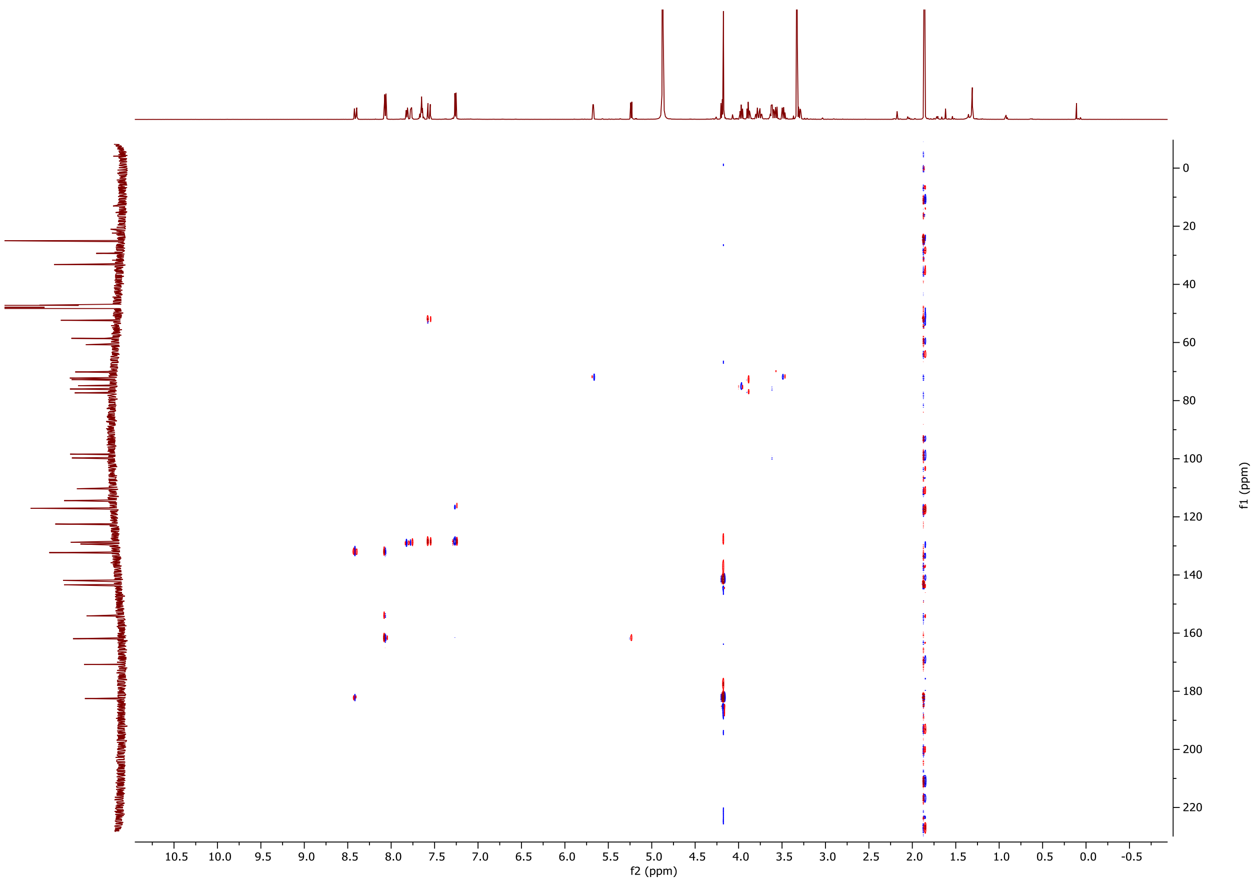

**Figure S42**. $^{1}H$-$^{13}C$ HMBC NMR spectrum of compound **6** in MeOD

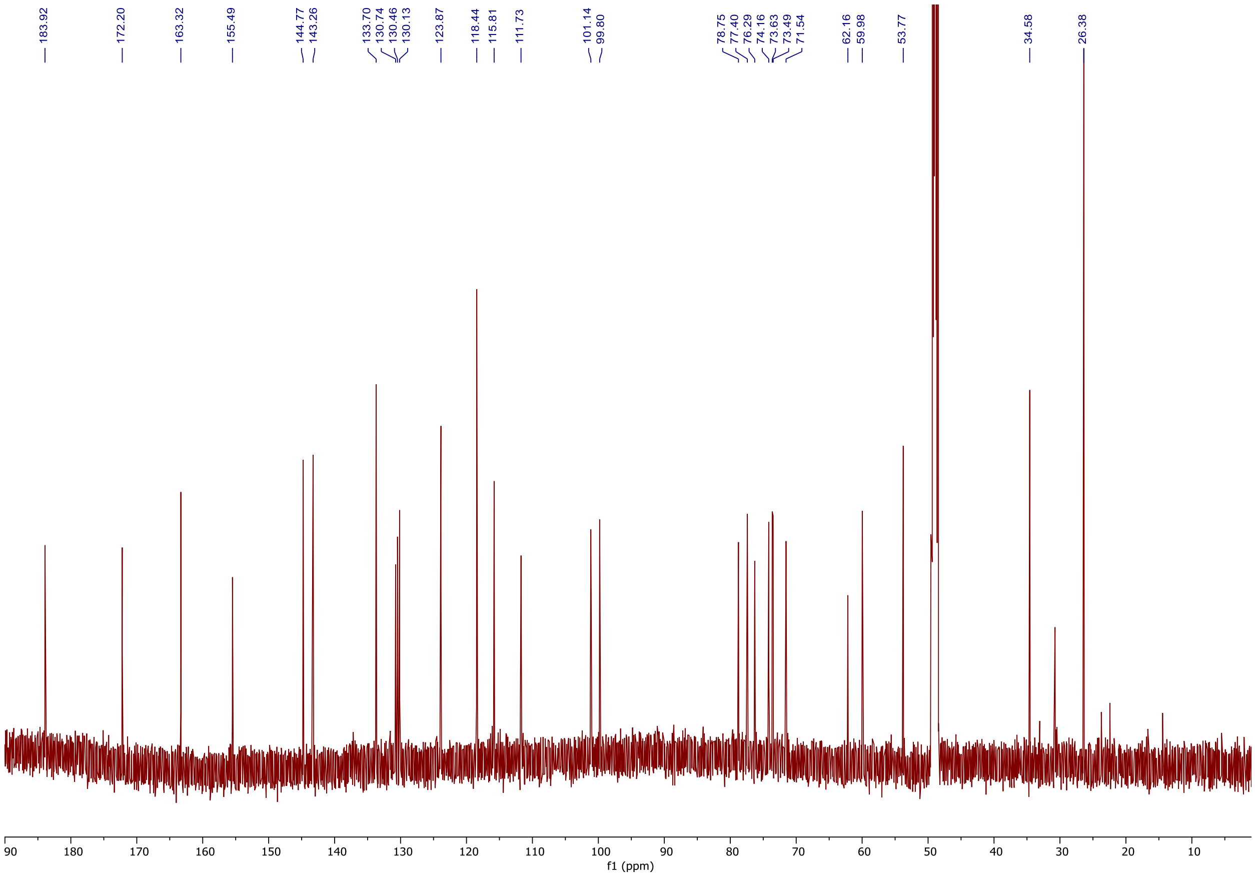

**Figure S43**. $^{13}C$ NMR spectrum of compound **6** in MeOD

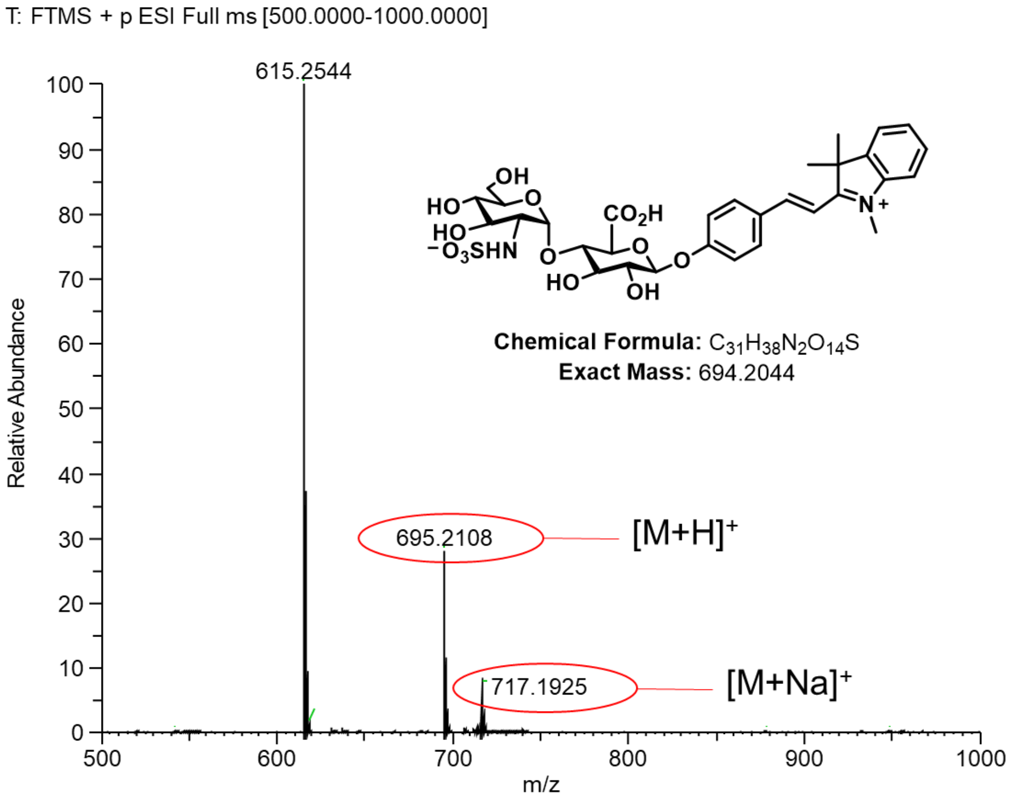

**Figure S44**. HRMS spectrum of compound **6**

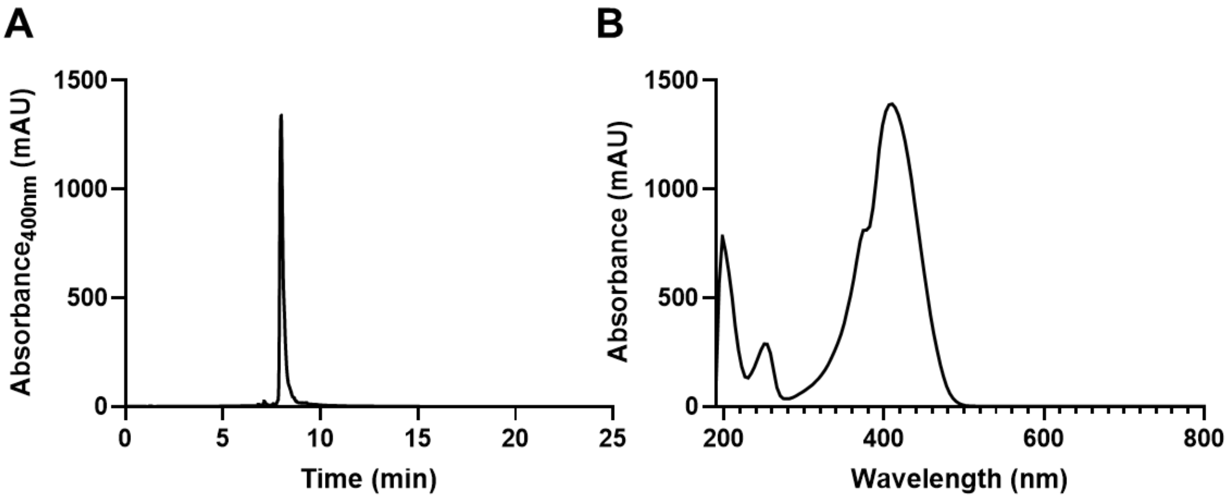

**Figure S45**. (**A**) HPLC trace of compound **6** (retention time, 7.98 min). (**B**) UV-VIS spectrum ($\lambda_{max}$, 408 nm) of compound **6.** HPLC Method: 2-100%, 0-20 min; 100%, 20-22.5 min; MeOH/ $H_2O$ with 0.1% Formic Acid.

### G. Compound 7

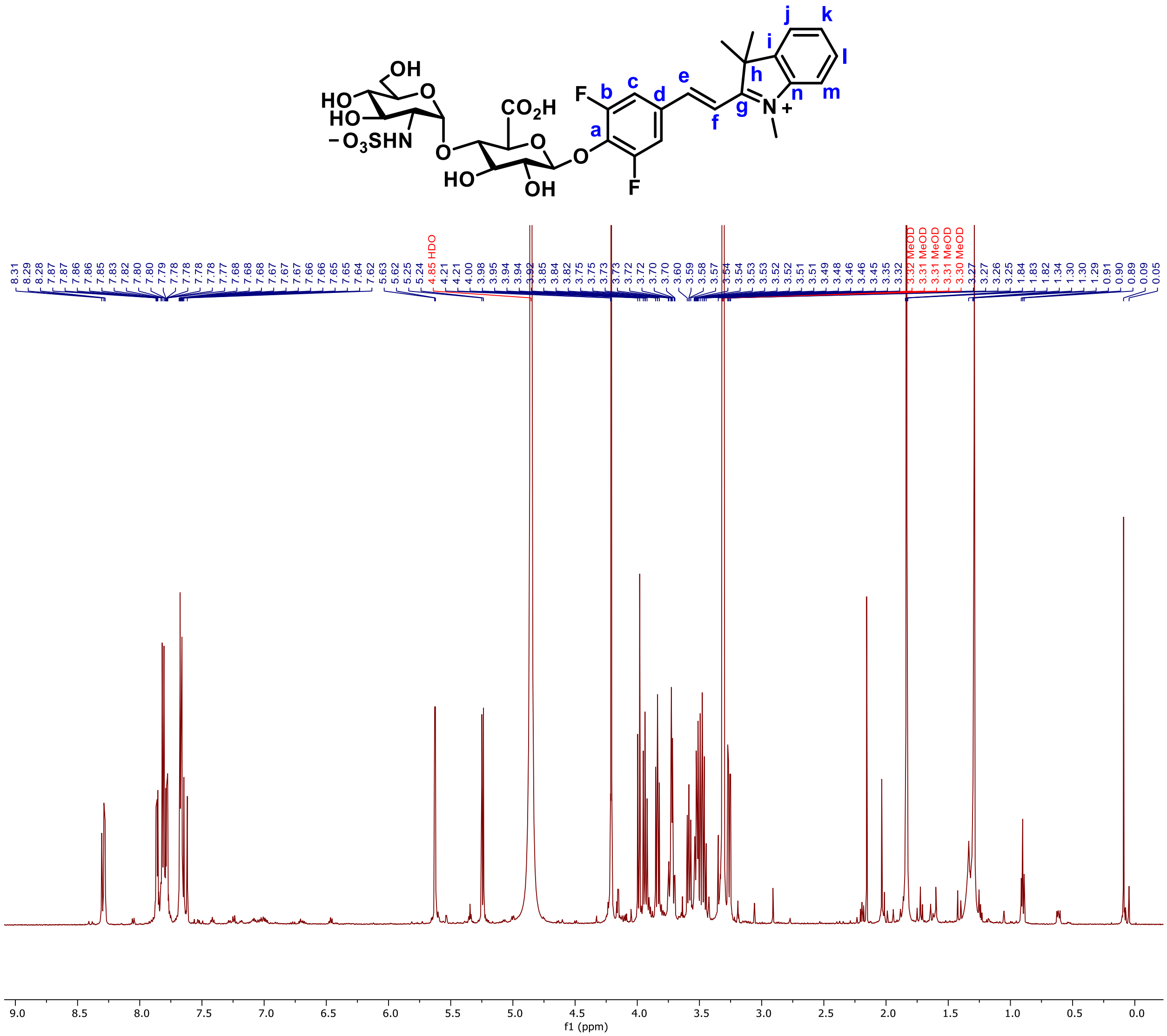

**Figure S46.** $^{1}H$ NMR spectrum of compound **7** in MeOD

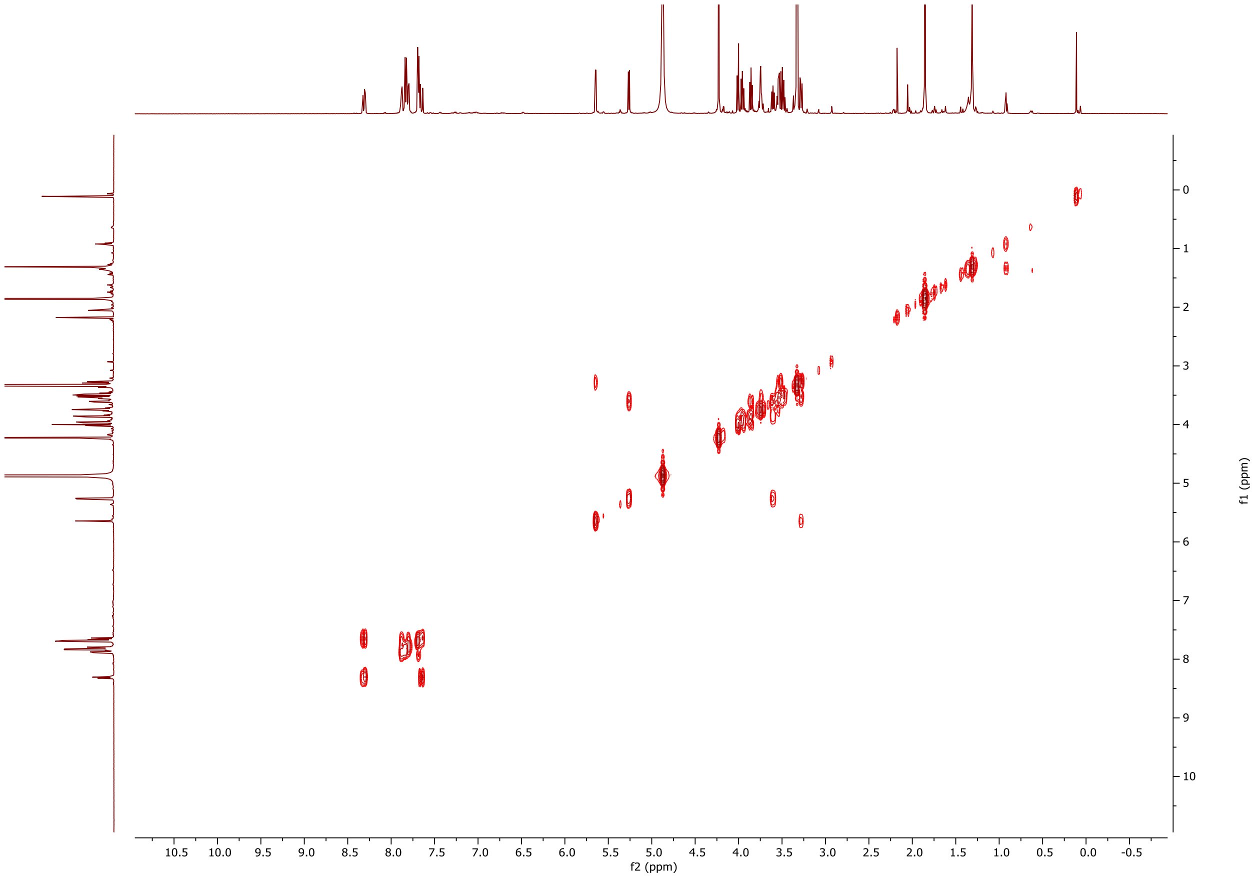

**Figure S47**. $^{1}$H-$^{1}$H COSY NMR spectrum of compound **7** in MeOD

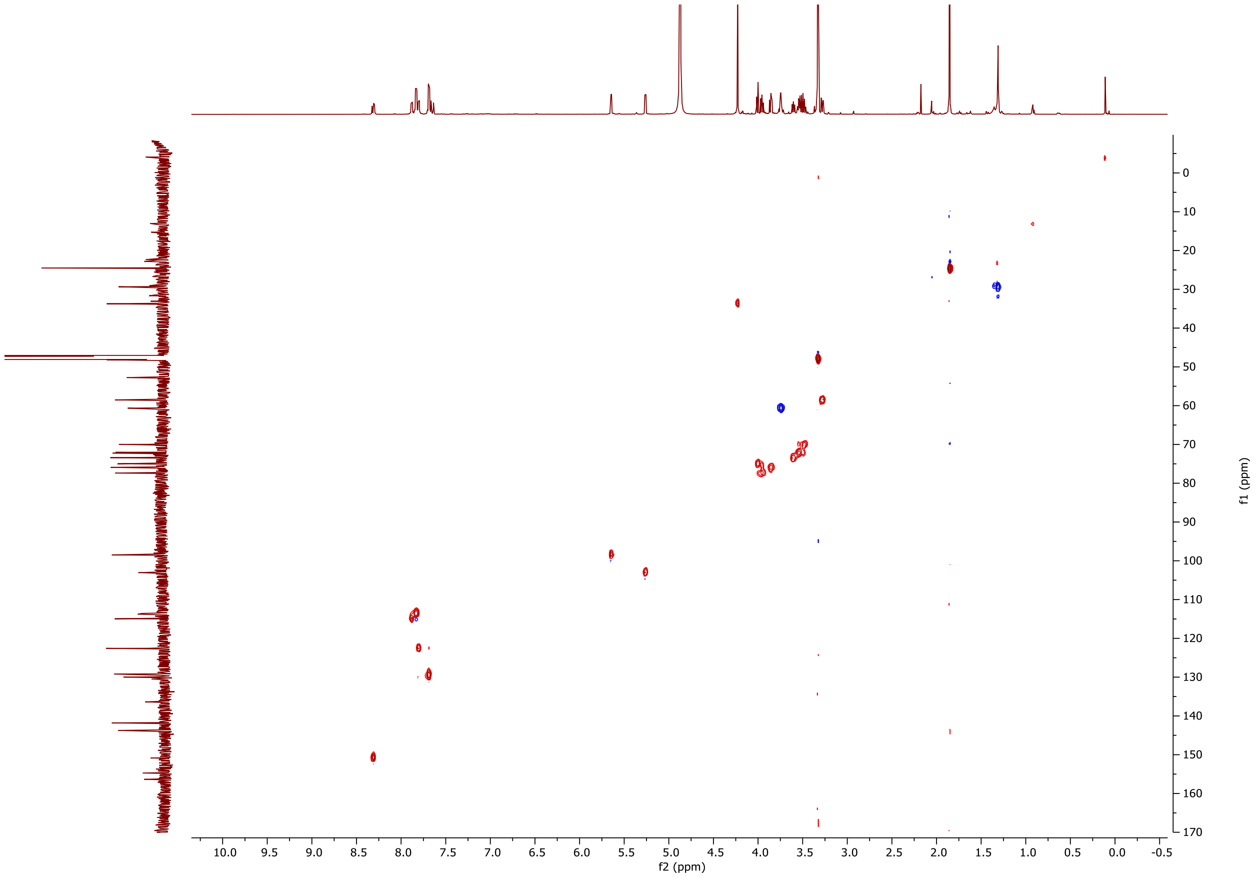

**Figure S48**. $^{1}H$-$^{13}C$ HSQC NMR spectrum of compound **7** in MeOD

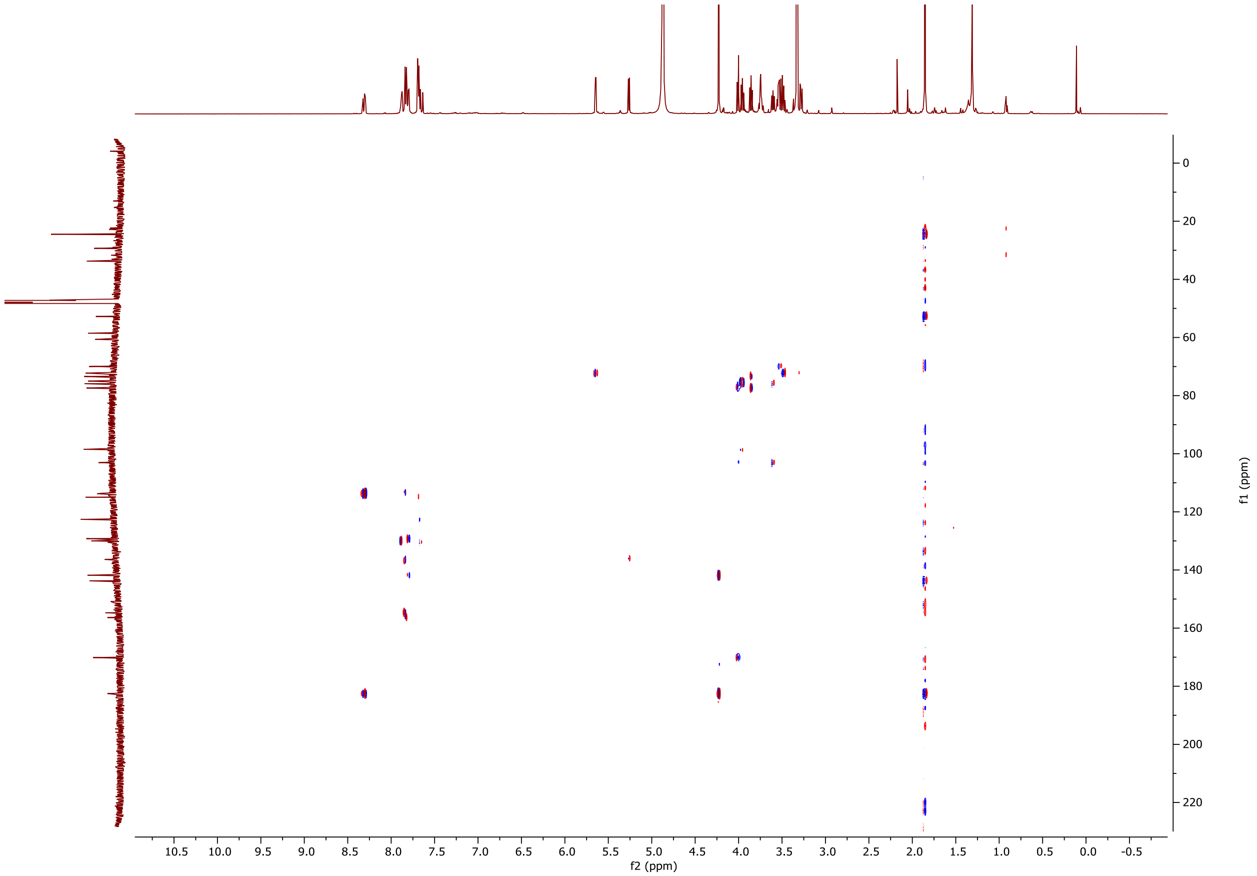

**Figure S49**. $^{1}H$-$^{13}C$ HMBC NMR spectrum of compound **7** in MeOD

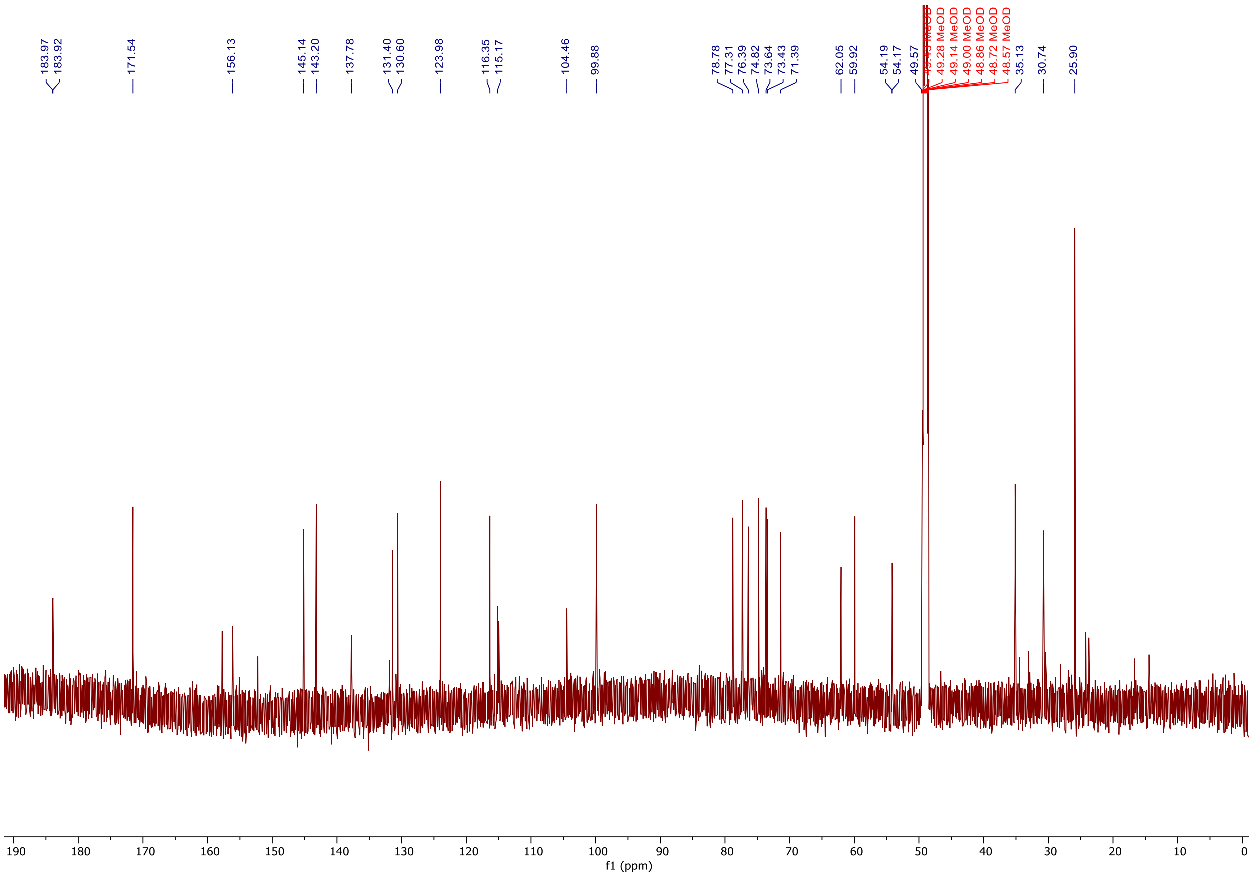

**Figure S50**. $^{13}$C NMR spectrum of compound **7** in MeOD

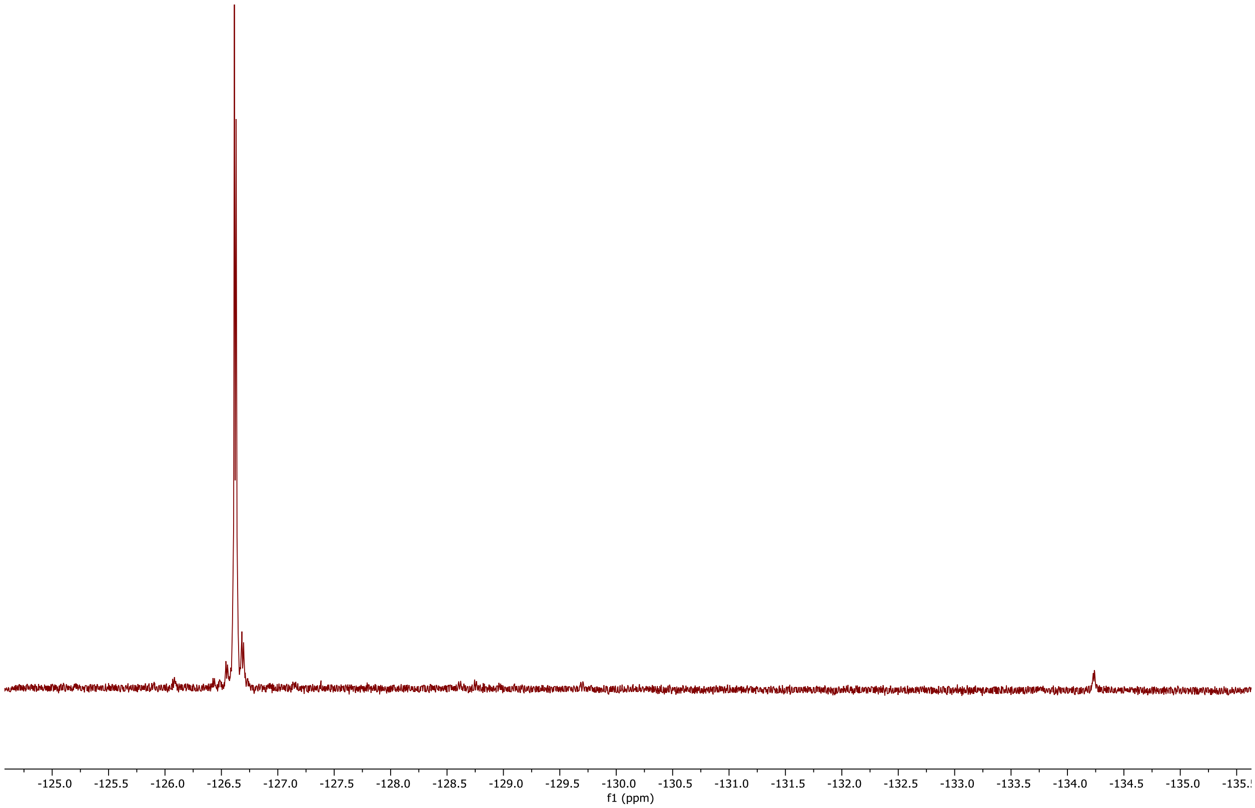

**Figure S51.** $^{19}F$ NMR spectrum of compound **7** in MeOD

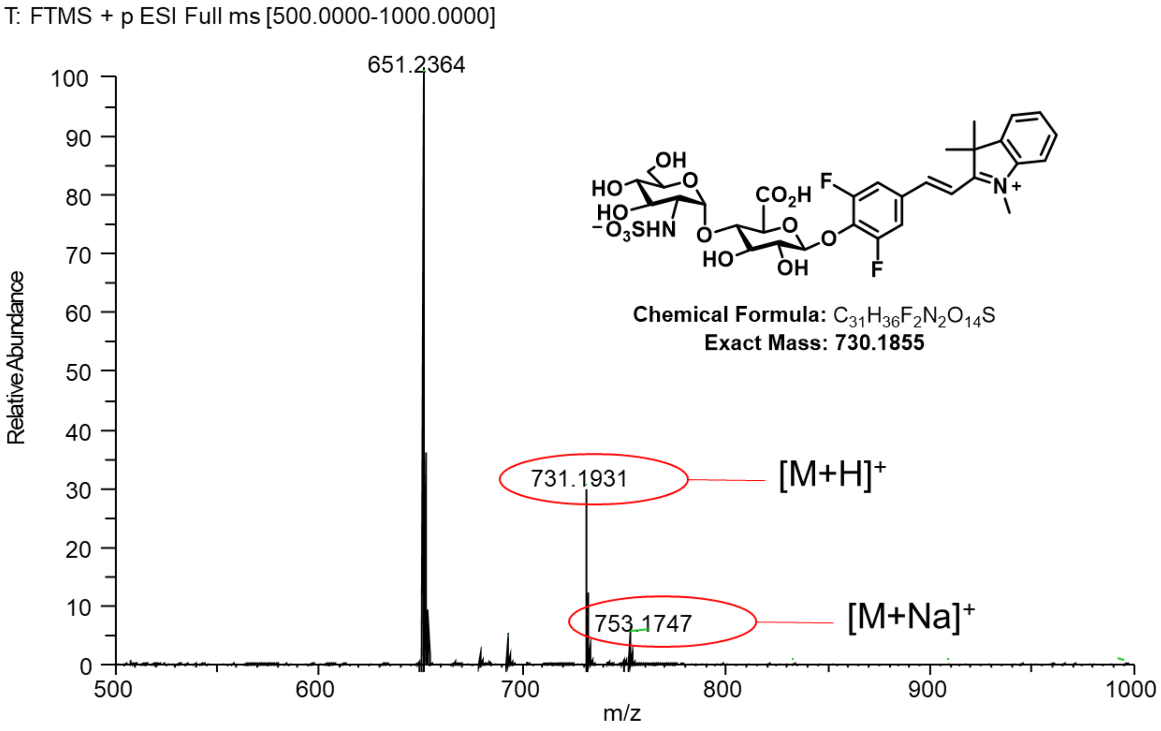

**Figure S52**. HRMS spectrum of compound **7**

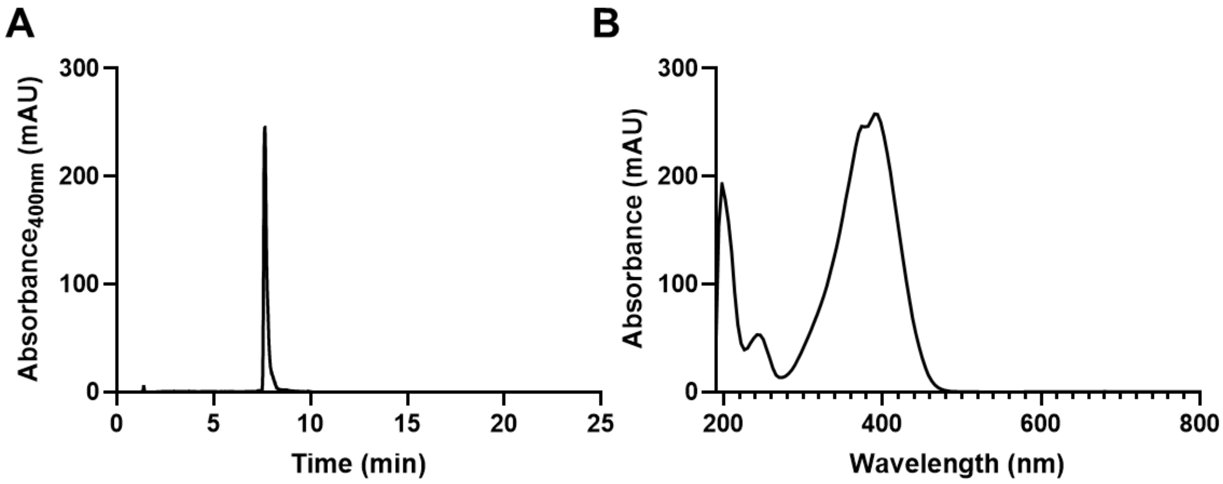

**Figure S53**. (**A**) HPLC trace of compound **7** (retention time, 7.64 min). (**B**) UV-VIS spectrum ($\lambda_{max}$, 392 nm) of compound **7.** HPLC Method: 2-100%, 0-20 min; 100%, 20-22.5 min; MeOH/$H_2O$ with 0.1% Formic Acid.
